# Microbial genomic trait evolution is dominated by frequent and rare pulsed evolution

**DOI:** 10.1101/2021.04.19.440498

**Authors:** Yingnan Gao, Martin Wu

## Abstract

On the macroevolutionary timescale, does trait evolution proceed gradually or by rapid bursts (pulses) separated by prolonged periods of stasis or slow evolution? Although studies have shown pulsed evolution is prevalent in animals, our knowledge about the tempo and mode of evolution across the tree of life is very limited. This long-standing debate calls for a test in bacteria and archaea, the most ancient and diverse forms of life with unique population genetic properties (asexual reproduction, large population sizes, short generation times, high dispersal rates and extensive lateral gene transfers). Using a likelihood-based framework, we analyzed evolutionary patterns of four microbial genomic traits (genome size, genome GC%, 16S rRNA GC% and the nitrogen use in proteins) on a broad macroevolutionary timescale. Our model fitting of phylogenetic comparative data shows that pulsed evolution is not only present, but also prevalent and predominant in microbial genomic trait evolution. Interestingly, for the first time, we detected two distinct types of pulsed evolution (small frequent and large rare jumps) that are predicted by the punctuated equilibrium and quantum evolution theories. Our findings suggest that major bacterial lineages could have originated in quick bursts and pulsed evolution is a common theme across the tree of life despite the drastically different population genetic properties of bacteria, archaea and eukaryotes.

## Introduction

There has been a long-standing debate about the tempo and mode of trait evolution on the macroevolutionary timescale. The gradualism theory states that evolution occurs gradually by small changes that accumulate over a long period of time (1). The pulsed evolution theory, on the other hand, argues that evolution mostly proceeds in bursts of larger changes (jumps) separated by long periods of stasis or slow evolution (1–3). Two types of jumps have been proposed in pulsed evolution. Simpson’s quantum evolution theory postulates that jumps happen when lineages shift into new adaptive zones and these jumps play an important role in the origination of higher taxa (2), while Eldredge and Gould’s later punctuated equilibrium theory focuses exclusively on jumps associated with speciation (1). Conceptually, these two types of jumps exist side-by-side but differ in their frequencies and magnitudes. Studies of animal fossil records support the punctuated equilibrium theory (4, 5), and more recent phylogenetic comparative studies of vertebrate body size (6–8) also provide evidence for quantum evolution. Together, they show that evolution is not solely composed of slow and gradual changes but also instant jumps on the macroevolution timescale.

Analogous studies in bacteria and archaea, the most ancient and diverse forms of life on Earth, are lacking, largely due to the scarcity of fossil records and well-measured quantitative phenotypic traits in microbes. Fortunately, the phenotypic evolution of microbial species can be reconstructed from extant genome sequences. Several genomic features are highly correlated with the microbial life strategy. For example, the GC% of the ribosomal RNA gene is correlated with the optimal growth temperature of bacteria and archaea (9). According to the genome streamlining theory, genome size, genomic GC% and the nitrogen use in proteins all evolve in response to changes of nutrient level in the environment (10, 11). These genomic features can be accurately determined from thousands of complete genomes currently available that represent a broad range of closely and distantly related lineages, making it possible to study the tempo and mode of trait evolution in microbes over a broad spectrum of macroevolution timescales.

Interestingly, long-term experimental evolution has shown evidence of pulsed evolution in *Escherichia coli* cell size (12). However, on the macroevolutionary timescale, the role of pulsed evolution in microbial trait evolution remains largely unknown. Although it is well known that there are large trait changes between bacterial clades (e.g., the genomic GC% of high GC vs low GC Gram-positives, AT-rich obligate intracellular bacteria vs their free-living relatives)(13, 14), it is unclear whether these large trait changes arose gradually or rapidly by jumps during the time the clades diverged from each other. Compared to animals and plants, bacteria and archaea reproduce asexually and have relatively large population sizes, high dispersal rates and short generation times. Another salient feature unique to microbes is that their genomes can often leap by large scale horizontal gene transfers (15), which obviously will have an impact on the tempo and mode of evolution. Given these unique features, a central question is whether the tempo and mode of microbial trait evolution are similar to those of eukaryotes and whether pulsed evolution is a universal theme across the tree of life, and if so, to what extent pulsed evolution contributes to microbial trait evolution?

## Results

### Gradual evolution does not explain microbial genomic trait evolution

We downloaded 10,616 and 263 complete bacterial and archaeal genomes respectively from the NCBI RefSeq database, from which we reconstructed genome trees and selected 6,668 and 247 representative genomes that passed quality control (Methods). For each representative genome, we calculated four genomic traits (genomic GC%, rRNA GC%, genome size and the average nitrogen atoms per residual side chain N-ARSC), all of which showing strong phylogenetic signals (Pagel’s *λ*>0.99). Using phylogenetically independent contrast (PIC) of tip-pairs, we found that although the four genomic traits are significantly correlated as previously reported (10, 16), the correlation appears to be weak (Fig. S1), with the proportion of variance explained by the other traits being less than 13.5% for all four traits. Therefore, to capture the possible variation in the tempo and mode of evolution, we chose to test each of the four traits separately. Notably, the PIC distributions of these traits in bacteria drastically deviate from the normal distribution expected by the Brownian motion (BM) model of gradual evolution (Fig. 1 A-D, two-sided Kolmogorov– Smirnov test, P<0.001 for all four traits). Specifically, all PIC distributions exhibit a strong leptokurtic (heavy-tailed) pattern with a positive excess kurtosis ranging from 5.79 to 13.47, indicating that extremely rapid large trait changes (pulses) occur more frequently than expected by the BM model. For archaea, the PIC distribution also deviates from the normal expectation (Fig. S2 A-D, two-sided Kolmogorov-Smirnov test, P<0.001, P=0.024, P=0.018 and P=0.155 for rRNA GC%, genomic GC%, genome size and N-ARSC, respectively), with the excess kurtosis ranging from 1.47 to 7.58. Although inconsistent with BM, such a heavy-tailed pattern can be explained by pulsed evolution. Extremely rapid large trait changes (|PIC|>3) take place more frequently than expected by the normal distribution (0.27%) throughout the bacterial evolutionary history (Fig. S3), suggesting repeated episodes of pulsed trait evolution.

**Figure 1.**
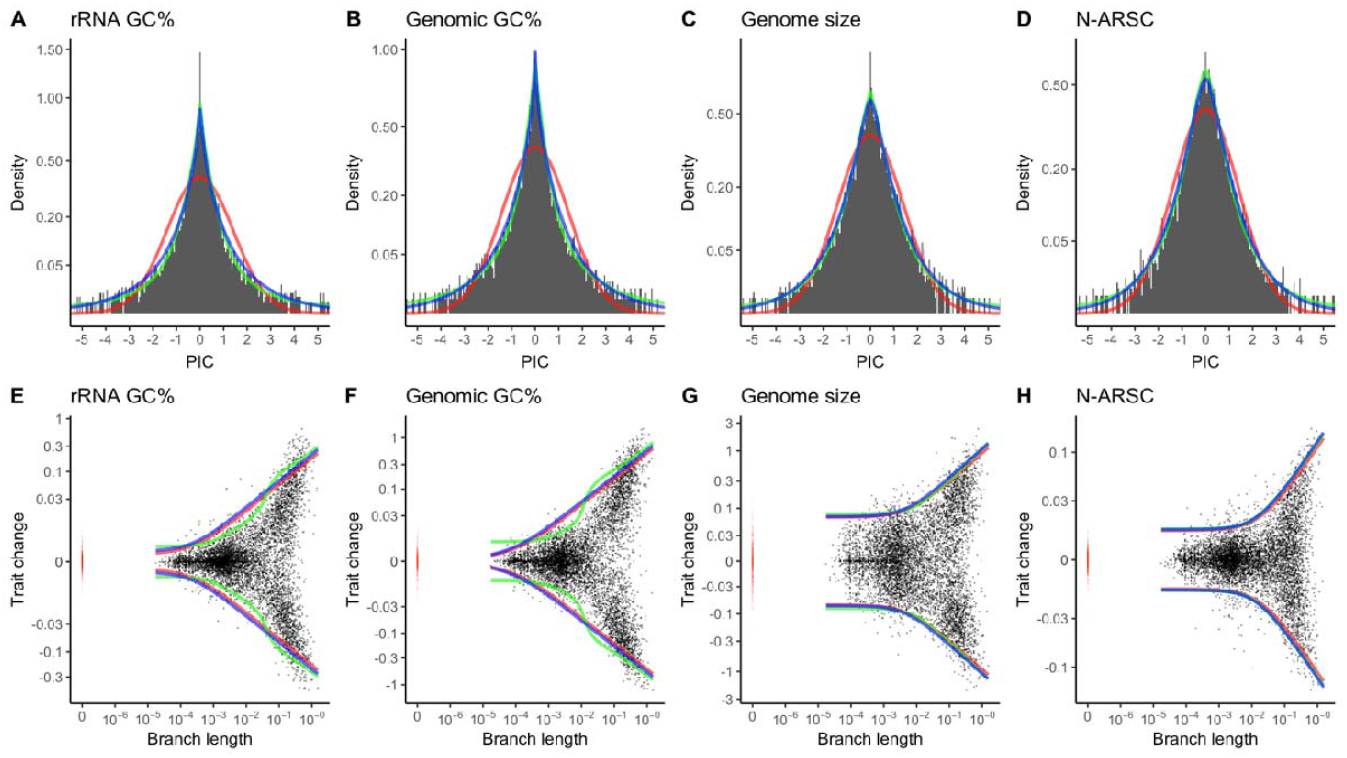
Pulsed evolution models fit bacterial trait evolution better than the BM model and the variable rate model. (A-D) PIC distributions (black bars) deviate significantly from the normal distribution of the BM model (red line). The pulsed evolution models that include two or three Poisson processes (PE2 or PE3, green line) greatly improve the fit to the overall PIC distributions. The variable rate model also improves the fit to the overall PIC distribution. Square-root transformation is applied to the y-axis (density) to better show the deviation in the frequency of large PICs. (E-H) Patterns of bacterial trait changes at different branch lengths. Trait changes derived from the bacterial phylogeny are shown in black dots. Trait differences between genomes separated by zero branch length are shown in red dots. The expected 95% confidence intervals (CI) of the models are shown in colored lines (red for the BM model, green for the pulsed evolution model, and blue for the variable rate model). Pseudo-log transformation is applied to the y-axis (trait change) to better show the trend of trait change in short branches.

### Modeling microbial genomic trait evolution

When plotted against the branch length, the trait changes between two sister nodes in the bacteria phylogeny display a “blunderbuss pattern” (Fig. 1 E-H). It starts with a period of stationary fluctuations where trait changes are bounded and the variance does not accumulate over time. Segmented linear regression analysis indicates that this phase of stasis lasts until the branch length reaches 0.001 substitutions/site for rRNA GC% (Fig. S4). On longer timescales, the stasis yields to a pattern of increasing divergence over time. The archaeal traits display similar patterns (Fig. S2 E-H). This “blunderbuss pattern”, first observed in the evolution of vertebrate body-size, is a signature of pulsed evolution (6). Interestingly, for rRNA GC%, we observed a second spike in the trait divergence rate at 0.025 substitutions/site (Fig. S4), indicating a change of evolution tempo at this point.

To formally test whether pulsed evolution explains the patterns, we model the trait change between two sister nodes using the Levy process (8). More specifically, we model the trait change as the sum of three independent stochastic variables: pulsed evolution, gradual evolution, and time-independent trait variation. We assume pulsed evolution occurs at a constant rate relative to the molecular divergence and the jump size follows a normal distribution with a mean of zero. As a result, the pulsed evolution is modeled as a compound Poisson process with normal jumps, with parameters *λ* and *σ*^2^ denoting the frequency (number of expected jumps per lineage per unit branch length) and the magnitude (variance of trait change) of the jumps, respectively. We model gradual evolution using the classic BM model with a single parameter 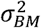 denoting the rate of the gradual trait change. Meanwhile, we observed trait variation between genomes separated by zero branch length, indicating the presence of time-independent variation in our phylogeny. Because this variation follows a leptokurtic distribution, we model the time-independent variation with the Laplace distribution with one single parameter ε denoting its variance for simplicity and convenience. It should be noted that a jump in the genomic trait may be coupled with an increase in the molecular divergence rate, especially for those traits affecting protein sequences (e.g., genomic GC% and N-ARSC). However, such correlation between the molecular branch length and the trait change will only reduce the signal of pulsed evolution, as the increased branch length provides greater power for gradual evolution to explain the trait variation (Fig. S5).

The changing tempos revealed by segmented linear regression suggests that one Poisson process may not adequately describe the patterns of pulsed evolution, prompting us to add multiple Poisson processes (with different jump magnitudes) to our modeling. Therefore, using the framework described above, we tested seven different models (Table 1). The BM model delineates gradual evolution with a constant rate while the VRG model describes gradual evolution with continuous variable rates. The PE1, PE2 and PE3 models describe pulsed evolution with one, two or three Poisson processes respectively. The PE(n)+BM models represent trait evolution with both pulsed and gradual evolution. Details of these models and the maximum likelihood framework are provided in the Supplementary Text. Using simulated data, we show that our maximum likelihood (ML) framework can distinguish the BM, continuous variable rate and pulsed evolution models (Table S1) and capture the frequency and magnitude of jumps (Table S2). Among pulsed evolution models, our ML framework can distinguish models with one Poisson process from those with more than one Poisson process, but it favors models with a BM component and tends to underestimate the contribution of pulsed evolution (Table S1). This is because it is difficult to distinguish between frequent jumps and gradual evolution on long branches (Fig. S5).

**Table 1.** The seven trait evolution models tested in this study.

| Models | Free parameters |
| --- | --- |
| Brownian motion (BM) | $\sigma_{BM}^2, \varepsilon$ |
| Variable rate model with Gamma distribution (VRG) | $p, k, \theta, \varepsilon$ |
| One Poisson process (PE1) | $\lambda_1, \sigma_1^2, \varepsilon$ |
| One Poisson process and Brownian motion (PE1+BM) | $\sigma_{BM}^2, \lambda_1, \sigma_1^2, \varepsilon$ |
| Two Poisson processes (PE2) | $\lambda_1, \sigma_1^2, \lambda_2, \sigma_2^2, \varepsilon$ |
| Two Poisson processes and Brownian motion (PE2+BM) | $\sigma_{BM}^2, \lambda_1, \sigma_1^2, \lambda_2, \sigma_2^2, \varepsilon$ |
| Three Poisson processes (PE3) | $\lambda_1, \sigma_1^2, \lambda_2, \sigma_2^2, \lambda_3, \sigma_3^2, \varepsilon$ |

### Microbial trait evolution is dominated by frequent and rare jumps

For the four traits we have examined in bacteria and archaea, the best model is always one with a pulsed evolution component. The relative support for the gradual evolution models is marginal, with AIC weights for the BM model <0.5% and those for the VRG model <2.1% (Table 2). Both the pulsed evolution and VRG models fit the overall PIC distributions much better than the BM (Fig. 1 A-D and Fig. S2 A-D). The pulsed evolution model also fits the patterns of genomic GC% and 16S rRNA GC% changes at different branch lengths better than the BM and VRG models (Fig. 1 E, F and Fig. S2 E, F). The strong support by AIC and improved fit to the PIC pattern across different branch lengths suggest that pulsed evolution is present in both bacterial and archaeal genomic traits. To test the prevalence of pulsed evolution in bacteria, we separately fitted our models in 17 bacterial families that each contained at least 100 genomes. We found that trait evolution in 76.5%, 88.2%, 52.9% and 23.5% of tested families were best explained by a model with a pulsed evolution component (PE1, PE1+BM, PE2, PE2+BM or PE3) for rRNA GC%, genomic GC%, genome size and N-ARSC respectively, indicating that pulsed evolution is prevalent in bacteria (Table S3). To examine the effect of sample size on the power of detecting pulsed evolution, we simulated trait evolution under the pulsed evolution model and conducted model selection on the simulated data. Our simulation shows that when the number of genomes decreases, the power to detect pulsed evolution also decreases (Table S4), suggesting that we might have underestimated the prevalence of pulsed evolution in the 17 bacterial families. We did not test the prevalence of pulsed evolution in archaea because of the insufficient number of archaeal genomes at the lower taxonomic levels.

**Table 2.** AIC values for each model fitted for bacterial and archaeal trait evolution. The AIC values for the best model and models that are not significantly inferior (AIC change <2) in each trait are in bold.

| Domain | Trait | BM | VRG | PE1 | PE1+BM | PE2 | PE2+BM | PE3 |
| --- | --- | --- | --- | --- | --- | --- | --- | --- |
| Bacteria | Ribosomal RNA GC% | -38431 | -41519 | -40747 | -41171 | **-41591** | -41589 | -41587 |
|  | Genomic GC% | -28539 | -30432 | -30921 | -31171 | **-31299** | -31297 | -31295 |
|  | Genome size | -15572 | -15990 | -15929 | -16022 | -16075 | -16090 | **-16107** |
|  | N-ARSC | -41944 | -42092 | -42123 | -42214 | -42214 | **-42218** | **-42217** |
| Archaea | Ribosomal RNA GC% | -483 | -601 | -606 | -636 | **-642** | -640 | -638 |
|  | Genomic GC% | -440 | -469 | -488 | **-503** | **-504** | -502 | -500 |
|  | Genome size | -348 | -351 | **-359** | -357 | -355 | -353 | -351 |
|  | N-ARSC | -1202 | -1205 | **-1212** | **-1211** | -1210 | -1208 | -1206 |

Using parameters of the best models (Table 3), we estimated the relative contribution of each compound Poisson process. The variable ε represents the trait variance in the initial stasis phase (Fig. 1). Its estimated value approximates the intraspecific trait variation between genomes with identical marker gene alignments (i.e., zero branch length) and therefore is used as the baseline. The jumps vary greatly in their frequencies and magnitudes, but can be roughly classified into two types: small and frequent, or large and rare (Table 3). For example, for rRNA GC%, rare jumps (1.96 jumps per lineage per unit branch length) are extremely large in magnitude, as the standard deviation of trait change introduced by one rare jump is approximately 60 times of 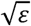, or roughly equivalent to that introduced by 700 million years (0.35 substitutions/site) of gradual evolution under the BM model, and approximately corresponds to 5.8 °C change in the optimal growth temperature. In comparison, the frequent jumps (118 jumps per lineage per unit branch length) are 60 times more frequent but their sizes are only about 3 times of 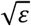. Overall, rare jumps predominate in trait evolution as they contribute more than 74% of variation in each trait over the whole phylogeny. Similarly, pulsed evolution also predominates in archaea as the PE1 and PE2 models are the best models in all archaeal traits (Table 2). Due to the limited number of archaeal genomes, we cannot robustly estimate the parameters of each jump process in archaea.

**Table 3.**
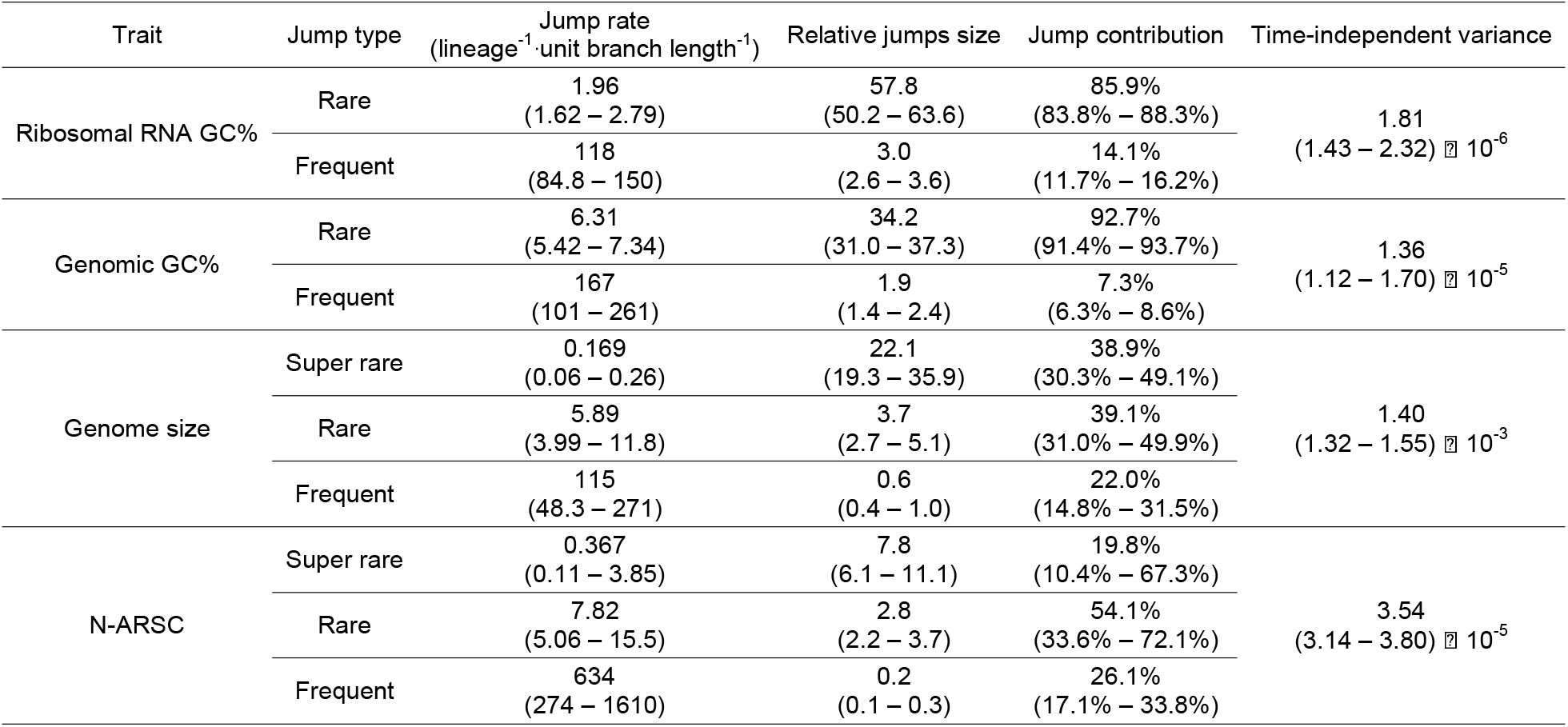
Model statistics of pulsed evolution in different bacterial traits. The 95% confidence interval for each model statistic is listed in the parentheses after the statistic.

| Trait | Jump type | Jump rate<br>(lineage<sup>-1</sup>·unit branch length<sup>-1</sup>) | Relative jumps size | Jump contribution | Time-independent variance |
| --- | --- | --- | --- | --- | --- |
| Ribosomal RNA GC% | Rare | 1.96<br>(1.62 – 2.79) | 57.8<br>(50.2 – 63.6) | 85.9%<br>(83.8% – 88.3%) | 1.81<br>(1.43 – 2.32) × 10<sup>-6</sup> |
|  | Frequent | 118<br>(84.8 – 150) | 3.0<br>(2.6 – 3.6) | 14.1%<br>(11.7% – 16.2%) |  |
| Genomic GC% | Rare | 6.31<br>(5.42 – 7.34) | 34.2<br>(31.0 – 37.3) | 92.7%<br>(91.4% – 93.7%) | 1.36<br>(1.12 – 1.70) × 10<sup>-5</sup> |
|  | Frequent | 167<br>(101 – 261) | 1.9<br>(1.4 – 2.4) | 7.3%<br>(6.3% – 8.6%) |  |
| Genome size | Super rare | 0.169<br>(0.06 – 0.26) | 22.1<br>(19.3 – 35.9) | 38.9%<br>(30.3% – 49.1%) | 1.40<br>(1.32 – 1.55) × 10<sup>-3</sup> |
|  | Rare | 5.89<br>(3.99 – 11.8) | 3.7<br>(2.7 – 5.1) | 39.1%<br>(31.0% – 49.9%) |  |
|  | Frequent | 115<br>(48.3 – 271) | 0.6<br>(0.4 – 1.0) | 22.0%<br>(14.8% – 31.5%) |  |
| N-ARSC | Super rare | 0.367<br>(0.11 – 3.85) | 7.8<br>(6.1 – 11.1) | 19.8%<br>(10.4% – 67.3%) | 3.54<br>(3.14 – 3.80) × 10<sup>-5</sup> |
|  | Rare | 7.82<br>(5.06 – 15.5) | 2.8<br>(2.2 – 3.7) | 54.1%<br>(33.6% – 72.1%) |  |
|  | Frequent | 634<br>(274 – 1610) | 0.2<br>(0.1 – 0.3) | 26.1%<br>(17.1% – 33.8%) |  |

To evaluate the effect of the tree topology on our model fitting, we fitted models on the genome tree of the family Enterobacteriaceae (with 748 genomes) made using either FastTree or RAxML. We found that the fitted model parameters are highly similar using these two trees (Table S5). We also validated our fitted model with the bacterial phylogeny downloaded from Genome Taxonomy Database (GTDB)(17), and found that pulsed evolution models with at least two Poisson processes are also the best models (Table S6). Because all the models tested in this study are time-reversible, rooting of the phylogeny at different points does not affect the results.

### Rare jumps are correlated with cladogenesis in bacteria

We use simulation to test whether we can detect specific jumps along the phylogeny using the posterior probability estimated under pulsed evolution models. Because pulsed evolution models with at least two Poisson processes are the best models for bacterial genomic trait evolution, we simulated trait evolution under the PE2 model. We found that posterior probability calculated under the pulsed evolution model can predict both frequent and rare jumps very well, with the receiver operating characteristic (ROC) curves having an average area under curve (AUC) of 0.97 for both frequent and rare jumps (Fig. S6). Using a posterior probability cutoff of 0.9, we can achieve a specificity of 0.99 for detecting both frequent and rare jumps. Additionally, we can detect rare jumps on short branches (defined as branches with a prior probability of having at least one jump <0.1) with a sensitivity of 0.6. As a comparison, we also tested the performance of BayesTraits, which assumes the continuous variable rate model VRG. BayesTraits failed to accurately capture the frequent and rare jumps, yielding an average AUC below 0.70 (Fig. S6). This shows that the continuous variable rate model is inadequate to capture the mode of pulsed evolution.

Using a posterior probability cutoff of 0.9, we mapped the rare jumps of the genomic traits onto the bacterial phylogeny. We found that jumps occurred throughout the phylogeny (Fig. 2.), again indicating that pulsed evolution is prevalent in bacterial evolution history. We detect recent rare jumps in genome size that happen on short branches separating recently diverged species. These jumps correspond to events of very recent genome reduction and expansion (Table S7). We also detect rare jumps in more ancient branches. Some of the rare jumps are associated with known key evolutionary adaptations, validating our predictions. For instance, A classic example of adaptation to endosymbiosis occurred within the family *Enterobacteriaceae*, in the lineage leading to a clade of insect endosymbionts that includes the genera *Buchnera, Wigglesworthia* and *Candidatus Blochmannia* (18). Our model detects large rare jumps at the base of the clade in the genome size and genomic GC% (posterior probability > 0.9, Fig. 2). The recently described order Candidatus Nanopelagicales within the phylum Actinobacteria makes up the most abundant free-living bacteria in freshwater. Nanopelagicales have adapted to live in the nutrient poor environment by streamlining their genomes (19). Compared to its high GC Gram-positive sister clade, Nanopelagicales has dramatically reduced genome size (∼1.4 Mbp) and genomic GC% (∼48%). We detect large rare jumps in all genomic traits at the base of the order with extremely high confidence (posterior probability >0.99), suggesting that the genomic streamlining process happened not gradually but by jumps. Similar patterns have also been observed in the branch leading to the most abundant free-living marine bacteria Pelagibacterales and the intracellular bacteria Rickettsiales and Holosporales. Our model also predicts large rare jumps at higher taxonomic levels such as those at the base branch leading to the α-, β-, γ- and δ-proteobacteria (posterior probability >0.96) and the branch that separates the γ-proteobacteria from the rest of the proteobacteria (posterior probability >0.99). Our results suggest that these key evolutionary adaptations evolved in rapid bursts instead of through slow divergence of species over prolonged periods of time as proposed by the gradual evolution model.

**Figure 2.**
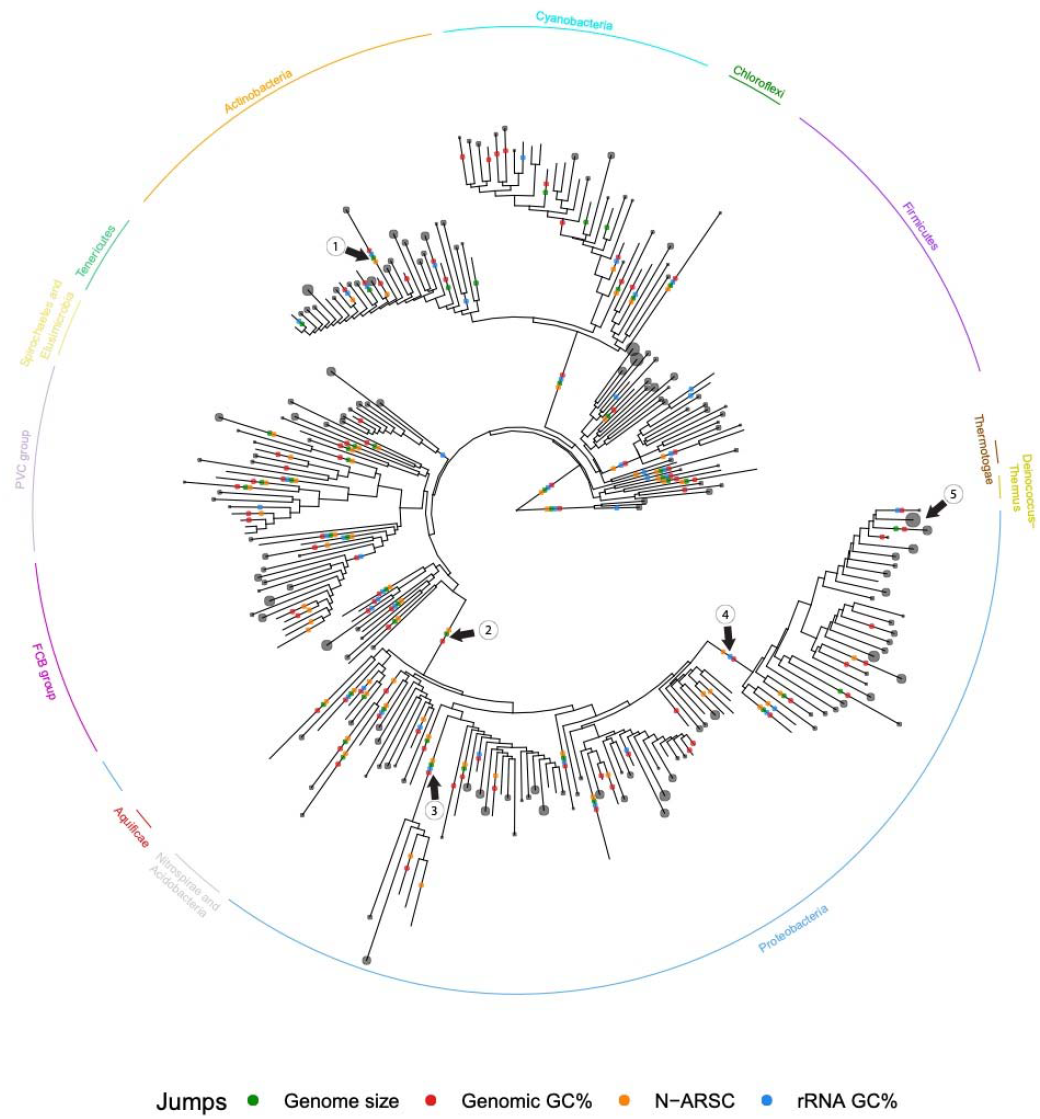
Rare jumps are widely distributed throughout the bacterial phylogeny. For clarity, clades have been collapsed at the taxonomic rank order and therefore the vast majority of the short branches in the tree are not shown in the figure. A collapsed order is represented by a grey circle at the tip whose diameter represents the number of genomes in the order. Colored dots are placed on branches where the posterior probability of having at least one rare or super rare jump event is greater than 0.9. Arrows point to branches leading to 1. the order Candidatus Nanopelagicales. 2. the α-, β-, γ- and δ-proteobacteria. 3. the orders Pelagibacterales, Rickettsiales and Holosporales. 4. γ-proteobacteria. 5. the genera *Buchnera, Wigglesworthia* and *Candidatus Blochmannia* within the family *Enterobacteriaceae*.

Next, we tested whether jumps are correlated with cladogenesis in bacteria by comparing the predicted frequency of jumps to the expected frequency for which we assume no correlation of jumps with cladogenesis (the null hypothesis). For example, if jumps happen significantly more frequently between two congener sister nodes than expected, jumps are considered correlated with the speciation event (cladogenesis at the species level). For frequent jumps, simulations indicated that we lacked the statistical power to reject the null hypothesis at every taxonomic level, and therefore we excluded them from this analysis. For rare and super rare jumps, we tested their correlation with cladogenesis from the species to order levels. We found that rare and super rare jumps occur more frequently than expected for all traits at the genus, family and order levels, except for N-ARSC at the genus level (Table 4). This increase in frequency is significant for rRNA GC%, genomic GC% and genome size at the genus and family levels (P≤0.050), and for rRNA and genomic GC% at the order level (P<0.001). Interestingly, we found that rare and super rare jumps happen less frequently than expected for all traits at the species level, although it is significant only for ribosomal GC% (P<0.001). Our results suggest that rare and super rare jumps are correlated with cladogenesis at higher taxonomic ranks.

**Table 4.** Differences in the percentage of contrasts with at least one rare or super rare jump between those inferred from the empirical data and the expectation from the null hypothesis. Significant differences are marked with asterisks. P-values and power (□) are listed in parentheses.

| Trait | Species | Genus | Family | Order |
| --- | --- | --- | --- | --- |
| Ribosomal GC% | -2.3%*<br>(P<0.001,<br>□>0.999) | +9.5%*<br>(P<0.001,<br>□>0.999) | +17.1%*<br>(P<0.001,<br>□=0.985) | +21.3%*<br>(P<0.001,<br>□=0.870) |
| Genomic GC% | -2.1%<br>(P=0.060,<br>□>0.999) | +8.1%*<br>(P<0.001,<br>□=0.995) | +8.9%*<br>(P<0.001,<br>□=0.610) | +11.5%*<br>(P<0.001,<br>□=0.205) |
| Genome size | -1.2%<br>(P=0.150,<br>□>0.999) | +3.5%*<br>(P=0.050,<br>□>0.999) | +4.9%*<br>(P<0.001,<br>□>0.999) | +4.1%<br>(P=0.075,<br>□>0.999) |
| N-ARSC | -0.2%<br>(P=0.630,<br>□>0.999) | -1.8%<br>(P=0.240,<br>□>0.999) | +4.1%<br>(P=0.06,<br>□=0.990) | +3.3%<br>(P=0.290,<br>□=0.770) |

## Discussion

The central question we try to address in this study is the tempo and mode of microbial trait evolution: whether the traits evolve mainly by gradual or pulsed evolution. Large trait differences between bacterial lineages are well known (13, 14), but it is less clear whether these large trait changes arose gradually over time or rapidly by jumps (the mode). Using a maximum likelihood framework, we explicitly test the mode of genomic trait evolution in bacteria and archaea and we show that pulsed evolution explains the patterns significantly better than both the constant and variable rate gradual evolution models. Our analysis suggests that pulsed evolution is not only present, but also prevalent and dominant in microbial genomic trait evolution.

Microbes are known for rapid evolution. Why are these genomic traits constrained for millions of years before they diverge? The stasis at the species level can be explained by stabilizing selection that eliminates variants falling outside of a stable niche (20). Alternatively, it can be maintained by gene flow, as suggested by Futuyma’s ephemeral divergence theory (21). Futuyma proposes that novel adaptive trait variation arises frequently in local populations, but the spatial and temporal mosaic nature of niches prevents such local adaptations from spreading to the entire species because they are wiped out by the gene flow from the prevailing intervening ancestral populations. As a result, trait changes perish and do not accumulate over time, resulting in stationary fluctuations, until speciation interrupts the gene flow. Although reproducing asexually, microbes do exchange genes through homologous recombination and there is evidence that gene flow plays a critical role in bacterial speciation at least under certain conditions (22–24). Interestingly, the transient trait variation in the initial stasis phase when jumps are absent (the ε term in our model) approximately matches the intraspecific trait variation.

At longer timescales or higher taxonomic levels, trait evolution can be constrained through stabilizing selection exerted by the adaptive zone (25), defined as a set of ecological niches to which a group of species are adapted (2). This will generate the pattern of phylogenetic conservatism where organisms in a clade tend to have similar traits (synapomorphy) and occupy similar habitats. Accordingly, both genome analyses and ecological studies support that ecological coherence exists at higher taxonomic levels in bacteria (26). For example, different bacterial clades have their unique set of genes (27) and analysis of thousands of cultured microbial strains showed that strains related at the genus, family or order levels occupy the same habitat more frequently than expected by chance (28).

Using simulation, we show that our ML framework can detect jumps with an extremely high specificity (0.99) when a posterior probability cutoff of 0.9 is used to predict jumps. On the other hand, the continuous variable rate model struggles to capture the mode of pulsed evolution and performs poorly when benchmarked using the ROC curve. Interestingly, for the first time, we detected two types of jumps in one dataset: small frequent jumps and large rare jumps. This is possible because the large bacterial dataset spans a wide range of macroevolutionary timescales. For example, the bacterial genome tree in our study has a total branch length of 442.9 substitutions/site. For super rare jumps (e.g., genome size jumps with a rate of 0.17 jump per lineage per unit branch length), it is estimated that there are still 75 events in the entire phylogeny. On the other hand, the resolution of our bacterial genome tree is 5⍰10^−5^ substitutions/site, meaning that we can detect jumps that occurs as frequently as 20,000 jumps per unit branch length on average. The large difference in the frequency and size of the jumps suggests that they represent different kinds of evolutionary events. Although our modeling does not stipulate the coupling of cladogenesis and pulsed evolution (as in the classical punctuated equilibrium theory), the rate of the frequent jumps in bacteria (115-634 jumps per lineage per unit branch length or 0.06-0.32 jumps per lineage per Myr) approximates the recently estimated bacterial speciation rate (0.03-0.05 speciation per lineage per Myr) where species is defined as having 99% identical 16S rRNAs (29), suggesting that frequent jumps and the speciation events may be correlated. Two features of the rare jumps fit the description of quantum evolution. First, the rare jumps are fairly large in magnitude, most likely resulting from shifting between major adaptive zones. Second, our test shows that rare jumps happen less frequently than expected at the species level but significantly more frequently than expected at higher taxonomic levels (genus, family and order), suggesting there is a correlation between rare jumps and the origination of higher taxa. Furthermore, some of the predicted rare jumps coincide with known major evolutionary adaptations in bacterial evolution history. A key insight from this observation is that major evolutionary adaptations in bacteria and the origination of major bacterial lineages may happen in quick bursts (quantum evolution) instead of through slow divergence of species over prolonged periods of time (gradual evolution)(30), which is consistent with earlier findings of rapid expansion of major microbial lineages (31, 32).

Microbial genomes are highly dynamic (33, 34). They can change by mutation, gene loss, gene duplication and horizontal gene transfer. Whatever the mechanism, our study suggests that large genome changes happen in episodes of bursts rather than gradually and slowly. These large changes are not due to the simple gain and loss of plasmids as we have excluded plasmids in our study. Chromosomes are in constant exchange with phages, plasmids and other mobile elements and can change by “quantum leaps” in the form of genomic islands (15). It is worth pointing out that jumps in our model represent trait changes that persist over time, not the processes that drive the changes. The rarity of detected jumps does not mean the evolutionary processes (e.g., selection, population bottleneck) that drive the jumps are rare. It merely means the success rate of such jumps is low. The rarity of jumps can result from adaptation to a large environmental shift that happens infrequently, or it can be manifestation of multiple frequent small jumps occurring in quick succession, which is also rare.

In conclusion, our modeling of phylogenetic comparative data shows that pulsed evolution is both prevalent and dominant in bacterial and archaeal genomic traits evolution. The signatures of pulsed evolution detected in this study are consistent with both the punctuated equilibrium and quantum evolution theories. More broadly, our results suggest that pulsed evolution is the rule rather than the exception across the tree of life, despite the drastically different population genetic properties of micro- and macro-organisms.

## Methods

### Bacterial and archaeal phylogeny and genomic traits

We downloaded 10,616 complete bacterial genomes and 263 complete archaeal genomes from the NCBI RefSeq database on September 6, 2018 (Table S8). From each genome we identified either 31 bacterial or 104 archaeal protein-coding marker genes using AMPHORA2 (35) with the default options and constructed a bacterial and an archaeal genome tree based on the concatenated and trimmed protein sequence alignment of the marker genes. We reconstructed the archaeal genome tree using RAxML (version 8.2.11) (36) with the option -m PROTCATLG. Because of its large size, it is impractical to make the bacterial genome tree using RAxML. Instead, we inferred the bacterial genome tree using FastTree (version 2.1.11) (37) with the option -wag -gamma. For better resolution, we re-optimized the branch length of the genome trees with the DNA sequence alignments of the marker genes using RAxML with the option -m GTRGAMMA. We removed genomes with identical alignments, extremely long branches, ambiguous bases or unreliable annotations from the genome trees. For each of the 6,668 bacterial and 247 archaeal genomes remained, we calculated four traits: the ribosomal RNA stem GC% (rRNA GC%), genomic GC%, genome size (excluding plasmids), and the average nitrogen atoms per residual side chain (N-ARSC). We transformed these traits (logit transformation for rRNA GC% and genomic GC%; log transformation for genome size and N-ARSC) to make them comply with the assumption of continuous trait evolution. For conversion from rRNA GC% to the optimal growth temperature, we used the empirical formula determined by Wang et al (9):

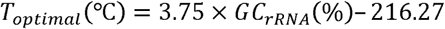

To test the effect of phylogeny uncertainty on our model fitting, we downloaded the bacterial reference phylogeny (release 202) from the Genome Taxonomy Database (GTDB) (17). Additionally, we inferred the phylogeny for the bacterial family Enterobacteriaceae using both FastTree and RAxML.

### Calculating PIC with time-independent variation

Phylogenetically independent contrast (PIC) assumes a Brownian motion (BM) model in which trait variance increases linearly with time (38). However, we observed variation in trait values between genomes that are separated by zero branch length (Fig. 1). Therefore, we introduce time-independent variation into the BM model and denote its variance with ε. When time-independent variation is normally distributed, the PIC between a pair of sister tips is calculated as

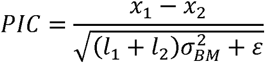

where *x*_*1*_ and *x*_*2*_ are the trait values of the tips, *l*_*1*_ and *l*_*2*_ are their branch lengths to the parent node, and 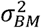 is the rate of Brownian motion. The uncertainty of the parent node’s trait value is calculated as

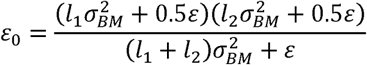

### Calculating pseudo-PIC under pulsed evolution model

When the distribution of trait change over a branch is not normally distributed, the assumption that PIC will follow the standard normal distribution does not hold. To remedy this assumption violation, we calculate pseudo-PIC, an analog of PIC under non-BM models. Specifically, the pseudo-PIC satisfies the equation:

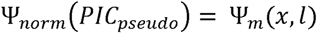

where Ψ_*norm*_ is the standard normal cumulative probability function and Ψ_*m*_(*x,l*) is the cumulative probability function of trait change *x* under the non-BM model given the branch length *l* and its model parameters.

### Testing the pairwise correlation between the four genomic traits

To remove dependence among extant trait values due to shared ancestry, we selected all 2,003 tip pairs in the bacterial genome tree and calculated their PICs for each trait. Using the PICs, we calculated Pearson correlation coefficient *r* and coefficient of determination *R*^2^ for each trait pair. To reduce the bias in correlation introduced by non-normality of the PICs, we also calculated the above statistics for each trait pair using pseudo-PICs that are normally distributed under the pulsed evolution model.

### Quantifying the frequency of extreme trait changes in bacterial evolution history

We tested whether extremely rapid trait changes happen throughout the bacterial evolutionary history. We calculated the relative distance from the root (last common ancestor of bacteria) to a node *i* as

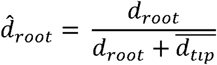

where *d*_*root*_ is the branch length of the node *i* to the root, and 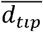 is the average branch length of the node *i* to all its descending tips. It should be noted that a PIC at the node *i* measures the trait difference between its two immediate descending nodes. We binned the PIC based on the relative distance to the root of the node into 7 bins with exponentially distributed boundaries and calculated the frequency of extremely rapid trait changes (|PIC| >= 3) for each bin.

### Segmented linear regression of absolute trait change on branch length

To analyze the trend of trait change, we applied segmented linear regression of the absolute trait changes over branch length as described in Uyeda et al (6). For each trait, we added a small fixed value (0.001) to the absolute trait changes and log-transformed it to obtain approximately normal distribution. To account for uncertainty introduced by ancestral state reconstruction, we adjusted the branch length as described by Felsenstein (38) and log-transformed it as well. When regressing the log-transformed absolute trait changes against the log-transformed adjusted branch lengths, we constrained the slope of the first segment to be zero (to capture the stasis) and allowed the slopes of the remaining regression lines to change at certain breakpoints, but the regression lines had to be continuous (connected). We compared linear regression models with 1 or 2 breakpoints and selected the one with the lowest Akaike Information Criterion (AIC), and used the breakpoints to mark the transitions between different evolution tempos.

### Evaluating pulsed evolution in bacteria and archaea

Using maximum likelihood (ML) method, we tested seven models of trait evolution (Table 1). For pulsed evolution models with more than one compound Poisson process, we restricted the variances of jump sizes between any two jumping processes to be at least 3-fold different. We fitted the models to trait changes between sister nodes given their branch lengths. For internal nodes, we reconstructed their trait values with Felsenstein’s method (38) but took time-independent variation into account. We removed contrasts of zero in rRNA GC% due to their excessiveness in the trait to avoid biased model fitting for pure pulsed evolution (i.e., PE1, PE2 and PE3) models. For all other traits, contrasts of zero were included in the fitting. We calculated confidence intervals for model parameters and statistics by bootstrapping with 50 replicates. We selected the best model using AIC.

We estimated two parameters for each compound Poisson process: frequency *λ*_*i*_ and variance of jump sizes 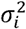, where *i* is the rank of the Poisson process. For further evaluation of pulsed evolution, we calculated the contribution and the relative jump size of each compound Poisson process in pulsed evolution. The contribution of a Poisson process (as proportion of variance explained, PVE) was calculated by

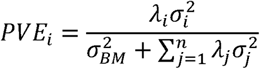

where *i* and *j* are the ranks of Poisson processes, and *n* is the total number of Poisson processes in the model. The relative jump size was calculated as 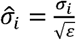. To roughly compare the overall rate of the frequent jumps to the bacterial speciation rate estimated in Myr (29), we calculated the phylogenetically weighted average branch length of all tips to the root in the tree, and then calibrated time assuming the average branch length is equivalent to 3.5 billion years of evolution (39, 40).

### Evaluating the power of detecting jumps in pulsed evolution

To evaluate the power of detecting jumps and distinguishing various models tested in this study, we simulated trait evolution under the seven models in Table 1 with the parameters in Table S2 and fitted these models to the simulated data. We simulated 20 replicates for each model, selected the best models based on AICs of fitted models and counted the frequency that each model was selected as the best model. To determine the effect of sample size on model fitting, from the data simulated using the PE2 model, we randomly sampled 2,250, 750, 250, and 100 pairs of sister nodes without replacement from the full bacterial phylogeny that contains 6,667 sister pairs. We did model selection as described above and counted the frequency that each model was selected as the best model.

### Identifying rare jumps using pulsed evolution model and BayesTraits

We evaluated the performance of using the pulsed evolution model and BayesTraits (41) to detect jumps in simulation. Using 10 replicates of data simulated under the PE2 model, we calculated the posterior probability of having at least one jump between sister nodes as described in Supplementary Text. Using different posterior probability cutoff values, we plotted the receiver operating characteristics (ROC) curve and calculated its area-under-curve (AUC). To test whether the continuous variable rate VRG model can also capture jumps, we applied BayesTraits to estimate the relative rate of evolution for each branch and used a cutoff of the relative rate to predict jumps. Similarly, we used different cutoffs to determine the ROC and AUC of BayesTraits. BayesTraits does not model time-independent trait variation directly. To eliminate the effect of time-independent variation on its rate estimation, for each tip branch in the phylogeny, we added a branch length so that the process variance over the added branch length is equal to half of the time-independent variation between tips. We then applied BayesTraits on the adjusted phylogeny with the VRG model and Bayesian method enabled (mode 7 and 2, respectively).

### Testing the correlation between rare jumps and cladogenesis in bacteria

We identified all contrasts between two congener sister nodes. Using a posterior probability threshold of 0.75, we calculated the frequency of having at least one rare jump in these contrasts for each trait (predicted frequency). We computed the expected distribution of this frequency through simulations using the estimated model parameters of pulsed evolution under the null hypothesis that there is no correlation between jumps and cladogenesis. By comparing the predicted frequency to the expected distribution of the null hypothesis, we calculated the two-sided P-value of the null hypothesis being true at the species level. We repeated the same statistical test at the genus, family and order levels.

## Data availability

The data that support the findings of this study are publicly available from the NCBI RefSeq database under the accession numbers listed in Table S8.

## Code availability

The customized R code and scripts used in this study are provided as Supplementary Files.

## Figures and Tables

**Figure S1.**
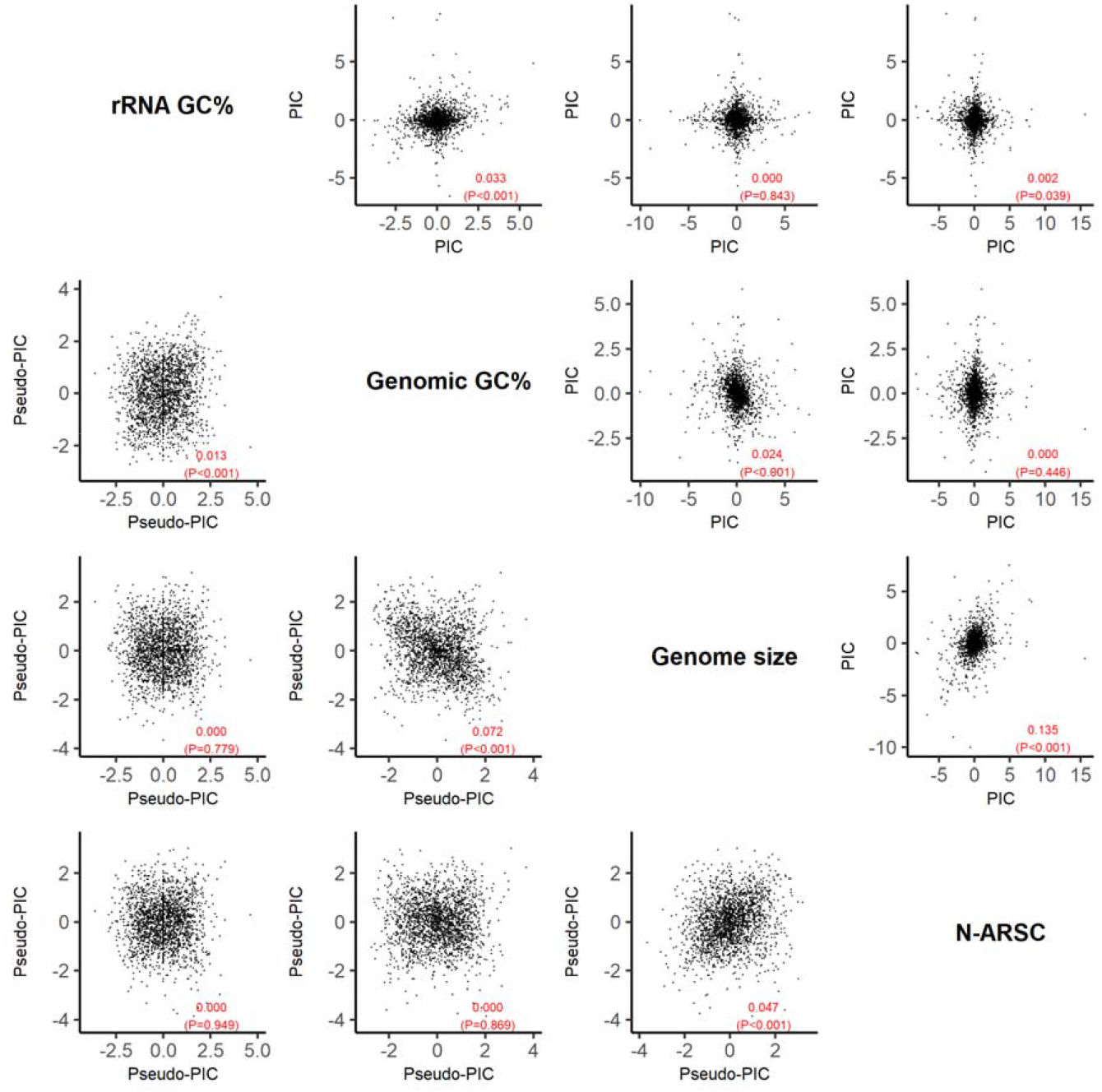
The pairwise correlation between the four genomic traits in bacteria. Upper triangle: the scatter plots of phylogenetically independent contrasts (PICs) between different traits. The corresponding coefficient of determination R^2^ and P value of the correlation are shown in each panel. Lower triangle: the scatter plots of pseudo-PICs between different traits. Names of traits are listed in the diagonal panels.

**Figure S2.**
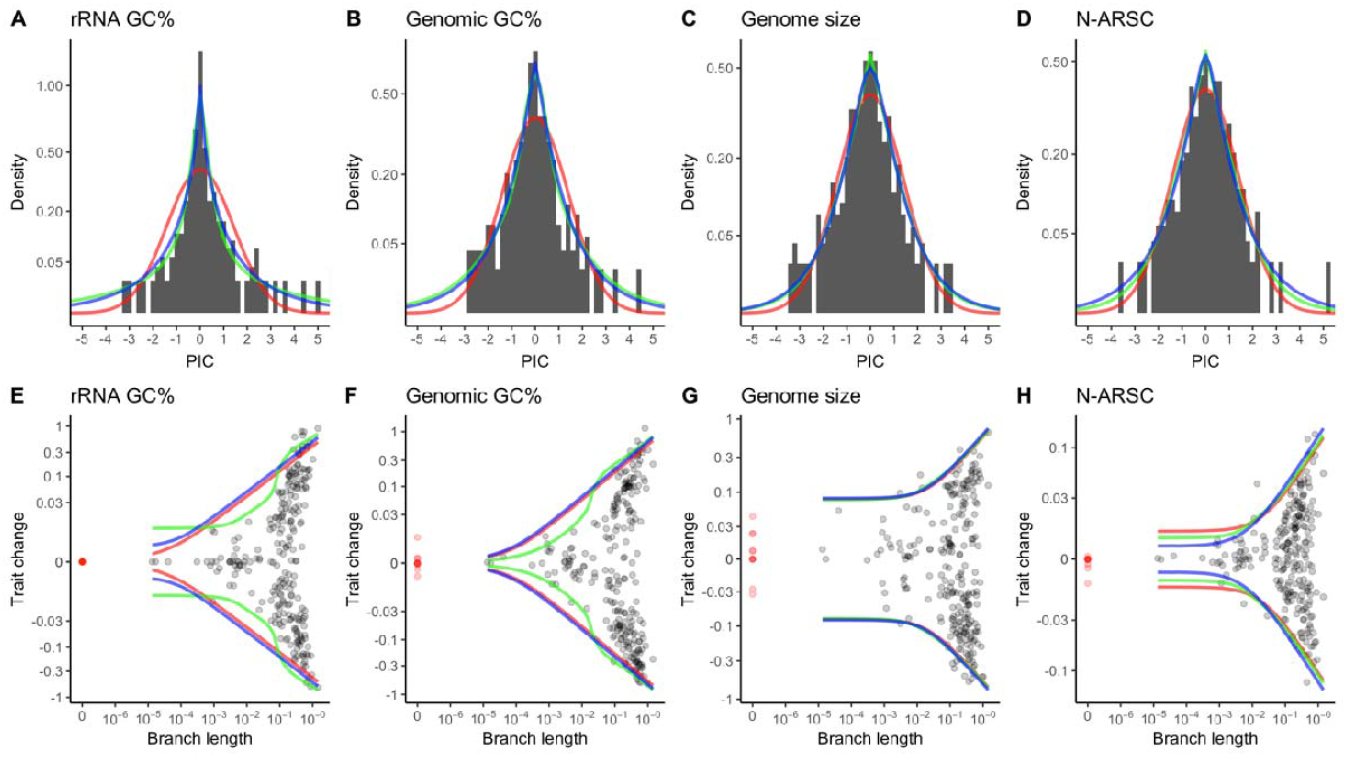
Pulsed evolution models fit archaeal trait evolution better than the BM model and the variable rate model. (A-D) Histogram shows that the PIC (black bar) pattern deviates significantly from the expectation of the BM model (red line), while it is better described by the pulsed evolution model with one or two Poisson processes (PE1 or PE2, green line) and the continuous variable rate model (blue line). The y-axis (density) is square-root transformed to better show the deviation in the frequency of large PICs. (E-H) Patterns of archaeal trait changes at different branch lengths. Trait changes derived from the archaeal phylogeny are shown in black dots. Trait differences derived from genomes separated by zero branch length are shown in red dots. The expected 95% CI of the BM, pulsed evolution (PE1 or PE2) and continuous variable rate models are shown by red, green and blue lines, respectively. The y-axis (trait change) is pseudo-log transformed to better show the trend of trait change in short branches.

**Figure S3.**
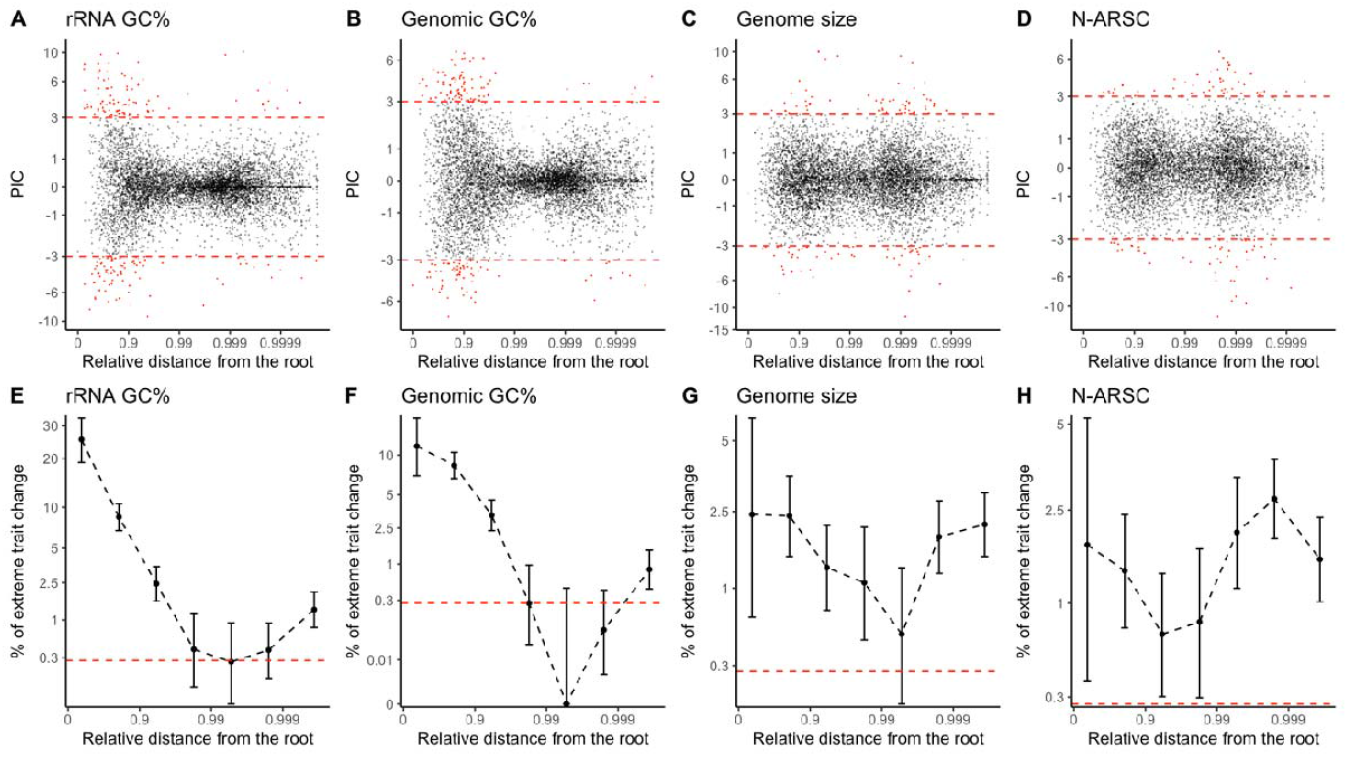
Extremely large PICs occur more frequently than expected by BM throughout the bacterial evolution history. Distributions of PIC over relative distance from the root are shown for (A) ribosomal RNA GC% (B) genomic GC% (C) genome size and (D) N-ARSC. Extremely large PICs (|PIC|>3, outside the red dashed lines) are highlighted in red. Frequencies of the extremely large PIC over relative distance from the root are shown for (E) ribosomal RNA GC% (F) genomic GC% (G) genome size and (H) N-ARSC. The red dashed lines in (E-H) represent the expected frequency of extreme PIC by the BM model. Error bars in (E-H) represent the 95% CI of extremely large PICs’ frequency in each bin.

**Figure S4.**
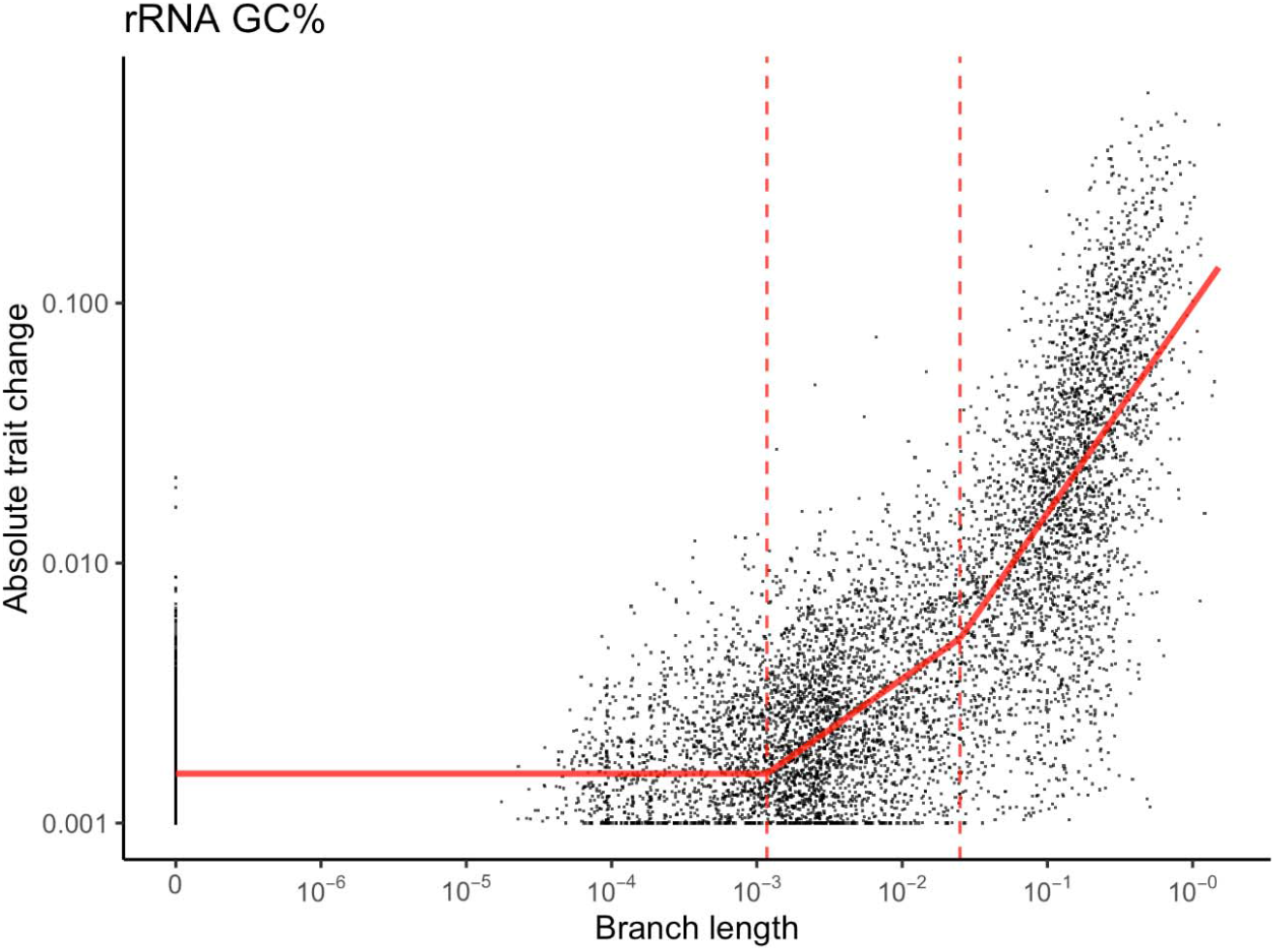
Segmented linear regression analysis indicates that the tempo of evolution changes at various points for ribosomal RNA GC%. The distribution of absolute trait change over branch length is shown for ribosomal RNA GC%. The fitted relationship between the mean absolute trait change and branch length is shown in solid red lines, and the time points where the tempo of evolution changes are marked by dashed red lines.

**Figure S5.**
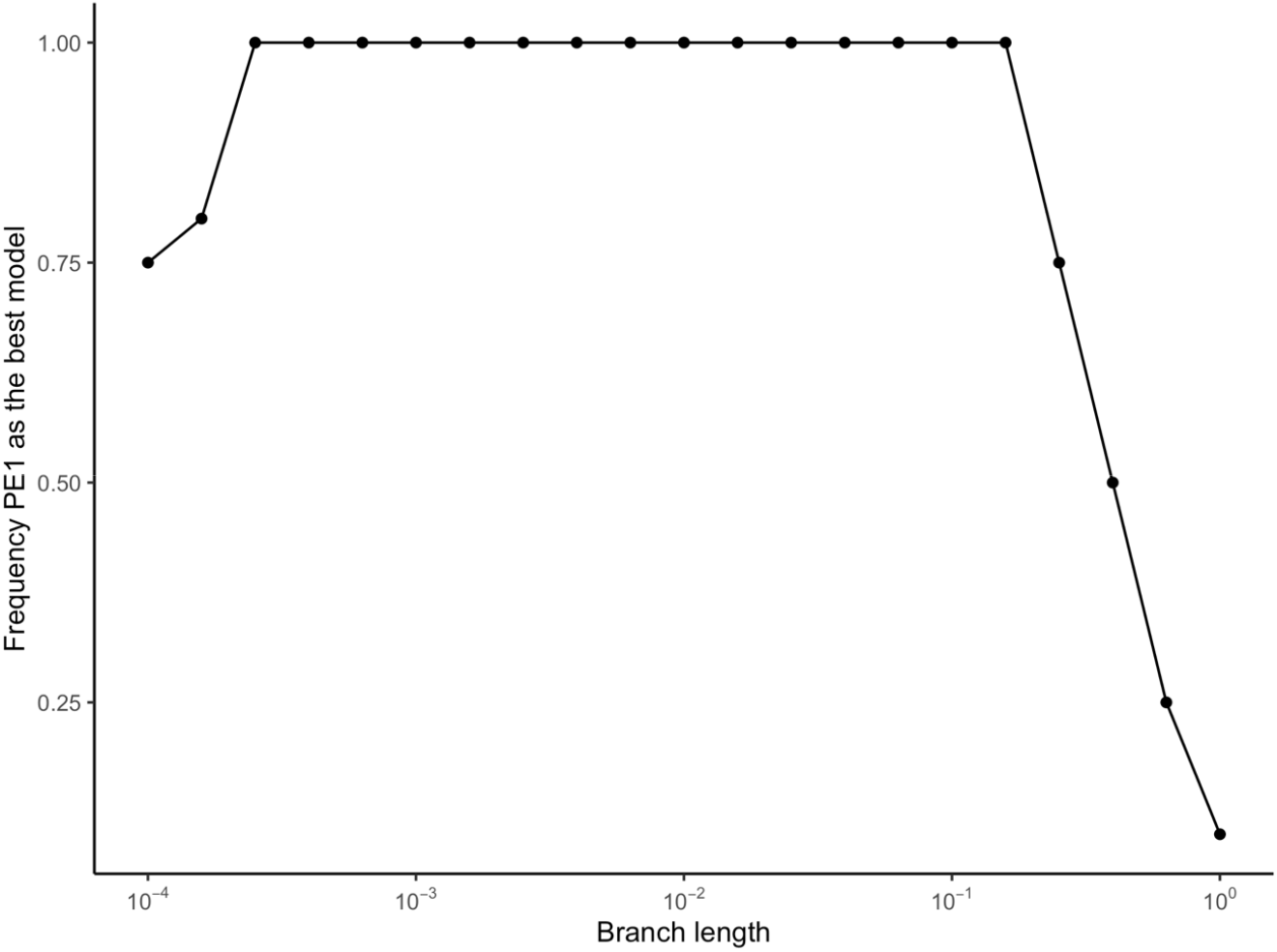
The power of distinguishing the pulsed evolution model and the BM model at different branch lengths. Trait evolution is simulated under the PE1 model. The frequency of correctly selecting the PE1 over the BM model by AIC at different branch lengths is shown in the figure.

**Figure S6.**
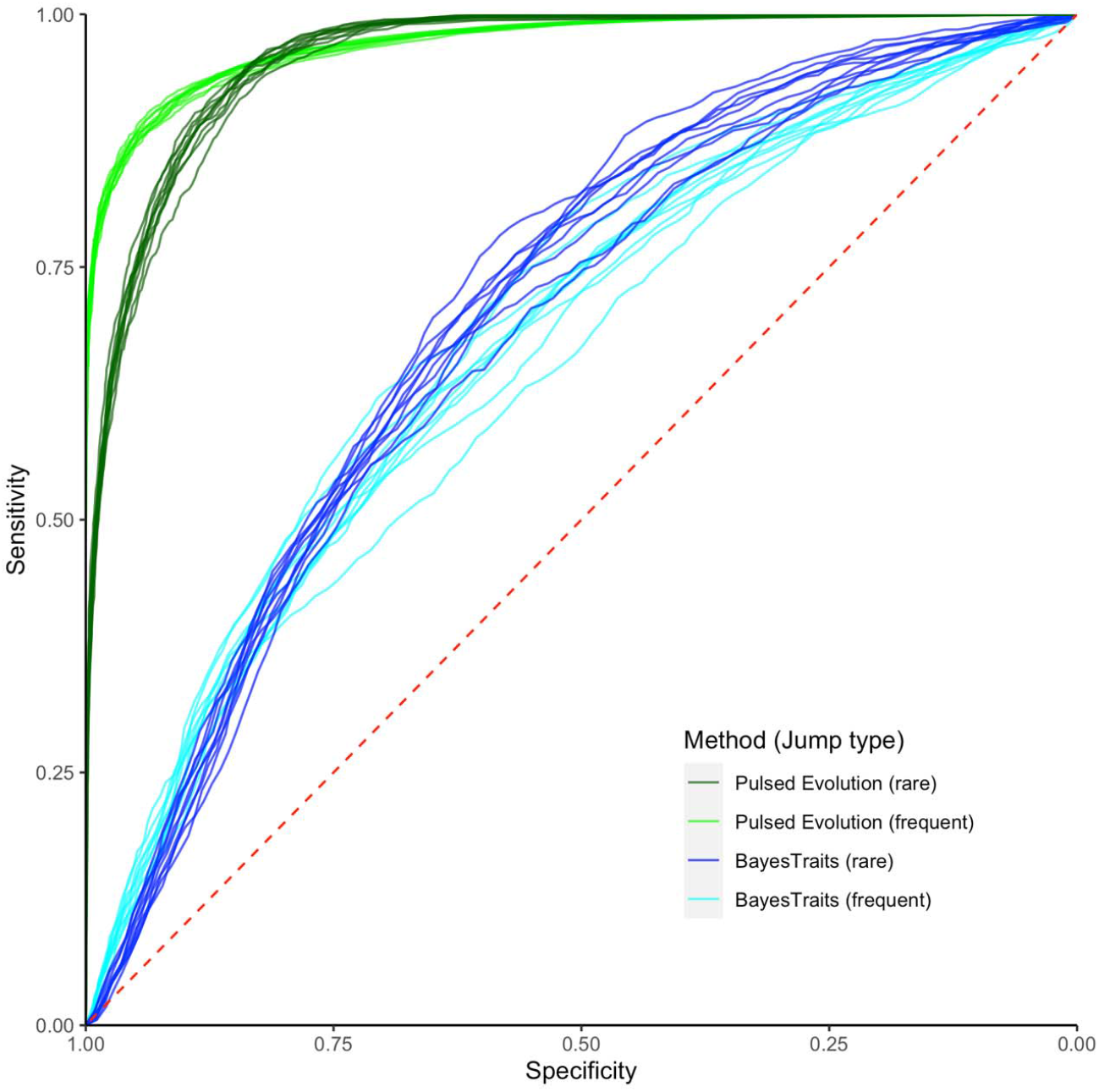
The receiver operating characteristic (ROC) curves for detecting frequent and rare jumps in simulated data using the posterior probability calculated under pulsed evolution or the relative rate estimated by BayesTraits. The four combinations are color-coded by: rare jumps by pulsed evolution (dark green lines), frequent jumps by pulsed evolution (green lines), rare jumps by BayesTraits (blue lines), or frequent jumps by BayesTraits (cyan lines). Ten replicates of trait evolution were simulated under the PE2 model. The red dashed line represents the baseline ROC of random guesses.

## Supplementary Text

### Methods

#### Probability density of trait change under pulsed evolution

For simplicity, we model pulsed evolution similar to the JN model described in Landis and Schraiber (8). Specifically, we assume that the pulsed evolution occurs at a constant rate relative to molecular divergence (branch length), causing sudden changes (jumps) in the trait value. We assume that the sizes of these jumps follow a normal distribution with a mean of zero and a fixed variance. As a result, the pulsed evolution is modeled as a compound Poisson process with normal jumps, with parameters *λ* and *σ*^2^ denoting the frequency and the magnitude (variance) of the jumps, respectively. And between any two tips with branch length *Δl* in the phylogeny, the trait change *Δx*_*PE*_ introduced by pulsed evolution follows the probability density function:

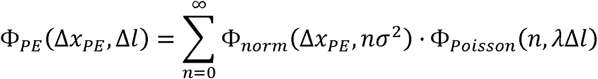

where *n* denotes the number of jumps occurred over the branch,*Φ*_*norm*_(*x,σ*^2^) denotes the normal probability density function of the random variable *x* with a mean of 0 and variance *σ*^2^, and *Φ*_*Poission*_(*n, λ*) denotes the Poisson probability mass function of *n* events with expected occurrence of *λ*. When more than one Poisson process is involved, the probability density function changes to:

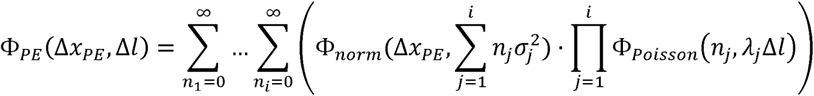

where *i* is the total number of Poisson processes, and *j* denotes the specific compound Poisson process a variable belongs to.

We model the gradual evolution using the classic Brownian motion model with a single parameter 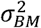 denoting the rate of the gradual trait change. The trait change *Δx*_*BM*_ introduced by gradual evolution follows the probability density function:

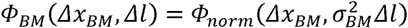

Meanwhile, we have observed trait variations between genomes with identical marker gene alignment (zero branch length), suggesting that branch length does not explain all the trait variation between tips. Consequently, we introduce the time-independent variation to the model. Because the distribution of the observed time-independent variation is leptokurtic (i.e., heavy-tailed, with positive excess kurtosis), we model it with the Laplace distribution with a mean of 0 and scale parameter *b* for simplicity and convenience. The probability density function of *x* following a Laplace distribution is denoted as *Φ*_*Laplace*_(*x,σ*^2^), where *2b*^2^ is its variance and corresponds to the *ε* term in our models (i.e., *ε* = *2b*^2^). Therefore, the time-independent trait change *Δx*_*ε*_ follows the probability density function:

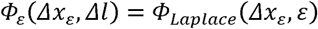

To put these three components together, we first derive the convoluted probability density function of a zero-mean normal distribution and a zero-mean Laplace distribution *Φ*^*^(*x,σ*^2^, *2b*^2^) as

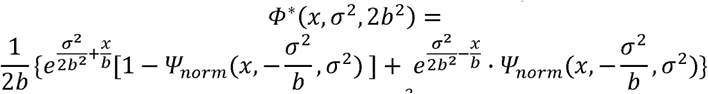

where *σ*^2^is the variance of the normal distribution, *2b*^2^ is the variance of the Laplace distribution and 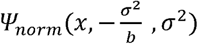 denotes the cumulative distribution function of a normally distributed *x* with a mean of 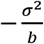 and variance *σ*^2^.

With the convoluted probability density function, we derive that the trait change *Δx* between any two tips with branch length *Δl* in the phylogeny (with one compound Poisson process) follows the probability distribution:

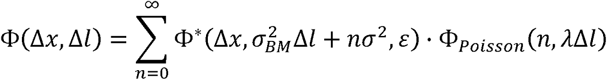

And in the case where more than one compound Poisson process is involved, the probability density function of the trait change becomes:

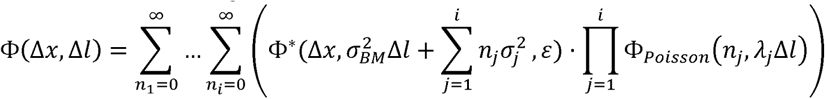

It should be noted that for trait changes that involve internal nodes, the uncertainty of estimated ancestral states (variance) will replace the *ε* term in the above calculation.

#### Probability density of trait change under a continuous variable rate model

As an alternative to pulsed evolution, we also model rate variation using a continuous variable rate model (VRG) implemented in BayesTraits (41). Specifically, for each contrast in the phylogeny, we assume that the evolution rate follows a mixture distribution of a gamma distribution and a constant. The gamma distribution is described by its shape parameter *k* and scale parameter *θ*, and the probability that the rate comes from the gamma distribution is *p*. The constant, on the other hand, is equal to the mean of the gamma distribution, *kθ*. Under this model, the probability density function of trait change is

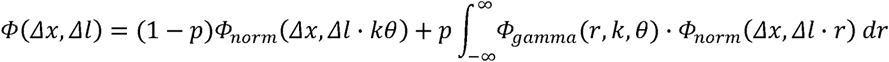

where *Φ*_*gamma*_(*r, k, θ*) denotes the gamma probability density function of random variable *rate r* with shape parameter *k* and scale parameter *θ*.

To simplify the calculation, we discretize the gamma distribution into 20 categories with equal probability weights. After discretization, the probability density function of trait change is

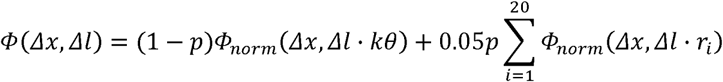

where *r*_*i*_ is the discretized rate.

Like in pulsed evolution, we include a Laplace-distributed time-independent variation in the model. The probability density of trait change with time-independent variation can be calculated through convolution as described above.

#### Posterior probability of jumps between two sister nodes

We calculate the posterior probability of having *n* jumps between two sister nodes. Specifically, with one compound Poisson process, for a trait change *Δx* over branch length *Δl*, the posterior probability of having *n* jumps between the two nodes is

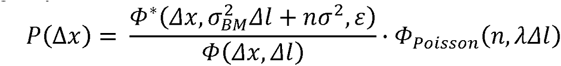

And in the case where more than one Poisson process is involved, the posterior probability of having *n*_l_ jumps in one of the Poisson processes becomes

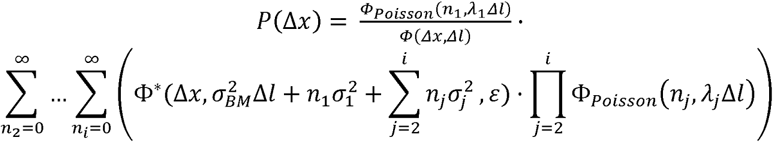

where *i* is the total number of Poisson processes in the pulsed evolution model.

#### Evaluating the power of ML method in distinguishing pulsed evolution and gradual evolution on different branch lengths

Previous study has found that the distribution of trait change under the pulsed evolution model will gradually converges to that under the BM model as the expected number of jumps increases (8). To investigate the power of maximum likelihood model fitting in distinguishing these two models, we simulated trait evolution over 6,667 branches with the PE1 model. We set the frequency of jump to 1,000 per unit branch length with a jump size of 10 and set the variance of time-independent variation to 1. We varied the branch length from short (10^−4^ substitutions/site) to long (1 substitutions/site) at exponentially distributed intervals. For each branch length, we simulated 20 replicates. We fitted the BM and PE1 models on the simulated data and selected the best model by AIC.

#### Implementation

The functions and algorithms described above are implemented in R.

