## Supplementary material for "Microbial genomic trait evolution is dominated by frequent and rare pulsed evolution": readme.txt

1. System requirements
Our custom scripts are built on R 3.6.3. They can run on Linux systems, macOS and Windows. They have been tested on Ubuntu 18.04.1 LTS and Darwin 17.0 (model fitting), and Windows 10 Home (result summary and plotting).

Our custom scripts require the following R packages for analysis:
ape
bbmle
pracma
doParallel
extraDistr
phytools

The following R packages are required for plotting graphs:
ggplot2
ggpubr
ggtree

No non-standard hardware is required to run the custom scripts.

2. Installation
The required packages (except ggtree) can be installed from CRAN by running the following lines in R:
  install.packages("ape")
  install.packages("bbmle")
  install.packages("pracma")
  install.packages("doParallel")
  install.packages("extraDistr")
  install.packages("phytool")
  install.packages("ggplot2")

ggtree should be installed using Bioconductor. Follow the link below to install it:
https://bioconductor.riken.jp/packages/3.10/bioc/html/ggtree.html

It may take up to 1 hour to install all required packages.

3. Usage on Linux and macOS
Scripts can be run using command 'Rscript' in a terminal.

a. fit_5models.R
This script fits 5 evolution models given a phylogeny and trait values:

  Rscript fit_5models.R [tree_file] [trait_file] [save_prefix] [number_of_cores] [exploration_range] [size_constraint] [remove_zero] [Laplace] [AIC_cutoff] [relative_tolerance] [scale]

Arguments: 
[tree_file] is mandatory. Path to the file that contains the phylogeny in the Newwick format.
[trait_file] is mandatory. Path to the file that contains trait values as texts.
[save_prefix] is mandatory. Path and prefix to the output files.
[number_of_cores] is optional. The number of cores to use. Default: 1
[exploration_range] is optional. The fold change in parameter exploration. Default: 10
[size_constraint] is optional. The fold constraint in jump sizes. Default: 10
[remove_zero] is optional. Remove zero-contrasts. Any value greater than 0 is considered TRUE, otherwise FALSE. Default: FALSE
[Laplace] is optional. Use Laplace distribution instead of normal distribution for time-independent variation. Any value greater than 0 is considered TRUE, otherwise FALSE. Default: TRUE
[AIC_cutoff] is optional. The threshold of AIC change before fitting stops. Default: 2
[relative_tolerance] is optional. The relative tolerance in the first phase of Nelder-Mead fitting. Default: 1e-4
[scale] is optional. Manually set the scaling coefficient to a number. Default: automatically scale by data.
 
Note:
An optional argument can be omitted only if no arguments after it will be supplied. If [tree_file] and [trait_file] are identical, the script expects a single RDS file that contains both phylogeny and trait values. 

Output: 
[save_prefix].data.RDS: stores the phylogeny and scaled trait values. 
[save_prefix.models.RDS]: stores the estimated parameter and AIC for each model.

Fitted model parameters and their AICs will also be printed to the standard output at the end.

b. fit_3PE_model.R
This script fits the PE3 model, which is seeded by the PE2+BM model fitted using fit_5models.R:
  
  Rscript fit_3PE_model.R [save_prefix] [number_of_cores] [exploration_range] [size_constraint] [remove_zero] [Laplace] [AIC_cutoff] [relative_tolerance]

Arguments: 
[save_prefix] is mandatory. Path and prefix to the output files.
[number_of_cores] is optional. The number of cores to use. Default: 1
[exploration_range] is optional. The fold change in parameter exploration. Default: 10
[size_constraint] is optional. The fold constraint in jump sizes. Default: 10
[remove_zero] is optional. Remove zero-contrasts. Any value greater than 0 is considered TRUE, otherwise FALSE. Default: FALSE
[Laplace] is optional. Use Laplace distribution instead of normal distribution for time-independent variation. Any value greater than 0 is considered TRUE, otherwise FALSE. Default: TRUE
[AIC_cutoff] is optional. The threshold of AIC change before fitting stops. Default: 2
[relative_tolerance] is optional. The relative tolerance in the first phase of Nelder-Mead fitting. Default: 1e-4

Note:
An optional argument can be omitted only if no arguments after it will be supplied.

Output:
[save_prefix.3PE.RDS]: stores the estimated parameter and AIC for the PE3 model.

Fitted model parameters and the AIC will be printed to the standard output at the end.

c. fit_gamma.R
This script fits the continuous variable rate model VRG.

  Rscript fit_gamma.R [tree_file] [trait_file] [save_prefix] [number_of_cores] [remove_zero] [Laplace] [scale]

Arguments: 
[tree_file] is mandatory. Path to the file that contains the phylogeny in the Newwick format.
[trait_file] is mandatory. Path to the file that contains trait values as texts.
[save_prefix] is mandatory. Path and prefix to the output files.
[number_of_cores] is optional. The number of cores to use. Default: 1
[remove_zero] is optional. Remove zero-contrasts. Any value greater than 0 is considered TRUE, otherwise FALSE. Default: FALSE
[Laplace] is optional. Use Laplace distribution instead of normal distribution for time-independent variation. Any value greater than 0 is considered TRUE, otherwise FALSE. Default: TRUE
[scale] is optional. Manually set the scaling coefficient to a number. Default: automatically scale by data.

Note:
An optional argument can be omitted only if no arguments after it will be supplied. If [tree_file] and [trait_file] are identical, the script expects a single RDS file that contains both phylogeny and trait values. 

Output:
[save_prefix.gamma.RDS]: stores the estimated parameter and AIC for the PE3 model.
Fitted model parameters and the AIC will be printed to the standard output at the end.

d. bootstrap_best_model.R
This script conducts bootstrapping for a specified model to estimate the confidence intervals of estimated model parameters.

  Rscript bootstrap_best_model.R [save_prefix] [number_of_cores] [model] [bootstrap_size] [size_constraint] [remove_zero] [Laplace] [AIC_cutoff] [relative_tolerance]

Arguments:
[save_prefix] is mandatory. Path and prefix to the output files.
[number_of_cores] is optional. The number of cores to use. Default: 1
[model] is optional. The model to fit in bootstrapping. The tag used for the models are: 'bme' for BM, 'pee' for PE1, 'pebme' for PE1+BM, 'pee2' for PE2, 'pebme2' for PE2+BM, 'pee3' for PE3.  Default: automatically determined by AIC
[bootstrap_size] is optional. The number of replicates to run. Default: 50
[size_constraint] is optional. The fold constraint in jump sizes. Default: 10
[remove_zero] is optional. Remove zero-contrasts. Any value greater than 0 is considered TRUE, otherwise FALSE. Default: FALSE
[Laplace] is optional. Use Laplace distribution instead of normal distribution for time-independent variation. Any value greater than 0 is considered TRUE, otherwise FALSE. Default: TRUE
[AIC_cutoff] is optional. The threshold of AIC change before fitting stops. Default: 2
[relative_tolerance] is optional. The relative tolerance in the first phase of Nelder-Mead fitting. Default: 1e-4

Note:
An optional argument can be omitted only if no arguments after it will be supplied.

Output:
[save_prefix.bootstrap.RDS]: stores the bootstarpped parameters.

e. correlation_by_posterior_probability.R
This script conducts correlation analysis.

  Rscript correlation_by_posterior_probability.R [save_prefix] [tip_sppid] [lineage_table] [model] [number_of_cores]

Arguments:
[save_prefix] is mandatory. Path and prefix to the output files.
[tip_sppid] is mandatory. Path to the file that contains the species ID of tips.
[lineage_table] is mandatory. Path to the file that contains the species lineage information.
[model] is mandatory. The model to use in the analysis.
[number_of_cores] is optional. The number of cores to use. Default: 1

Output:
[save_prefix.clado.RDS]: stores the result of correlation analysis.

4. Usage on Windows
On Windows, we suggest to run the custom scripts in Rstudio (e.g. RStudio Version 1.3.1093). 

Manually edit the following lines in the scripts.
Set the current working directory to the folder that contains the scripts:
  setwd("[path_to_folder_containing_the_script]")

Open the script you would like to run in the editor and change the following line
  arg=commandArgs(TRUE)
to a custom line that includes the arguments described in the Linux/macOS section. For example:
  arg=c("Data/kingdom/combo.cds.HQ.6668.tre","Data/data/genomic_GC.logit.txt","Results/Bacterial_result/bac.gGC")
  
And change the following line
  fullarg = commandArgs(FALSE)
to
  fullarg = c("--file=./Placeholder.R")
  
Then source the script to run it:
  source("script_to_run")

5. Demo
Below is a demo on running the scripts on Linux or macOS:

To fit 5 models, run the script using the included data in the demo folder: bacteria.demo.tre and bacteria.demo.trait.txt

  Rscript Scripts/fit_5models.R demo/bacteria.demo.tre demo/bacteria.demo.trait.txt demo/bac.demo 4 3 3 -1

To fit the PE3 model after that, run
  
  Rscript Scripts/fit_3PE_model.R demo/bac.demo 4 3 3 -1

It takes about 1.5 to 2 hours to finish running this demo on a typical desktop computer. The expected output files are included in the demo folder: bac.demo.models.RDS and bac.demo.3PE.RDS. The models in the standard output are: 'bme' for BM, 'pee' for PE1, 'pebme' for PE1+BM, 'pee2' for PE2, 'pebme2' for PE2+BM and 'pee3' for PE3.

6. Reproduce analyses of the empirical data
Note that the following procedures will overwrite the downloaded supplementary files in the folder, e.g., files in Results/Bacteria_result/.

To reproduce the model fitting on bacterial traits along the full phylogeny, run the following commands with 15 cores
a. Genomic GC%
  Rscript Scripts/fit_5models.R Data/kingdom/combo.cds.HQ.6668.tre Data/data/genomic_GC.logit.txt Results/Bacterial_result/bac.gGC 15 3 3 -1
  Rscript Scripts/fit_3PE_model.R Results/Bacteria_result/bac.gGC 15 3 3 -1
  Rscript Scripts/fit_gamma.R Data/kingdom/combo.cds.HQ.6668.tre Data/data/genomic_GC.logit.txt Results/Bacterial_result/bac.gGC 15 -1

b. 16S rRNA GC%  
  Rscript Scripts/fit_5models.R Data/kingdom/combo.cds.HQ.6668.tre Data/data/rRNA_stem_GC.logit.txt Results/Bacterial_result/bac.rGC 15 3 3 1
  Rscript Scripts/fit_3PE_model.R Results/Bacteria_result/bac.rGC 15 3 3 1
  Rscript Scripts/fit_gamma.R Data/kingdom/combo.cds.HQ.6668.tre Data/data/rRNA_stem_GC.logit.txt Results/Bacterial_result/bac.rGC 15 1
  
c. Genome size
  Rscript Scripts/fit_5models.R Data/kingdom/combo.cds.HQ.6668.tre Data/data/genome_size.log.txt Results/Bacterial_result/bac.GS 15 3 3 -1
  Rscript Scripts/fit_3PE_model.R Results/Bacteria_result/bac.GS 15 3 3 -1
  Rscript Scripts/fit_gamma.R Data/kingdom/combo.cds.HQ.6668.tre Data/data/genome_size.log.txt Results/Bacterial_result/bac.GS 15 -1

d. N-ARSC
  Rscript Scripts/fit_5models.R Data/kingdom/combo.cds.HQ.6668.tre Data/data/N_ARSC.log.txt Results/Bacterial_result/bac.N 15 3 3 -1
  Rscript Scripts/fit_3PE_model.R Results/Bacteria_result/bac.N 15 3 3 -1
  Rscript Scripts/fit_gamma.R Data/kingdom/combo.cds.HQ.6668.tre Data/data/N_ARSC.log.txt Results/Bacterial_result/bac.N 15 -1

To reproduce the bootstrapping results on the best model, run
  Rscript Scripts/bootstrap_best_model.multithread.R Results/Bacterial_result/bac.gGC 15 pee2 50 3 -1
  Rscript Scripts/bootstrap_best_model.multithread.R Results/Bacterial_result/bac.rGC 15 pee2 50 3 1
  Rscript Scripts/bootstrap_best_model.multithread.R Results/Bacterial_result/bac.GS 15 pee3 50 3 -1
  Rscript Scripts/bootstrap_best_model.multithread.R Results/Bacterial_result/bac.N 15 pee3 50 3 -1
(Note that it may take several days to finish bootstrapping even run in parallel)

To reproduce the correlation analysis, run
  Rscript Scripts/correlation_by_posterior_probability.R Results/Bacterial_result/bac.gGC Data/taxonomy/acc2taxid.RDS Data/taxonomy/spp.tax.df.RDS pee2 15 
  Rscript Scripts/correlation_by_posterior_probability.R Results/Bacterial_result/bac.rGC Data/taxonomy/acc2taxid.RDS Data/taxonomy/spp.tax.df.RDS pee2 15 
  Rscript Scripts/correlation_by_posterior_probability.R Results/Bacterial_result/bac.GS Data/taxonomy/acc2taxid.RDS Data/taxonomy/spp.tax.df.RDS pee3 15 
  Rscript Scripts/correlation_by_posterior_probability.R Results/Bacterial_result/bac.N Data/taxonomy/acc2taxid.RDS Data/taxonomy/spp.tax.df.RDS pee3 15 
(Note that it may take a day or more to run the correlation analysis even run in parallel)

To re-simulate trait evolution under different models, run
  Rscript simulate_with_track.R

The figures in the manuscript can be reproduced by setting the working directory to Scripts/ and sourcing the following scripts in RStudio:

  source("Fig_1.R")
  source("Fig_2.R")
  source("Fig_S1.R")
  source("Fig_S2.R")
  source("Fig_S3S4.R")
  source("Fig_S5.R")
  source("Fig_S6.R")
  
