## Supplementary material for "Microbial genomic trait evolution is dominated by frequent and rare pulsed evolution": ape_addition.R

```
# This script contains some useful extensions for tree manipulation based on the ape package.
# By Yingnan Gao, 20181229; 

# load required library
library(ape)
#
getLatestAncestor = function (phy, x){
  # x should be the index of the focal node, not the node/tip label
  if (x == Ntip(phy) + 1) return(NA)
  i <- which(phy$edge[, 2] == x)
  return(phy$edge[i, 1])
}
#
getAncestor = function (phy, x,level=1,trace=FALSE){
  # returning the ancestor of a specified degree, compatiable with other functions with the same name
  # e.g. Kemble's 2012 paper
  if(trace) all.anc = numeric(level)
  anc = x
  for(i in 1:level){
    anc = getLatestAncestor(phy=phy,x=anc)
    if(trace) all.anc[i]=anc
    if(is.na(anc)) break
  }
  if(trace){
    return(all.anc[-which(all.anc==0)])
  }else{
    return(anc) 
    }
}
#
getRoot = function(phy){
  return(Ntip(phy) + 1)
}
#
getNextDescendants = function (phy, x){
  # x should be the index
  if (x <= Ntip(phy)) 
    return(NA)
  i <- which(phy$edge[, 1] == x)
  return(phy$edge[i, 2])
}
#
findSisters <- function(phy,x){
  # x should be the index
  parent = getLatestAncestor(phy,x)
  if(is.na(parent)) return(NA)
  twin = getNextDescendants(phy,parent)
  sis = twin[twin!=x]
  return(sis)
}
#
findEdges = function(phy,node,ancestor = FALSE){
  # node should be the index
  if(ancestor){
    edge.index = which(phy$edge[,2]==node)
  }else{
    edge.index = which(phy$edge[,1]==node)
  }
  return(edge.index)
}
#
getConnected = function(phy,x){
  conn = c(phy$edge[phy$edge[,2]==x,1],phy$edge[phy$edge[,1]==x,2])
  return(conn)
}
#
simulateBMTrait = function(phy,start=0){
  trait.sim = numeric(length = Ntip(phy)+Nnode(phy))+start
  if(is.null(phy$node.label)){
    names(trait.sim) = c(phy$tip.label,as.character(Ntip(phy)+(1:phy$Nnode)))
  }else{
    names(trait.sim) = c(phy$tip.label,phy$node.label)
  }
  phy = reorder.phylo(phy,order = "postorder")
  trait.delta = rnorm(length(phy$edge.length))*sqrt(phy$edge.length)
  for(i in 1:length(phy$edge.length)){
    l = length(phy$edge.length)+1-i
    trait.sim[phy$edge[l,2]] = trait.sim[phy$edge[l,1]] + trait.delta[l]
  }
  return(trait.sim)
}
#
predTraitByPIC = function(phy, trait){
  # predicts trait value from known extant specis using PIC (BM-ML)
  # this function is modified from Kembel's function in picante package for a more appropriate SE
  # get names of known and unknown species
  refSpecies = names(trait)
  tarSpecies = phy$tip.label[is.na(match(phy$tip.label,refSpecies))]
  #
  pred = as.data.frame(matrix(nrow=length(tarSpecies), ncol=2, dimnames=list(tarSpecies, c("estimate","se"))))
  #
  for (i in tarSpecies) {
    # keep only the focal target species and the references
    this.tree = drop.tip(phy = phy,tip = tarSpecies[tarSpecies!=i])
    # reroot the tree at the focal target species
    this.tree = root(this.tree, i, resolve.root=FALSE)
    # get branch length leading to the focal species in the rerooted tree for standard errors
    focal.bl = this.tree$edge.length[which(this.tree$edge[,2]==which(this.tree$tip.label==i))]
    # trim the focal species
    this.tree = drop.tip(this.tree, i)
    # get the adjusted branch lengths for the root estimate
    pic.tree = pic(x = trait,phy = this.tree,rescaled.tree = TRUE)$rescaled.tree
    root.bl = pic.tree$edge.length[pic.tree$edge[,1]==getRoot(pic.tree)]
    # use PIC (BM-ML analytical solution) to estimate trait value and error
    est = ace(x = trait[this.tree$tip.label], phy = this.tree, method="pic",type = "continuous")$ace[1]
    se = sqrt(1/sum(1/root.bl) + focal.bl)
    pred[i,] <- data.frame(estimate=est, se=se)
  }
  return(pred)
}
#
continuousAceByPIC = function(x,phy){
  res = numeric(phy$Nnode)
  if(is.null(phy$node.label)){
    phy$node.label = as.character(Ntip(phy)+1:phy$Nnode)
    names(res) = phy$node.label
  }else{
    names(res) = phy$node.label
  }
  se = res
  res[1] = ace(phy = phy,x = x[phy$tip.label],type = "continuous",method = "pic")$ace[1]
  pic.phy = pic(x = x[phy$tip.label],phy = phy,rescaled.tree = TRUE)[[2]]
  se[1] = 1/sum(1/pic.phy$edge.length[pic.phy$edge[,1]==(Ntip(pic.phy)+1)])
  for(i in phy$node.label[-1]){
    new.phy = root(phy,node=which(phy$node.label==i)+Ntip(phy),resolve.root=TRUE)
    res[i] = ace(phy = new.phy,x = x[new.phy$tip.label],type = "continuous",method = "pic")$ace[1]
    pic.phy = pic(x = x[new.phy$tip.label],phy = new.phy,rescaled.tree = TRUE)[[2]]
    se[i] = sqrt(1/sum(1/pic.phy$edge.length[pic.phy$edge[,1]==(Ntip(pic.phy)+1)]))
  }
  return(data.frame(est=res,se=se,row.names = phy$node.label))
}
#
```
