## Supplementary material for "Microbial genomic trait evolution is dominated by frequent and rare pulsed evolution": bootstrap_best_model.R

```
# read in key arguments for the fitting
arg=commandArgs(TRUE)
fullarg = commandArgs(FALSE)
# setup the environment for model fitting
work.dir = getwd()
source.dir = fullarg[grep(pattern = "--file=",x = fullarg,fixed = TRUE)[1]]
source.dir = dirname(substr(source.dir,8,nchar(source.dir)))
setwd(source.dir)
source("fitPE_functions.R")
setwd(work.dir)
# interpret the arguments
# the job to continue analyses on
job.name = arg[1]
if(is.na(job.name)) stop("Job name is required and cannot be omitted!")
# read in phylogeny and trait from 1 RDS file
data.file = sprintf("%s.data.RDS",job.name)
tree = readRDS(data.file)
trait = tree$dat
tree = tree$phy
# read in fitting results
model.file = sprintf("%s.models.RDS",job.name)
res = readRDS(model.file)
pe3.file = sprintf("%s.3PE.RDS",job.name)
scale.coef = res$scale
if(file.exists(pe3.file)){
  pe3.res = readRDS(pe3.file)
  model.AIC = c(res$stat,pe3.res$stat)
  model.params = c(res$params,pe3.res$params)
}else{
  model.AIC = res$stat
  model.params = res$params
}
# how many cores to use in the fitting
numCores=ceiling(as.numeric(arg[2]))
if(is.na(numCores)) numCores=1
# which is the best model
best.model = arg[3]
if(is.na(best.model)) best.model = which.min(model.AIC)
best.params = model.params[[best.model]]
# how many bootstraps to run for a confidence interval
numBoot = ceiling(as.numeric(arg[4]))
if(is.na(numBoot)) numBoot = 50
# set manual fold-constraint between sizes
size.ratio = as.numeric(arg[5])
if(is.na(size.ratio)) size.ratio=10
#
ignore.zero = (as.numeric(arg[6])>0)
if(is.na(ignore.zero)) ignore.zero = FALSE
#
laplace = (as.numeric(arg[7])>0)
if(is.na(laplace)) laplace=TRUE
#
# when to stop the optimization in while loop
AIC.cutoff = as.numeric(arg[8])
if(is.na(AIC.cutoff)) AIC.cutoff=2
#
reltol = as.numeric(arg[9])
if(is.na(reltol)) reltol = 1e-4
# initialize key arguments
model.functions = c(bme="fitPE.phy",pee = "fitPE.phy",pebme ="fitPE.phy",
                    pee2= "fitPE2.phy",pebme2="fitPE2.phy",
                    pee3 = "fitPE3.phy")
fixed.params = list(bme=list(lambda=log(0),size=log(0)),
                    pee = list(sigma=log(0)),pebme =list(),
                    pee2= list(sigma=log(0)),pebme2=list(),
                    pee3= list(sigma = log(0)))
ini.functions = c(bme="initialize.parameters.SP",
                  pee = "initialize.parameters.SP",pebme ="initialize.parameters.SP",
                  pee2= "initialize.parameters.MP",pebme2="initialize.parameters.MP",
                  pee3 = "initialize.parameters.TP")
# set bootstrapped data
bootstrap.index = lapply(1:numBoot,function(i){
  sample(x = 1:Nnode(tree),size = Nnode(tree),replace = TRUE)
})
# fit bootstrapped data
bootstrap.res = lapply(1:numBoot,function(i){
  start.params = do.call(ini.functions[best.model],list(best.params))
  if(any(sapply(start.params,is.nan))) next
  mdl= fit.model.byNM.2step(fit.func = model.functions[[best.model]],
                            start.params = list(start.value=start.params),
                            params1 =list(fixed=fixed.params[[best.model]],size.ratio=size.ratio,ignore.zero=ignore.zero,laplace = laplace,bootstrap = bootstrap.index[[i]],
                                          phy=tree,x=sqrt(scale.coef)*trait[tree$tip.label],numCores=numCores,reltol=reltol),
                            params2 =list(fixed=fixed.params[[best.model]],size.ratio=size.ratio,ignore.zero=ignore.zero,laplace = laplace,bootstrap = bootstrap.index[[i]],
                                          phy=tree,x=sqrt(scale.coef)*trait[tree$tip.label],numCores=numCores),
                            AIC.func = "get.AIC",
                            params.func = "get.params",
                            ini.func = ini.functions[best.model],
                            AIC.cutoff = AIC.cutoff)
  return(mdl)
})
# save results
saveRDS(list(model=names(model.functions)[best.model],res = bootstrap.res,index = bootstrap.index),
        sprintf("%s.bootstrap.RDS",job.name))
#
stopCluster(cl)
#
warnings()
```
