## Supplementary material for "Microbial genomic trait evolution is dominated by frequent and rare pulsed evolution": correlation_by_posterior_probability.R

```
# read in key arguments for the fitting
arg=commandArgs(TRUE)
fullarg = commandArgs(FALSE)
# setup the environment for model fitting
work.dir = getwd()
source.dir = fullarg[grep(pattern = "--file=",x = fullarg,fixed = TRUE)[1]]
source.dir = dirname(substr(source.dir,8,nchar(source.dir)))
setwd(source.dir)
source("fitPE_functions.R")
setwd(work.dir)
library(phytools)
# interpret the arguments
# the job to continue analyses on
job.name = arg[1]
if(is.na(job.name)) stop("Job name is required and cannot be omitted!")
# read in phylogeny and trait from 1 RDS file
data.file = sprintf("%s.data.RDS",job.name)
tree = readRDS(data.file)
trait = tree$dat
tree = tree$phy
# read in fitting results
model.file = sprintf("%s.models.RDS",job.name)
pe3.file = sprintf("%s.3PE.RDS",job.name)
res = readRDS(model.file)
res.3pe = readRDS(pe3.file)
scale.coef = res$scale
model.AIC = c(res$stat,res.3pe$stat)
model.params = c(res$params,res.3pe$params)#
# read in taxonomy information
taxids = readRDS(arg[2])
taxlin = readRDS(arg[3])
# which model to test correlation
best.model = arg[4]
best.params = model.params[[best.model]]
# how many cores to use
numCores=ceiling(as.numeric(arg[5]))
if(is.na(numCores)) numCores=1
# getting ancestral states using the best model
numPE = length(best.params)/2-1
rate = sum(sapply(1:numPE,function(i){
  best.params[2*i-1]*best.params[2*i]
}))+best.params[2*numPE+1]
best.ancs = reconstructAncestralStates(phy = tree,x = sqrt(scale.coef)*trait[tree$tip.label],rate = unname(rate),epsilon = unname(best.params["epsilon"]))
# setting functions to use
model.functions = c(bme="posteriorJumpProbability",pee = "posteriorJumpProbability",pebme ="posteriorJumpProbability",
                    pee2= "posteriorJumpProbability.laplace2",pebme2="posteriorJumpProbability.laplace2",pee3 = "posteriorJumpProbability.laplace3")
#
ini.functions = c(bme="simulatePET.phy",
                  pee = "simulatePET.phy",pebme ="simulatePET.phy",
                  pee2= "simulatePET2.phy",pebme2="simulatePET2.phy",
                  pee3= "simulatePET3.phy")
rfunctions = c(bme="rPEpoislaplace",
               pee = "rPEpoislaplace",pebme ="rPEpoislaplace",
               pee2= "rPEpois2laplace",pebme2="rPEpois2laplace",
               pee3= "rPEpois3laplace")
numPE = c(bme=1,
          pee = 1,pebme =1,
          pee2= 2,pebme2=2,
          pee3= 3)
# Setup parallel backend to use many processors
cores=detectCores()
cat(sprintf("There's %d cores available\n",cores))
cl <- makeCluster(min(cores[1]-1,numCores),type = "FORK") # Not to overload
cat(sprintf("Using %d cores\n",min(cores[1]-1,numCores)))
registerDoParallel(cl)

# calculate empirical posterior probabilities
pps = parSapply(cl = cl,X = 1:length(best.ancs$contrast),FUN = function(i){
  do.call(model.functions[best.model],c(list(x=best.ancs$contrast[i],t=best.ancs$l[i],epsilon=best.ancs$epsilon[i]),as.list(best.params[-length(best.params)])))
})
# calculate simulated posterior probabilities under the null hypothesis
sim.pps = parLapply(cl=cl,1:1000,function(n){
  sim.trait = do.call(ini.functions[best.model],c(list(phy=tree),as.list(best.params)))
  sim.ancs = reconstructAncestralStates(phy = tree,x = sim.trait[tree$tip.label],rate = unname(rate),epsilon = unname(best.params["epsilon"]))
  this.pps = sapply(1:length(sim.ancs$contrast),FUN = function(i){
    do.call(model.functions[best.model],c(list(x=sim.ancs$contrast[i],t=sim.ancs$l[i],epsilon=sim.ancs$epsilon[i]),as.list(best.params[-length(best.params)])))
  })
  return(this.pps)
})
# check taxonomy
undef.tips = which(is.na(match(taxids[tree$tip.label],taxlin$taxid)))
if(length(undef.tips)>0) stop("There's taxonomically undefined tip(s) in the tree!")
# at each level, find the contrasts that lead to a certain clade
levels=c(class=4,order=5,family=6,genus=7)
clado.contrast = lapply(levels,function(tax){
  tip.tax = as.character(taxlin[match(taxids[tree$tip.label],taxlin$taxid),tax])
  tip.tax.higher = as.character(taxlin[match(taxids[tree$tip.label],taxlin$taxid),tax-1])
  tip.tax[is.na(tip.tax)] = "Unclassified"
  count.tax = sort(table(tip.tax),decreasing = TRUE)
  unique.tax = names(count.tax)
  unique.tax = unique.tax[unique.tax!="Unclassified"]
  leading.node = sapply(unique.tax,function(x){
    node = getMRCA(phy = tree,tip = which(tip.tax==x))
    if(is.null(node)) node = which(tip.tax==x)
    return(node)
  })
  if(tax<7) leading.node = leading.node[1:sum(table(tip.tax)>=2)]
  leading.status.mono = sapply(leading.node,function(x){
    if(x<=Ntip(tree)) return(TRUE)
    subclade = extract.clade(tree,x)
    subtips = match(subclade$tip.label,tree$tip.label)
    return(length(unique(tip.tax[subtips]))==1)
  })
  leading.node = leading.node[leading.status.mono]
  contrast.node = unique(sapply(leading.node,getAncestor,phy=tree))
  contrast.node.mono = sapply(contrast.node,function(x){
    subclade = extract.clade(tree,x)
    subtips = match(subclade$tip.label,tree$tip.label)
    return(length(unique(tip.tax.higher[subtips]))==1)
  })
  if(tax==7) contrast.node=contrast.node[contrast.node.mono]
  return(contrast.node)
})
# simulation for power analysis
node.sim.pps = lapply(seq(0,1,0.25),function(k){
  print(k)
  level.res = lapply(clado.contrast,function(x){
    print(length(x))
    this.l = best.ancs$l[x-Ntip(tree)]
    this.epsilon = best.ancs$epsilon[x-Ntip(tree)]
    this.perm = parLapply(cl=cl,1:1000,function(n){
      this.dx = do.call(rfunctions[best.model],c(list(n=length(this.l),t=this.l,epsilon=this.epsilon),as.list(best.params[-length(best.params)])))
      this.jump = rbinom(n = length(this.l),size = 1,prob = k)
      this.pps = lapply(1:numPE[best.model],function(j){
        this.ddx = this.dx + rnorm(n = length(this.l),mean = 0,sd = sqrt(this.jump*best.params[2*j]))
        sapply(1:length(this.l),function(i){
          do.call(model.functions[best.model],c(list(x=this.ddx[i],t=this.l[i],epsilon=this.epsilon[i]),as.list(best.params[-length(best.params)])))
        })
      })
      names(this.pps) = as.character(1:numPE[best.model])
      return(this.pps)
    })
  })
  names(level.res) = names(levels)
  return(level.res)
})
names(node.sim.pps) = as.character(seq(0,1,0.25))
stopCluster(cl)
# calculate power
sim.power = lapply(node.sim.pps,function(k.data){
  this.k.power = lapply(1:length(k.data),function(l.idx){
    print(l.idx)
    level.data = k.data[[l.idx]]
    this.contrast.idx = clado.contrast[[l.idx]]
    this.power = lapply(level.data,function(jump.data){
      sapply(jump.data,function(this.jump){
        #
        power.ratio = apply(this.jump,1,function(x){
          sum(x<0.25)/length(x)
        })
        #
        simulated.ratio = sapply(sim.pps,function(y){
          sub.y = y[,this.contrast.idx-Ntip(tree)]
          sapply(1:dim(y)[1],function(i){
            sum(sub.y[i,]<0.25)/length(this.contrast.idx)
          })
        })
        #
        cumulative.prob = sapply(1:length(power.ratio),function(i){
          (sum(simulated.ratio[i,]<power.ratio[i])+sum(simulated.ratio[i,]==power.ratio[i])/2)/dim(simulated.ratio)[2]
        })
        return(cumulative.prob)
      })
    })
    power.summary = this.power[[1]]*0
    for(nn in 1:1000){
      power.summary = power.summary + as.numeric(this.power[[nn]]>0.975|this.power[[nn]]<0.025)
    }
    power.summary = power.summary/1000
    return(power.summary)
  })
  names(this.k.power) = names(k.data)
  return(this.k.power)
})

# check the proportion of contrasts that have a jump and compare it to the null hypothesis
jump.ratio = lapply(clado.contrast,function(x){
  sub.pps = pps[,x-Ntip(tree)]
  empirical.ratio = sapply(1:dim(pps)[1],function(i){
    sum(sub.pps[i,]<0.25)/length(x)
  })
  simulated.ratio = sapply(sim.pps,function(y){
    sub.y = y[,x-Ntip(tree)]
    sapply(1:dim(y)[1],function(i){
      sum(sub.y[i,]<0.25)/length(x)
    })
  })
  cumulative.prob = sapply(1:length(empirical.ratio),function(i){
    (sum(simulated.ratio[i,]<empirical.ratio[i])+sum(simulated.ratio[i,]==empirical.ratio[i])/2)/dim(simulated.ratio)[2]
  })
  return(list(empirical=empirical.ratio,sim=simulated.ratio,p.value=cumulative.prob))
})

# save results
saveRDS(list(model=best.model,pps=pps,stat = jump.ratio,power.p = sim.power),
        sprintf("%s.clado.RDS",job.name))
#
warnings()
```
