## Supplementary material for "Microbial genomic trait evolution is dominated by frequent and rare pulsed evolution": Fig_1.R

```
# set up environments
source("fitPE_functions.R")
source("seg_lm.R")
library(ggplot2)
library(ggpubr)
library(e1071)

# the path to result files
job.names = c("../Results/Bacteria_result/bac.gGC",
              "../Results/Bacteria_result/bac.rGC",
              "../Results/Bacteria_result/bac.GS",
              "../Results/Bacteria_result/bac.N")
names(job.names) = c("Genomic GC%","rRNA GC%","Genome size","N-ARSC")

# read in trait value and phylogeny
all.data = lapply(job.names,function(job){
  dat = readRDS(paste0(job,".data.RDS"))
  return(dat)
})

# read in fitting results
all.res = lapply(job.names,function(job){
  res = readRDS(paste0(job,".models.RDS"))
  res.3pe = list()
  res.gamma = list()
  if(file.exists(paste0(job,".3PE.RDS"))) res.3pe = readRDS(paste0(job,".3PE.RDS"))
  if(file.exists(paste0(job,".gamma.RDS"))) res.gamma = readRDS(paste0(job,".gamma.RDS"))
  return(list(scale=res$scale,stat = c(res$stat,res.3pe$stat,res.gamma$stat),
              params = c(res$params,res.3pe$params,res.gamma$params)))
})

# summarize AIC values
all.AIC = lapply(names(all.res),function(nn){
  x = all.res[[nn]]
  if(nn=="rRNA GC%"){
    this.ancs = 
      reconstructAncestralStates(
        phy = all.data[[nn]]$phy,x = all.data[[nn]]$dat,rate = 1,epsilon = 0)
    ss = sum(this.ancs$contrast!=0)
  }else{
    ss = Nnode(all.data[[nn]]$phy)
  }
  normalize.AIC(AIC = x$stat,scale = x$scale,sample.size = ss)
})
names(all.AIC) = names(all.res)
print(lapply(all.AIC,round))

# AIC weights of BM model 
bm.AICw = sapply(all.AIC,function(x){
  this.likelihood = exp(c(BM=unname(min(x[2:6])-x["bme"]),VRG=unname(min(x[2:6])-x["gamma"]),
                          PE=0)/2)
  return(this.likelihood/unname(sum(this.likelihood)))
})
print(bm.AICw)

# summarize all fitted parameters
all.summary = lapply(all.res,function(x){
  lapply(x$params,summarize.model.parameter,scale=x$scale,time=2000)
})

# which one is the best model for each trait?
best.model = sapply(all.AIC,function(x){names(x)[which.min(x)]})
best.model["N-ARSC"] = "pee3"

# and the best parameters
best.summary = lapply(1:length(best.model),function(i){
  mdl = best.model[i]
  all.summary[[i]][[mdl]]
})
names(best.summary) = names(best.model)
print(best.summary)

# how many jumps are expected over the phylogeny
expected.jumps = lapply(1:length(all.summary),function(i){
  best.params =all.summary[[i]][[best.model[i]]]$params
  numPE = length(best.params)/2-1
  total.branch = sum(all.data[[i]]$phy$edge.length)
  exp.jp = total.branch*best.params[(2*(1:numPE)-1)]
  return(exp.jp)
})

# get bootstrapping results
all.bootstrap = lapply(job.names,function(job){
  res = readRDS(paste0(job,".bootstrap.RDS"))
  params.mat = sapply(res$res,function(x){expand.model.parameter(x$params)})
  return(params.mat)
})

# calculate CI
best.CI = lapply(1:length(all.bootstrap),function(i){
  this.params = expand.model.parameter(all.res[[i]]$params[[best.model[i]]])
  numPE = length(this.params)/4-1
  scale.row = rep(1,length(this.params))
  scale.row[c((1:numPE)*2,2*numPE+(1:(numPE+3)))] = all.res[[i]]$scale
  this.bootstrap = all.bootstrap[[i]]
  this.CI = sapply(1:length(this.params),function(j){
    if(this.params[j]<=0){
      this.row = calculateBootstrapCI(stat = this.params[j],bootstrap.stat = this.bootstrap[j,])
    }else{
      if(j>(3*(numPE+1))){
        if(this.params[j]>=1){
          this.row = calculateBootstrapCI(stat = this.params[j],bootstrap.stat = this.bootstrap[j,])
        }else{
          this.row = sigmoid(calculateBootstrapCI(stat = logit(this.params[j]),bootstrap.stat = logit(this.bootstrap[j,])))
        }
      }else{
        this.row = exp(calculateBootstrapCI(stat = log(this.params[j]),bootstrap.stat = log(this.bootstrap[j,])))
      }
    }
    this.row = this.row/scale.row[j]
  })
  return(this.CI)
})
names(best.CI) = names(job.names)
#
# set cumulative probability functions
pfunction = c(bme = "pPEpoislaplace",pee="pPEpoislaplace",pebme = "pPEpoislaplace",
              pee2 = "pPEpois2laplace",pebme2 = "pPEpois2laplace",
              pee3 = "pPEpois3laplace",vrg="pVRgamma")
# calculate conventional PIC with normal epsilon
bm.pic = lapply(1:length(all.data),function(i){
  dat = all.data[[i]]
  bm.params = all.summary[[i]][["bme"]]$params
  p.pic = pseudo.pic(phy = dat$phy,x = dat$dat[dat$phy$tip.label],pfunc = pfunction["bme"],params = bm.params)
  base.ancs = reconstructAncestralStates(phy = dat$phy,dat$dat[dat$phy$tip.label],
                                         rate = 1,epsilon = 0)
  this.ancs = reconstructAncestralStates(phy = dat$phy,dat$dat[dat$phy$tip.label],
                                         rate = total.process.variance(bm.params),epsilon = bm.params[length(bm.params)])
  return(data.frame(ppic=p.pic,pic=this.ancs$contrast/sqrt(this.ancs$lprime),contrast=this.ancs$contrast,
                    l=this.ancs$l,lprime=base.ancs$lprime,lbme = this.ancs$lprime/total.process.variance(bm.params),
                    trait = names(job.names)[i],stringsAsFactors = FALSE))
})
# calculate pseudo-PIC with Laplace epsilon
best.pic = lapply(1:length(all.data),function(i){
  dat = all.data[[i]]
  best.params = all.summary[[i]][[best.model[i]]]$params
  p.pic = pseudo.pic(phy = dat$phy,x = dat$dat[dat$phy$tip.label],pfunc = pfunction[best.model[i]],params = best.params)
  return(data.frame(ppic=p.pic,
                    trait = names(job.names)[i],stringsAsFactors = FALSE))
})
# check excecss kurtosis
bm.kurtosis = sapply(bm.pic,function(x){
  kurtosis(x$pic)
})
print(bm.kurtosis)
best.kurtosis = sapply(best.pic,function(x){
  kurtosis(x$ppic)
})
print(best.kurtosis)

# check normality
bm.ks.test = lapply(bm.pic,function(x){
  ks.test(x$pic,pnorm)
})
print(sapply(bm.ks.test,function(x){x$p.value}))
best.ks.test = lapply(best.pic,function(x){
  ks.test(x$ppic,pnorm)
})

# estimate stasis time
stasis.time = sapply(1:length(best.summary),function(i){
  this.summary = best.summary[[i]]
  this.l = estimate.stasis.period(params = this.summary$params,pfunc = pfunction[best.model[i]])
  names(this.l) = pfunction[best.model[i]]
  return(this.l)
})
names(stasis.time) = names(best.summary)
print(stasis.time*2000)

# BM expectation line
normal.line.df = data.frame(x=seq(-10,10,0.05),
                            density = sapply(seq(-10,10,0.05),function(x){pnorm(x+0.025)-pnorm(x-0.025)})/0.05)
# PE expectation line
fitted.line.df = lapply(1:length(all.data),function(i){
  print(i)
  dat = all.data[[i]]
  bm.params = all.summary[[i]][["bme"]]$params
  this.ancs = reconstructAncestralStates(phy = dat$phy,dat$dat[dat$phy$tip.label],
                                         rate = total.process.variance(bm.params),epsilon = bm.params[length(bm.params)])
  best.params = all.summary[[i]][[best.model[i]]]$params
  best.ancs = reconstructAncestralStates(phy = dat$phy,dat$dat[dat$phy$tip.label],
                                         rate = total.process.variance(best.params),epsilon = best.params[length(best.params)])
  xs = seq(-10,10,0.05)
  x.dp = sapply(seq(-10,10,0.05),function(x){
    cat(sprintf("%f\r",x))
    mean(sapply(seq(1,length(this.ancs$l),100),function(j){
      do.call(pfunction[best.model[i]],
              c(list(q=sqrt(this.ancs$lprime[j])*(x+0.025),t = this.ancs$l[j],epsilon=best.ancs$epsilon[j]),
                as.list(best.params[-length(best.params)])))-
        do.call(pfunction[best.model[i]],
                c(list(q=sqrt(this.ancs$lprime[j])*(x-0.025),t = this.ancs$l[j],epsilon=best.ancs$epsilon[j]),
                  as.list(best.params[-length(best.params)])))
    }))
  })
  data.frame(x=xs,
             density = x.dp/0.05,
             trait = names(job.names)[i],stringsAsFactors = FALSE)
})
fitted.gamma.df = lapply(1:length(all.data),function(i){
  print(i)
  dat = all.data[[i]]
  bm.params = all.summary[[i]][["bme"]]$params
  this.ancs = reconstructAncestralStates(phy = dat$phy,dat$dat[dat$phy$tip.label],
                                         rate = total.process.variance(bm.params),epsilon = bm.params[length(bm.params)])
  gamma.params = all.summary[[i]][["gamma"]]$params
  best.ancs = reconstructAncestralStates(phy = dat$phy,dat$dat[dat$phy$tip.label],
                                         rate = gamma.params["k"]*gamma.params["theta"],epsilon = gamma.params["epsilon"])
  xs = seq(-10,10,0.05)
  x.dp = sapply(seq(-10,10,0.05),function(x){
    cat(sprintf("%f\r",x))
    mean(sapply(seq(1,length(this.ancs$l),100),function(j){
      do.call("pVRgamma",
              c(list(q=sqrt(this.ancs$lprime[j])*(x+0.025),t = this.ancs$l[j],epsilon=best.ancs$epsilon[j]),
                as.list(gamma.params[-length(gamma.params)])))-
        do.call("pVRgamma",
                c(list(q=sqrt(this.ancs$lprime[j])*(x-0.025),t = this.ancs$l[j],epsilon=best.ancs$epsilon[j]),
                  as.list(gamma.params[-length(gamma.params)])))
    }))
  })
  data.frame(x=xs,
             density = x.dp/0.05,
             trait = names(job.names)[i],stringsAsFactors = FALSE)
})

# plot PIC distribution
bm.pic.plot = lapply(1:length(bm.pic),function(i){
  ggplot()+
    geom_histogram(data=bm.pic[[i]],mapping = aes(x=pic,y=..density..,group=trait),binwidth = 0.05)+
    geom_line(data =normal.line.df,mapping = aes(x=x,y=density),color="red",alpha=0.7,size=1)+
    geom_line(data =fitted.line.df[[i]],mapping = aes(x=x,y=density,group=trait),color="green",alpha=0.7,size=1)+
    geom_line(data =fitted.gamma.df[[i]],mapping = aes(x=x,y=density,group=trait),color="blue",alpha=0.7,size=1)+
    coord_cartesian(xlim = c(-5,5))+
    scale_y_continuous(trans = "sqrt",breaks = c(0.05,0.2,0.5,1.0,1.5))+
    scale_x_continuous(breaks=seq(-5,5,1))+
    xlab("PIC")+ylab("Density")+ggtitle(names(job.names)[i])+
    theme(legend.position = "none",
          panel.background = element_blank(),
          axis.line = element_line())
})

# get zero-branch length genome pairs from the full phylogeny
full.tree = multi2di(read.tree("../Data/identical_alignment/bacteria.10616.tre"))
full.dist = cophenetic.phylo(full.tree)
tip.list = 1:Ntip(full.tree)
iden.acc =list()
while(length(tip.list)>0){
  this.iden = which(full.dist[,tip.list[1]]<=0)
  iden.acc[[length(iden.acc)+1]] = this.iden
  tip.list = setdiff(tip.list,this.iden)
}
iden.acc.group = iden.acc[sapply(iden.acc,length)>1]
iden.pairs = do.call(c,lapply(iden.acc.group,function(x){
  lapply(0:floor(length(x)/2-1),function(i){
    names(c(x[2*i+1],x[2*i+2]))
  })
}))
rm(full.dist)

# calculate zero-branch variation
full.trait = lapply(c("../Data/data/genomic_GC.logit.txt",
                      "../Data/data/rRNA_stem_GC.logit.txt",
                      "../Data/data/genome_size.log.txt",
                      "../Data/data/N_ARSC.log.txt"),
       function(f){
         trait = read.table(f,sep="\t",stringsAsFactors = FALSE)[,2]
         names(trait) = read.table(f,sep="\t",stringsAsFactors = FALSE)[,1]
         return(trait)
       })
#
zero.var.df = lapply(1:length(full.trait),function(i){
  trait = full.trait[[i]]
  data.frame(x=sapply(iden.pairs,function(x){
    trait[x[1]]-trait[x[2]]
  }),l=1e-7,trait = names(job.names)[i])
})
#
normal.dx.df = lapply(1:length(all.data),function(i){
  dat = all.data[[i]]
  base.ancs = reconstructAncestralStates(phy = dat$phy,dat$dat[dat$phy$tip.label],
                                         rate = 1,epsilon = 0)
  df = data.frame(x=exp(seq(min(log(base.ancs$lprime)),max(log(base.ancs$lprime)),length.out = 100)),
                  trait =names(job.names)[i],stringsAsFactors = FALSE)
  df$y = sapply(df$x,function(x){
    this.p = function(y){
      0.975-do.call(what = "pPEpoisnorm",args = c(list(q=y,t=x),as.list(all.summary[[i]][["bme"]]$params)))
    }
    this.root = uniroot(f = this.p,interval = c(0,5))
    return(this.root$root)
  })
  df$yfit = sapply(df$x,function(x){
    this.p = function(y){
      0.975-do.call(what = pfunction[best.model[i]],args = c(list(q=y,t=x),as.list(all.summary[[i]][[best.model[i]]]$params)))
    }
    this.root = uniroot(f = this.p,interval = c(0,5))
    return(this.root$root)
  })
  df$ygamma = sapply(df$x,function(x){
    this.p = function(y){
      0.975-do.call(what = "pVRgamma",args = c(list(q=y,t=x),as.list(all.summary[[i]][["gamma"]]$params)))
    }
    this.root = uniroot(f = this.p,interval = c(0,5))
    return(this.root$root)
  })
  return(df)
})

#
all.pseudo.scale = c(250,500,80,125)
contrast.plot = lapply(1:length(bm.pic),function(i){
  pseudo.scale = all.pseudo.scale[i]
  y.break = c(0,0.03,0.1,0.3,1,3)
  y.break = c(-y.break,0,y.break)
  ggplot()+
    geom_point(data = bm.pic[[i]],mapping = aes(x=lprime,y=contrast*pseudo.scale,group=trait),alpha=0.5,shape=".")+
    geom_point(data = zero.var.df[[i]],mapping = aes(x=l,y=x*pseudo.scale,group=trait),color="red",alpha=0.05,shape=".")+
    geom_line(data = normal.dx.df[[i]],mapping = aes(x=x,y=y*pseudo.scale),color="red",size=1,alpha=0.7)+
    geom_line(data = normal.dx.df[[i]],mapping = aes(x=x,y=yfit*pseudo.scale),color="green",size=1,alpha=0.7)+
    geom_line(data = normal.dx.df[[i]],mapping = aes(x=x,y=-y*pseudo.scale),color="red",size=1,alpha=0.7)+
    geom_line(data = normal.dx.df[[i]],mapping = aes(x=x,y=-yfit*pseudo.scale),color="green",size=1,alpha=0.7)+
    geom_line(data = normal.dx.df[[i]],mapping = aes(x=x,y=-ygamma*pseudo.scale),color="blue",size=1,alpha=0.7)+
    geom_line(data = normal.dx.df[[i]],mapping = aes(x=x,y=ygamma*pseudo.scale),color="blue",size=1,alpha=0.7)+
    scale_y_continuous(trans = "pseudo_log",breaks=y.break*pseudo.scale,labels = y.break)+
    scale_x_continuous(trans = "log",breaks = 10^seq(-7,0,1),labels = c(0,expression(10^-6),expression(10^-5),
                                                                        expression(10^-4),expression(10^-3),
                                                                        expression(10^-2),expression(10^-1),
                                                                        expression(10^-0)))+
    xlab("Branch length")+ylab("Trait change")+ggtitle(names(job.names)[i])+
    theme(legend.position = "none",
          panel.background = element_blank(),
          axis.line = element_line())
})
#
pic.contrast.plot = ggarrange(plotlist = c(bm.pic.plot[c(2,1,3,4)],contrast.plot[c(2,1,3,4)]),
                              ncol = 4,nrow = 2,align = "hv",labels = "AUTO")
ggsave(plot = pic.contrast.plot,filename = "Fig_1.pdf",device = "pdf",scale = 1.5,
       width = 8,height = 4.5,units = "in",dpi = "print")
if(Sys.info()["sysname"]=="Windows"){
  ggsave(plot = pic.contrast.plot,filename = "Fig_1.png",device = "png",scale = 1.5,
         width = 8,height = 4.5,units = "in",dpi = "print",type="cairo")
}else{
  ggsave(plot = pic.contrast.plot,filename = "Fig_1.png",device = "png",scale = 1.5,
         width = 8,height = 4.5,units = "in",dpi = "print")
}

# save.image("Fig_1.RData")
```
