## Supplementary material for "Microbial genomic trait evolution is dominated by frequent and rare pulsed evolution": Fig_2.R

```
# set up environments
source("fitPE_functions.R")
library(ggplot2)
library(ggpubr)
library(ggtree)
# the path to result files
job.names = c("../Results/Bacteria_result/bac.gGC","../Results/Bacteria_result/bac.rGC","../Results/Bacteria_result/bac.GS","../Results/Bacteria_result/bac.N")
names(job.names) = c("Genomic GC%","rRNA GC%","Genome size","N-ARSC")
# get phylogeny and trait value
all.data = lapply(job.names,function(job){
  dat = readRDS(paste0(job,".data.RDS"))
  return(dat)
})
# get fitted model
all.res = lapply(job.names,function(job){
  res = readRDS(paste0(job,".models.RDS"))
  res.3pe = readRDS(paste0(job,".3PE.RDS"))
  return(list(scale=res$scale,stat = c(res$stat,res.3pe$stat),params = c(res$params,res.3pe$params)))
})
# extract AIC stats
all.AIC = lapply(all.res,function(x){
  normalize.AIC(AIC = x$stat,scale = x$scale,sample.size = Nnode(all.data[[1]]$phy))
})
# translate model parameters
all.summary = lapply(all.res,function(x){
  lapply(x$params,summarize.model.parameter,scale=x$scale,time=2000)
})
# which one is the best model for each trait?
best.model = sapply(all.AIC,function(x){names(x)[which.min(x)]})
best.model["N-ARSC"] = "pee3"
# how many jumps are expected over the phylogeny
best.params = lapply(1:length(best.model),function(i){
  all.summary[[i]][[best.model[i]]]$params
})
#
postfunction = c(bme = "posteriorJumpProbability",pee="posteriorJumpProbability",pebme = "posteriorJumpProbability",
               pee2 = "posteriorJumpProbability.laplace2",pebme2 = "posteriorJumpProbability.laplace2",
               pee3 = "posteriorJumpProbability.laplace3")
pps.row = c(bme = 1,pee=1,pebme = 1,
            pee2 = 2,pebme2 = 2,
            pee3 = 7)
#
post.prob = lapply(1:length(all.data),function(i){
  this.trait = all.data[[i]]$dat
  this.tree = all.data[[i]]$phy
  this.anc = reconstructAncestralStates(phy = this.tree,x = this.trait[this.tree$tip.label],
                                        rate = total.process.variance(best.params[[i]]),
                                        epsilon = best.params[[i]][length(best.params[[i]])])
  pps = sapply(1:(Ntip(this.tree)+Nnode(this.tree)),function(j){
    this.parent = getLatestAncestor(phy = this.tree,j)
    if(is.na(this.parent)) return(do.call(postfunction[best.model[i]],
                                   c(list(x=0,t=10,epsilon=1),best.params[[i]][-length(best.params[[i]])])))
    this.dx = this.anc$trait[j]-this.anc$trait[this.parent]
    this.l = this.tree$edge.length[this.tree$edge[,2]==j]
    this.epsilon = this.anc$var[j]
    this.pp = do.call(postfunction[best.model[i]],
                      c(list(x=this.dx,t=this.l,epsilon=this.epsilon),best.params[[i]][-length(best.params[[i]])]))
    return(this.pp)
  })
  return(pps)
})
#
tree = all.data[[1]]$phy
taxids = readRDS("../Data/taxonomy/acc2taxid.RDS")
taxlin = readRDS("../Data/taxonomy/spp.tax.df.RDS")
tip.phyla = as.character(taxlin$phylum[match(taxids[tree$tip.label],taxlin$taxid)])
tip.phyla[is.na(tip.phyla)] = "Unclassified"
#
attr(tree,"phyla") = sapply(1:(Ntip(tree)+tree$Nnode),function(j){
  if(j<=Ntip(tree)){
    if(is.na(tip.phyla[j])) return("Unclassified")
    return(tip.phyla[j])
  }
  subclade = extract.clade(tree,j)
  clade.phylum = unique(tip.phyla[match(subclade$tip.label,tree$tip.label)])
  if(length(clade.phylum)>1) return("Mixed")
  return(clade.phylum)
})
major.clade = names(sort(table(tip.phyla),decreasing = TRUE))[1:6]
#
maxMono.node = numeric(0)
clade.name = character(0)
node2search = Ntip(tree)+1
while(length(node2search)>0){
  this.node = node2search[1]
  node2search=node2search[-1]
  if(this.node<=Ntip(tree)){
    next
  }
  if(attr(tree,"phyla")[this.node]!="Mixed"&attr(tree,"phyla")[this.node]!="Unclassified"){
    maxMono.node = c(maxMono.node,this.node)
    this.name = attr(tree,"phyla")[this.node]
    if((this.name %in% clade.name)|!(this.name %in% major.clade)){
      clade.name = c(clade.name,"")
    }else{
      clade.name = c(clade.name,this.name)
    }
  }else{
    node2search = c(node2search,tree$edge[tree$edge[,1]==this.node,2])
  }
}
colormap = rep("black",(length(unique(attr(tree,"phyla")))))
names(colormap) = unique(attr(tree,"phyla"))
colormap[names(sort(table(tip.phyla),decreasing = TRUE))[1:6]] = c("steelblue2","purple1","orange1","magenta3","seagreen3","cyan2")
#
gc.phy.plot = list()
trees = list()
for(i in 1:length(post.prob)){
  this.ppj = post.prob[[i]][pps.row[best.model[i]],]
  this.ppj = as.character(as.numeric(this.ppj<0.05))
  trees[[i]] = tree
  attr(trees[[i]],which = "ppj") = this.ppj
  gc.phy.plot[[i]] = ggtree(tr = trees[[i]],mapping = aes(color=ppj,size=ppj),layout='circular',branch.length = 1)+
    scale_color_manual(values = c("1"="red","0"="black"))+
    scale_size_manual(values = c("1"=0.25,"0"=0.1))+
    ggtitle(names(job.names)[i])+
    theme(legend.position="bottom")
  for(n in 1:length(maxMono.node)){
    gc.phy.plot[[i]] = gc.phy.plot[[i]]+geom_cladelabel(label=clade.name[n],node = maxMono.node[n],offset = 0,geom = "text",align = TRUE,barsize = 1,offset.text = 0.5,fontsize = 4,color=colormap[attr(tree,"phyla")[maxMono.node[n]]])
  }
}
all.phy.plot = ggarrange(plotlist = gc.phy.plot[c(2,1,3,4)],ncol = 2,nrow = 2,labels = "AUTO",align = "hv",common.legend = TRUE,legend = "bottom")
```
