## Supplementary material for "Microbial genomic trait evolution is dominated by frequent and rare pulsed evolution": Fig_S1.R

```
#
trait.cor = readRDS("../Results/Bacteria_result/jump.cor.test.cor.RDS")
jump.cor = readRDS("../Results/Bacteria_result/jump.cor.test.jump.cor.RDS")
#
tree = read.tree("../Data/kingdom/combo.cds.HQ.6668.tre")
#
all.comb = combn(x = length(trait.cor$plot),m = 2)
tip.pairs = table(tree$edge[tree$edge[,2]<=Ntip(tree),1])
tip.pairs = as.numeric(names(tip.pairs[tip.pairs>1]))
#
plot.grid = lapply(1:length(trait.cor$plot),function(i){
  lapply(1:length(trait.cor$plot),function(x){list()})
})#
library(ggplot2)
library(ggpubr)
#
trait.names = c("rRNA GC%","Genomic GC%","Genome size","N-ARSC")
trait.labels = lapply(trait.names,function(x){
  ggplot()+annotate(geom = "text",x = 0,y = 0,label=x,angle=0,fontface="bold")+
    theme(legend.position = "none",
          panel.background = element_blank(),
          axis.title = element_blank(),
          axis.line = element_blank(),
          axis.text = element_blank(),
          axis.ticks = element_blank())
})
#
place.holder =  ggplot()+
  theme(legend.position = "none",
        panel.background = element_blank(),
        axis.title = element_blank(),
        axis.line = element_blank(),
        axis.text = element_blank(),
        axis.ticks = element_blank())
#
pic.plots = apply(all.comb,2,function(i){
  this.df = trait.cor$pic$pic[tip.pairs-Ntip(tree),i]
  colnames(this.df) = c("x","y")
  this.test = cor.test(this.df$x,this.df$y)
  this.lm = lm
  if(this.test$p.value<0.001){
    this.label = sprintf("%.3f\n(P<0.001)",this.test$estimate^2)
  }else{
    this.label = sprintf("%.3f\n(P=%.3f)",this.test$estimate^2,this.test$p.value)
  }
  mark.x = 0.75*max(this.df$y)+0.25*min(this.df$y)
  mark.y = 0.05*max(this.df$x)+0.95*min(this.df$x)
  ggplot()+
    geom_point(mapping = aes(x=y,y=x),data = this.df,shape=".",alpha=0.8)+
    annotate(geom = "text",x = mark.x,y = mark.y,label=this.label,angle=0,size=2,color="red")+
    xlab("PIC")+ylab("PIC")+
    theme(legend.position = "none",
          panel.background = element_blank(),
          axis.title = element_text(size = 8),
          axis.line = element_line())
})

pic.cor.data = apply(all.comb,2,function(i){
  this.df = trait.cor$pic$pic[tip.pairs-Ntip(tree),i]
  colnames(this.df) = c("x","y")
  this.test = cor.test(this.df$x,this.df$y)
  this.lm = lm(formula = x~y,data = this.df)
  return(list(cor=this.test,lm=this.lm))
})

ppic.plots = apply(all.comb,2,function(i){
  this.df = trait.cor$pic$ppic[tip.pairs-Ntip(tree),i]
  colnames(this.df) = c("x","y")
  this.test = cor.test(this.df$x,this.df$y)
  this.lm = lm
  if(this.test$p.value<0.001){
    this.label = sprintf("%.3f\n(P<0.001)",this.test$estimate^2)
  }else{
    this.label = sprintf("%.3f\n(P=%.3f)",this.test$estimate^2,this.test$p.value)
  }
  mark.x = 0.75*max(this.df$x)+0.25*min(this.df$x)
  mark.y = min(this.df$y)
  ggplot()+
    geom_point(mapping = aes(x=x,y=y),data = this.df,shape=".",alpha=0.8)+
    annotate(geom = "text",x = mark.x,y = mark.y,label=this.label,angle=0,size=2,color="red")+
    scale_x_continuous(expand = c(0.1,0))+
    scale_y_continuous(expand = c(0.1,0))+
    xlab("Pseudo-PIC")+ylab("Pseudo-PIC")+
    theme(legend.position = "none",
          panel.background = element_blank(),
          axis.title = element_text(size = 8),
          axis.line = element_line())
})

for(i in 1:length(trait.cor$plot)){
  plot.grid[[i]][[i]] = trait.labels[[i]]
}
for(i in 1:dim(all.comb)[2]){
  plot.grid[[all.comb[1,i]]][[all.comb[2,i]]] = pic.plots[[i]]
  plot.grid[[all.comb[2,i]]][[all.comb[1,i]]] = ppic.plots[[i]]
}
combine.plot = ggarrange(plotlist = do.call(c,plot.grid),ncol = length(trait.cor$plot),nrow = length(trait.cor$plot),align = "hv")
if(Sys.info()["sysname"]=="Windows"){
  ggsave(filename = paste0("Fig_S1.png"),plot = combine.plot,device = "png",scale = 4.5,
         width = length(trait.cor$plot),height = length(trait.cor$plot),units = "cm",type="cairo")
}else{
  ggsave(filename = paste0("Fig_S1.png"),plot = combine.plot,device = "png",scale = 4.5,
         width = length(trait.cor$plot),height = length(trait.cor$plot),units = "cm")
  
}
ggsave(filename = paste0("Fig_S1.pdf"),plot = combine.plot,device = "pdf",scale = 4.5,
       width = length(trait.cor$plot),height = length(trait.cor$plot),units = "cm")
```
