## Supplementary material for "Microbial genomic trait evolution is dominated by frequent and rare pulsed evolution": Fig_S3S4.R

```
# set up environments
source("fitPE_functions.R")
source("seg_lm.R")
library(ggplot2)
library(ggpubr)
library(e1071)
# the path to result files
job.names = c("../Results/Bacteria_result/bac.gGC",
              "../Results/Bacteria_result/bac.rGC",
              "../Results/Bacteria_result/bac.GS",
              "../Results/Bacteria_result/bac.N")
names(job.names) = c("Genomic GC%","rRNA GC%","Genome size","N-ARSC")
# read in trait value and phylogeny
all.data = lapply(job.names,function(job){
  dat = readRDS(paste0(job,".data.RDS"))
  return(dat)
})
# read in results
all.res = lapply(job.names,function(job){
  res = readRDS(paste0(job,".models.RDS"))
  res.3pe = list()
  if(file.exists(paste0(job,".3PE.RDS"))) res.3pe = readRDS(paste0(job,".3PE.RDS"))
  return(list(scale=res$scale,stat = c(res$stat,res.3pe$stat),params = c(res$params,res.3pe$params)))
})
gamma.res = lapply(job.names,function(job){
  res = readRDS(paste0(job,".gamma.RDS"))
  return(res)
})

# summarize all fitted parameters
all.summary = lapply(all.res,function(x){
  lapply(x$params,summarize.model.parameter,scale=x$scale,time=2000)
})

# set cumulative probability functions
pfunction = c(bme = "pPEpoislaplace",pee="pPEpoislaplace",pebme = "pPEpoislaplace",
              pee2 = "pPEpois2laplace",pebme2 = "pPEpois2laplace",
              pee3 = "pPEpois3laplace",vrg="pVRgamma")
# calculate conventional PIC with normal epsilon
bm.pic = lapply(1:length(all.data),function(i){
  dat = all.data[[i]]
  bm.params = all.summary[[i]][["bme"]]$params
  p.pic = pseudo.pic(phy = dat$phy,x = dat$dat[dat$phy$tip.label],pfunc = pfunction["bme"],params = bm.params)
  base.ancs = reconstructAncestralStates(phy = dat$phy,dat$dat[dat$phy$tip.label],
                                         rate = 1,epsilon = 0)
  this.ancs = reconstructAncestralStates(phy = dat$phy,dat$dat[dat$phy$tip.label],
                                         rate = total.process.variance(bm.params),epsilon = bm.params[length(bm.params)])
  return(data.frame(ppic=p.pic,pic=this.ancs$contrast/sqrt(this.ancs$lprime),contrast=this.ancs$contrast,
                    l=this.ancs$l,lprime=base.ancs$lprime,lbme = this.ancs$lprime/total.process.variance(bm.params),
                    trait = names(job.names)[i],stringsAsFactors = FALSE))
})

# get zero-branch length genome pairs from the full phylogeny
full.tree = multi2di(read.tree("../Data/identical_alignment/bacteria.10616.tre"))
full.dist = cophenetic.phylo(full.tree)
tip.list = 1:Ntip(full.tree)
iden.acc =list()
while(length(tip.list)>0){
  this.iden = which(full.dist[,tip.list[1]]<=0)
  iden.acc[[length(iden.acc)+1]] = this.iden
  tip.list = setdiff(tip.list,this.iden)
}
iden.acc.group = iden.acc[sapply(iden.acc,length)>1]
iden.pairs = do.call(c,lapply(iden.acc.group,function(x){
  lapply(0:floor(length(x)/2-1),function(i){
    names(c(x[2*i+1],x[2*i+2]))
  })
}))
rm(full.dist)

# calculate zero-branch variation
full.trait = lapply(c("../Data/data/genomic_GC.logit.txt",
                      "../Data/data/rRNA_stem_GC.logit.txt",
                      "../Data/data/genome_size.log.txt",
                      "../Data/data/N_ARSC.log.txt"),
                    function(f){
                      trait = read.table(f,sep="\t",stringsAsFactors = FALSE)[,2]
                      names(trait) = read.table(f,sep="\t",stringsAsFactors = FALSE)[,1]
                      return(trait)
                    })
#
zero.var.df = lapply(1:length(full.trait),function(i){
  trait = full.trait[[i]]
  data.frame(x=sapply(iden.pairs,function(x){
    trait[x[1]]-trait[x[2]]
  }),l=1e-7,trait = names(job.names)[i])
})

# segmented linear regression
lm.data = lapply(1:length(all.summary),function(i){
  base.dat = reconstructAncestralStates(phy = all.data[[i]]$phy,x = all.data[[i]]$dat[all.data[[i]]$phy$tip.label],
                                   rate = 1,epsilon = 0)
  dat = reconstructAncestralStates(phy = all.data[[i]]$phy,x = all.data[[i]]$dat[all.data[[i]]$phy$tip.label],
                                   rate = total.process.variance(all.summary[[i]]$bme$params),
                                   epsilon = all.summary[[i]]$bme$params[length(all.summary[[i]]$bme$params)])
  df = data.frame(x= c(dat$contrast,zero.var.df[[i]]$x),
                  l=c(base.dat$lprime,
                      rep(1e-7,length(zero.var.df[[i]]$x))),
                  shape="dot",color="black")
  return(df)
})
l1.aic.fix = lapply(lm.data,function(dat){
  best.1.model = find.best.1.break.fix(l = dat$l,dat$x)
  best.2.model = find.best.2.break.fix(l = dat$l,dat$x)
  return(list(best1=best.1.model,best2 = best.2.model))
})
#
print(c(l1.aic.fix[[2]]$best2$l1,l1.aic.fix[[2]]$best2$l2))
print(c(l1.aic.fix[[2]]$best2$l1,l1.aic.fix[[2]]$best2$l2)*2000)
#
lm.plot = lapply(1:length(lm.data),function(i){
  dat = l1.aic.fix[[i]]
  df = data.frame(l=lm.data[[i]]$l,dx = abs(lm.data[[i]]$x)+0.001)
  best.aic = sapply(dat,function(x){x$aic})
  if(which.min(best.aic)>1){
    lm.line.df = data.frame(x=c(min(df$l),dat[[2]]$l1,dat[[2]]$l2),
                            y=exp(c(dat[[2]]$model$y0+log(min(df$l))*dat[[2]]$model$k1,dat[[2]]$model$y1,dat[[2]]$model$y2)),
                            xend=c(dat[[2]]$l1,dat[[2]]$l2,max(df$l)),
                            yend=exp(c(dat[[2]]$model$y1,dat[[2]]$model$y2,(log(max(df$l))-log(dat[[2]]$l2))*dat[[2]]$model$k3+dat[[2]]$model$y2)),
                            color=1:3)
  }else{
    lm.line.df = data.frame(x=c(min(df$l),dat[[1]]$l1),
                            y=exp(c(dat[[1]]$model$y0+log(min(df$l))*dat[[2]]$model$k1,dat[[1]]$model$y1)),
                            xend=c(dat[[1]]$l1,max(df$l)),
                            yend=exp(c(dat[[1]]$model$y1,(log(max(df$l))-log(dat[[1]]$l1))*dat[[1]]$model$k2+dat[[1]]$model$y1)),
                            color=1:2)
  }
  #print(lm.line.df)
  vline.df = data.frame(x = lm.line.df$x[-1])
  ggplot()+
    geom_point(mapping = aes(x=l,y=dx),data = df,alpha=0.75,shape=".")+
    geom_segment(mapping = aes(x=x,y=y,xend=xend,yend=yend),data = lm.line.df,alpha=0.8,color="red",size=1)+
    geom_vline(mapping = aes(xintercept = x),data = vline.df,alpha=0.8,color="red",linetype="dashed")+
    scale_x_continuous(trans = "log10",breaks=c(1e-7,10^seq(-6,0,1)),
                       labels = c(0,expression(10^-6),expression(10^-5),
                                  expression(10^-4),expression(10^-3),
                                  expression(10^-2),expression(10^-1),
                                  expression(10^-0)))+
    scale_y_continuous(trans = "log10",breaks=10^seq(-3,0))+
    xlab("Branch length")+ylab("Absolute trait change")+ggtitle(names(job.names)[i])+
    theme(legend.position = "none",
          panel.background = element_blank(),
          axis.line = element_line())
})
#
lm.combined.plot = ggarrange(plotlist = lm.plot[c(2,1,3,4)],ncol = 2,nrow = 2,align = "hv",labels = "AUTO")
ggsave(plot = lm.plot[[2]],filename = "Fig_S4.pdf",device = "pdf",scale = 1.5,
       width = 4,height = 3,units = "in",dpi = "print")
if(Sys.info()["sysname"]=="Windows"){
  ggsave(plot = lm.plot[[2]],filename = "Fig_S4.png",device = "png",scale = 1.5,
       width = 4,height = 3,units = "in",dpi = "print",type="cairo")
}else{
  ggsave(plot = lm.plot[[2]],filename = "Fig_S4.png",device = "png",scale = 1.5,
         width = 4,height = 3,units = "in",dpi = "print")
}

# generate PIC-Time plot
rela.dist.root = readRDS("../Data/kingdom/relative.dist.root.RDS")
bins= c(0,1-0.3^(1:6),1)
midpoints = (bins[-1]+bins[-length(bins)])/2
short.branch = bm.pic[[1]]$l<1e-5
df.count = lapply(1:length(bm.pic),function(i){
  dff = bm.pic[[i]]
  r2r = rela.dist.root[-(0:length(dff$contrast)+1)]
  sapply(2:length(bins)-1,function(i){
    idx = which(!short.branch&r2r>=bins[i]&r2r<bins[i+1])
    pppic = dff$pic[idx]
    c1 = sum(abs(pppic)>3)
    c2 = length(pppic)-c1
    return(c(c1,c2))
  })
})
df.count.test = lapply(df.count,function(dff){
  apply(dff,2,function(x){
    binom.test(x = x,p = 2*pnorm(q = -3),conf.level = 0.95)
  })
})
#
df.count.data = lapply(1:length(df.count.test),function(i){
  dff = df.count.test[[i]]
  data.frame(d=midpoints,
             pseudo.d = -log10((1-midpoints)),
             p = exp(0.25*log(unname(sapply(dff,function(x){x$estimate})))),
             uci = exp(0.25*log(unname(sapply(dff,function(x){x$conf.int[2]})))),
             lci = exp(0.25*log(unname(sapply(dff,function(x){x$conf.int[1]})))),
             trait = names(job.names)[i],
             stringsAsFactors = FALSE)
})
freq.plot = lapply(1:length(bm.pic),function(i){
  ggplot()+
    geom_line(mapping = aes(x=pseudo.d,y=p),color="black",data=df.count.data[[i]],position = position_dodge(0.1),linetype="dashed")+
    geom_point(mapping = aes(x=pseudo.d,y=p,shape=trait),data=df.count.data[[i]],position = position_dodge(0.1))+
    geom_errorbar(mapping = aes(x=pseudo.d,group=trait,ymin=lci,ymax=uci),data=df.count.data[[i]],width=0.1)+
    geom_hline(yintercept = exp(0.25*log(2*pnorm(-3))),color="red",linetype="dashed")+
    scale_y_continuous(expand=c(0.01,0),breaks=exp(0.25*log(c(0,0.0001,0.003,0.01,0.025,0.05,0.1,0.2,0.3))),labels=c(0,0.01,0.3,1,2.5,5,10,20,30))+
    scale_x_continuous(breaks=-log10(10^seq(0,-3,-1)),labels = 1-10^seq(0,-3,-1))+
    xlab("Relative distance from the root")+
    ylab("% of extreme trait change")+
    ggtitle(names(job.names)[i])+
    theme(legend.position = "none",
          panel.background = element_blank(),
          axis.line = element_line())
})

#
df.dat = lapply(1:length(bm.pic),function(i){
  dff = bm.pic[[i]]
  r2r = rela.dist.root[-(0:length(dff$contrast)+1)]
  dtf = data.frame(pic=dff$pic,l=dff$l,d=r2r,pseudo.d=-log10((1-r2r)),color=as.character(abs(dff$pic)>3))
  return(dtf)
})
#
pic.plot = lapply(1:length(df.dat),function(i){
  dff = df.dat[[i]]
  ggplot()+
    geom_point(mapping = aes(x=pseudo.d,y=pic,color=color,alpha=color),shape=".",data = dff)+
    geom_hline(yintercept = 3,color="red",alpha=0.9,linetype="dashed")+
    geom_hline(yintercept = -3,color="red",alpha=0.9,linetype="dashed")+
    scale_y_continuous(trans="pseudo_log",breaks=c(-15,-10,-6,-3,-1,0,1,3,6,10,15),
                       labels = c(-15,-10,-6,-3,-1,0,1,3,6,10,15))+
    scale_x_continuous(breaks=-log10(10^seq(0,-5,-1)),labels = 1-10^seq(0,-5,-1))+
    scale_color_manual(values = c("TRUE"="red","FALSE"="black"))+
    scale_alpha_manual(values = c("TRUE"=1,"FALSE"=0.5))+
    xlab("Relative distance from the root")+
    ylab("PIC")+
    ggtitle(names(job.names)[i])+
    theme(legend.position = "None",
          panel.background = element_blank(),
          axis.line = element_line())
})
#
combine.plot = ggarrange(plotlist = c(pic.plot[c(2,1,3,4)],freq.plot[c(2,1,3,4)]),ncol=4,nrow=2,align = "hv",labels = c("AUTO"))
ggsave(plot = combine.plot,filename = "Fig_S3.pdf",device = "pdf",scale = 1.5,
       width = 8,height = 4.5,units = "in",dpi = "print")
if(Sys.info()["sysname"]=="Windows"){
  ggsave(plot = combine.plot,filename = "Fig_S3.png",device = "png",scale = 1.5,
         width = 8,height = 4.5,units = "in",dpi = "print",type="cairo")
}else{
  ggsave(plot = combine.plot,filename = "Fig_S3.png",device = "png",scale = 1.5,
         width = 8,height = 4.5,units = "in",dpi = "print")
}
#
```
