## Supplementary material for "Microbial genomic trait evolution is dominated by frequent and rare pulsed evolution": Fig_S5.R

```
#
source("fitPE_functions.R")
library(ggplot2)
library(ggpubr)
#
l = 10^(seq(-4,0,0.2))
#
pe1.res = readRDS("../Results/simulation_result/sim.PE1.supp.RDS")
#
pe1.df = data.frame(l=l,
                    prob=sapply(pe1.res,function(x){
                      sum(apply(x,2,which.min)>1)/dim(x)[2]
                    }))
#
pe1.plot = ggplot()+
  geom_line(data=pe1.df,mapping = aes(x=l,y=prob))+
  geom_point(data=pe1.df,mapping = aes(x=l,y=prob))+
  scale_x_continuous(trans="log10",breaks=10^(seq(-4,1,1)),
                     labels = sapply(seq(-4,1,1),function(x){
                       parse(text = sprintf("10^%d",x))
                       }))+
  xlab("Branch length")+ylab("Frequency PE1 as the best model")+
  theme(panel.background = element_blank(),
        axis.line = element_line())
if(Sys.info()["sysname"]=="Windows"){
  ggsave(plot = pe1.plot,filename = "Fig_S5.png",device = "png",scale = 1.75,
         width = 4,height = 3,units = "in",dpi = "print",type="cairo")
}else{
  ggsave(plot = pe1.plot,filename = "Fig_S5.png",device = "png",scale = 1.75,
         width = 4,height = 3,units = "in",dpi = "print")
}
ggsave(plot = pe1.plot,filename = "Fig_S5.pdf",device = "pdf",scale = 1.75,
       width = 4,height = 3,units = "in")

#
```
