## Supplementary material for "Microbial genomic trait evolution is dominated by frequent and rare pulsed evolution": Fig_S6.R

```
# validate jump detection
source("fitPE_functions.R")
sim.truth = lapply(1:10,function(i){readRDS(sprintf("../Results/simulation_result/Sim_PE2_%d.RDS",i))})
#
sim.data = lapply(1:10,function(i){readRDS(sprintf("../Results/simulation_result/Sim_PE2_%d.data.RDS",i))})
sim.res = lapply(1:10,function(i){readRDS(sprintf("../Results/simulation_result/Sim_PE2_%d.PE2.RDS",i))})
bayes.res = lapply(1:10,function(i){read.tree(sprintf("../Results/BayesTraits_result/TIV.Sim_PE2_%d_BayesTrait.rescaled.tre",i))})
bayes.initial = read.nexus("../Results/BayesTraits_result/bac.tiv.sim.nexus")
#
best.model = sapply(sim.res,function(x){which.min(x$stat)})
#
res.params = lapply(sim.res,function(x){
  lapply(x$params,function(xx){
    summarize.model.parameter(params = xx,scale = x$scale)$params   
    })
  })
#
best.ancs = lapply(1:10,function(i){
  best.params = res.params[[i]][[names(best.model[i])]]
  reconstructAncestralStates(phy = sim.data[[i]]$phy,x = sim.data[[i]]$dat[sim.data[[i]]$phy$tip.label],
                             rate = unname(total.process.variance(best.params)),
                             epsilon = unname(best.params["epsilon"]))
})
bayes.ancs = lapply(1:10,function(i){
  reconstructAncestralStates(phy = bayes.res[[i]],x = sim.data[[i]]$dat[bayes.res[[i]]$tip.label],
                             rate = 1,
                             epsilon = 0)
})
# setting functions to use
model.functions = c(bme="posteriorJumpProbability",pee = "posteriorJumpProbability",pebme ="posteriorJumpProbability",
                    pee2= "posteriorJumpProbability.laplace2",pebme2="posteriorJumpProbability.laplace2",pee3 = "posteriorJumpProbability.laplace3")
# calculate prior probability of jump
prs = lapply(1:10,function(j){
  best.params = res.params[[j]][[names(best.model[j])]]
  lambdas = best.params[seq(from = 1,to = length(best.params)-2,by = 2)]
  do.call(cbind,lapply(1:length(best.ancs[[j]]$contrast),function(i){
    pp = dpois(x = 0,lambda = lambdas*best.ancs[[j]]$l[i])
    names(pp) = c("p01","p02")
    return(pp)
  }))
})
# calculate posterior probability of jump
pps = lapply(1:10,function(j){
  best.params = res.params[[j]][[names(best.model[j])]]
  do.call(cbind,lapply(1:length(best.ancs[[j]]$contrast),function(i){
    do.call(model.functions[names(best.model[j])],
            c(list(x=best.ancs[[j]]$contrast[i],t=best.ancs[[j]]$l[i],epsilon=best.ancs[[j]]$epsilon[i]),
              as.list(best.params[-length(best.params)])))
  }))
})
# check the true number of jumps per contrast
sim.jump1.contrast = lapply(1:10,function(j){
  x= sim.data[[j]]
  sapply((1:Nnode(x$phy))+Ntip(x$phy),
         function(i){sum(sim.truth[[j]]$nj1[x$phy$edge[x$phy$edge[,1]==i,2]])})
})
sim.jump2.contrast = lapply(1:10,function(j){
  x= sim.data[[j]]
  sapply((1:Nnode(x$phy))+Ntip(x$phy),
         function(i){sum(sim.truth[[j]]$nj2[x$phy$edge[x$phy$edge[,1]==i,2]])})
})
# simplify to jump or no jump
sim.jump1.binary = lapply(sim.jump1.contrast,function(x){x>0})
sim.jump2.binary = lapply(sim.jump2.contrast,function(x){x>0})
# predict jump by posterior probability
pp.threshold = 0.1
pp.jump1.binary = lapply(pps,function(x){x["p01",]<pp.threshold})
pp.jump2.binary = lapply(pps,function(x){x["p02",]<pp.threshold})
# predict jump by prior probability
pr.jump1.binary = lapply(prs,function(x){x["p01",]>=(1-pp.threshold)})
pr.jump2.binary = lapply(prs,function(x){x["p02",]>=(1-pp.threshold)})
#
short.jump1.candidate = lapply(1:10,function(i){
  sim.jump1.binary[[i]]&pr.jump1.binary[[i]]
})
short.jump2.candidate = lapply(1:10,function(i){
  sim.jump2.binary[[i]]&pr.jump2.binary[[i]]
})
#
short.jump1.recovery = sapply(1:10,function(i){
  sum(short.jump1.candidate[[i]]&pp.jump1.binary[[i]])/sum(short.jump1.candidate[[i]])
})
short.jump2.recovery = sapply(1:10,function(i){
  sum(short.jump2.candidate[[i]]&pp.jump2.binary[[i]])/sum(short.jump2.candidate[[i]])
})
#
pp.jump1.cm = lapply(1:10,function(i){table(pred=pp.jump1.binary[[i]],obs=sim.jump1.binary[[i]])})
pp.jump2.cm = lapply(1:10,function(i){table(pred=pp.jump2.binary[[i]],obs=sim.jump2.binary[[i]])})
#
pp.jump1.sensitivity = sapply(pp.jump1.cm,function(x){
  c(sensitivity=x[2,2]/sum(x[,2]),specificity=x[1,1]/sum(x[,1]))
})
pp.jump2.sensitivity = sapply(pp.jump2.cm,function(x){
  c(sensitivity=x[2,2]/sum(x[,2]),specificity=x[1,1]/sum(x[,1]))
})
#
bayes.scale = lapply(1:10,function(i){
  bm.ancs = reconstructAncestralStates(phy = bayes.initial,x = sim.data[[i]]$dat,rate = 1,epsilon = 0)
  this.scale = bayes.ancs[[i]]$l/bm.ancs$l
  this.scale = this.scale/median(this.scale)
})

bayes.roc.data = do.call(rbind,c(lapply(1:10,function(i){
  this.cutoff = seq(0,1,0.01)
  this.pps = pps[[i]]
  this.jump1 = sim.jump1.binary[[i]]
  this.jump2 = sim.jump2.binary[[i]]
  data.frame(scale.cutoff = c(this.cutoff,this.cutoff),
             sensitivity=c(sapply(this.cutoff,function(x){
               sum(this.pps["p01",this.jump1]<x)/sum(this.jump1)
             }),sapply(this.cutoff,function(x){
               sum(this.pps["p02",this.jump2]<x)/sum(this.jump2)
             })),
             specificity = c(sapply(this.cutoff,function(x){
               1-sum(this.pps["p01",!this.jump1]<x)/sum(!this.jump1)
             }),sapply(this.cutoff,function(x){
               1-sum(this.pps["p02",!this.jump2]<x)/sum(!this.jump2)
             })),
             replicate=rep(c(paste0("PE_rare",i),paste0("PE_freq",i)),each=length(this.cutoff)),
             method=rep(c("Pulsed Evolution (rare)","Pulsed Evolution (frequent)"),
                        each=length(this.cutoff)))
}),
                                 lapply(1:10,function(i){
  this.cutoff = quantile(bayes.scale[[i]],probs=seq(0,1,0.01))
  this.scale = bayes.scale[[i]]
  this.jump1 = sim.jump1.binary[[i]]
  this.jump2 = sim.jump2.binary[[i]]
  data.frame(scale.cutoff = c(this.cutoff,this.cutoff),
             sensitivity=c(sapply(this.cutoff,function(x){
               sum(this.scale[this.jump1]>x)/sum(this.jump1)
             }),sapply(this.cutoff,function(x){
               sum(this.scale[this.jump2]>x)/sum(this.jump2)
             })),
             specificity = c(sapply(this.cutoff,function(x){
               1-sum(this.scale[!this.jump1]>x)/sum(!this.jump1)
             }),sapply(this.cutoff,function(x){
               1-sum(this.scale[!this.jump2]>x)/sum(!this.jump2)
             })),
             replicate=rep(c(paste0("BT_rare",i),paste0("BT_freq",i)),each=length(this.cutoff)),
             method=rep(c("BayesTraits (rare)","BayesTraits (frequent)"),each=length(this.cutoff)))
  })))

roc.plot = ggplot(data=bayes.roc.data)+
  geom_line(mapping = aes(x=specificity,y=sensitivity,color=method,group=replicate),alpha=0.75)+
  geom_abline(slope = 1,intercept = 1,color="red",linetype="dashed")+
  scale_color_manual(values = c("Pulsed Evolution (rare)"="darkgreen",
                                "Pulsed Evolution (frequent)"="green",
                                "BayesTraits (rare)"="blue",
                                "BayesTraits (frequent)"="cyan"),name="Method (Jump type)")+
  coord_cartesian(expand = FALSE)+
  scale_x_reverse()+
  xlab("Specificity")+ylab("Sensitivity")+
  theme(legend.position = c(0.75,0.2),
        panel.background = element_blank(),
        plot.margin = unit(c(0.2,0.2,0.2,0.2),"in"),
        axis.line = element_line())
if(Sys.info()["sysname"]=="Windows"){
  ggsave(plot = roc.plot,filename = "Fig_S6.png",device = "png",scale = 1.75,
         width = 4,height = 4,units = "in",dpi = "print",type="cairo")
}else{
  ggsave(plot = roc.plot,filename = "Fig_S6.png",device = "png",scale = 1.75,
         width = 4,height = 4,units = "in",dpi = "print")
}
ggsave(plot = roc.plot,filename = "Fig_S6.pdf",device = "pdf",scale = 1.75,
       width = 4,height = 4,units = "in")

auc = function(x,y){
  xidx = sort(x,index.return=TRUE)$ix
  x = x[xidx]
  y = y[xidx]
  xs = x[-1]-x[-length(x)]
  ys = (y[-length(y)]+y[-1])/2
  sum(xs*ys)
}

aucs = sapply(unique(bayes.roc.data$replicate),function(x){
  this.row = bayes.roc.data$replicate==x
  auc(y = bayes.roc.data$sensitivity[this.row],x = bayes.roc.data$specificity[this.row])
})
  
  
  
```
