## Supplementary material for "Microbial genomic trait evolution is dominated by frequent and rare pulsed evolution": fit_3PE_model.R

```
# read in key arguments for the fitting
arg=commandArgs(TRUE)
fullarg = commandArgs(FALSE)
# setup the environment for model fitting
work.dir = getwd()
source.dir = fullarg[grep(pattern = "--file=",x = fullarg,fixed = TRUE)[1]]
source.dir = dirname(substr(source.dir,8,nchar(source.dir)))
setwd(source.dir)
source("fitPE_functions.R")
setwd(work.dir)
# interpret the arguments
# the job to continue analyses on
job.name = arg[1]
if(is.na(job.name)) stop("Job name is required and cannot be omitted!")
# read in phylogeny and trait from 1 RDS file
data.file = sprintf("%s.data.RDS",job.name)
tree = readRDS(data.file)
trait = tree$dat
tree = tree$phy
# read in fitting results
model.file = sprintf("%s.models.RDS",job.name)
res = readRDS(model.file)
scale.coef = res$scale
model.AIC = res$stat
model.params = res$params
# how many cores to use in the fitting
numCores=ceiling(as.numeric(arg[2]))
if(is.na(numCores)) numCores=1
# how far randomized starting points can go from the initial estimate
repRange = as.numeric(arg[3])
if(is.na(repRange)) repRange =10
# set manual fold-constraint between sizes
size.ratio = as.numeric(arg[4])
if(is.na(size.ratio)) size.ratio=10
#
ignore.zero = (as.numeric(arg[5])>0)
if(is.na(ignore.zero)) ignore.zero = FALSE
#
laplace = (as.numeric(arg[6])>0)
if(is.na(laplace)) laplace=TRUE
# when to stop the optimization in while loop
AIC.cutoff = as.numeric(arg[7])
if(is.na(AIC.cutoff)) AIC.cutoff=2
#
reltol = as.numeric(arg[8])
if(is.na(reltol)) reltol = 1e-4
# which is the best model
pebme2.params = model.params$pebme2
# fitting the models
# initialize key arguments
model.functions = c(pee3= "fitPE3.phy")
fixed.params = list(pee3= list(sigma=log(0)))
ini.functions = c(pee3= "initialize.parameters.TP")
# fit the model by fixing the first two Poisson process to get a good starting point 
ini.res = fitPE3.phy(phy = tree,x = sqrt(scale.coef)*trait[tree$tip.label],numCores = numCores,
                     size.ratio=size.ratio,ignore.zero=ignore.zero,laplace = laplace,
                      fixed = list(lambda1=log(unname(pebme2.params[1])),size1=log(unname(pebme2.params[2])),
                                   lambda2=log(unname(pebme2.params[3])),size2=log(unname(pebme2.params[2]/pebme2.params[4]-size.ratio)),
                                   sigma=log(0),epsilon=log(unname(pebme2.params[6]))),
                      start.value = list(lambda3=log(unname((1+size.ratio)*pebme2.params[5]/pebme2.params[4])),size3=log(1)))
#
start.params = do.call(ini.functions["pee3"],list(ini.res$params))
if(any(sapply(start.params,is.nan))) stop("Invalid starting parameters")
# fit the full PE3 model without restriction from PE2 models
tpe.res = fit.model.byNM.2step(fit.func = model.functions[["pee3"]],
                          start.params = list(start.value=start.params),
                          params1 =list(fixed=fixed.params[["pee3"]],size.ratio=size.ratio,ignore.zero=ignore.zero,laplace=laplace,
                                        phy=tree,x=sqrt(scale.coef)*trait[tree$tip.label],numCores=numCores,reltol=reltol),
                          params2 =list(fixed=fixed.params[["pee3"]],size.ratio=size.ratio,ignore.zero=ignore.zero,laplace=laplace,
                                        phy=tree,x=sqrt(scale.coef)*trait[tree$tip.label],numCores=numCores),
                          AIC.func = "get.AIC",
                          params.func = "get.params",
                          ini.func = ini.functions["pee3"],
                          AIC.cutoff = AIC.cutoff)
# save results
saveRDS(list(scale=scale.coef,res = list(list(tpe.res)),stat = c(pee3=tpe.res$AIC),params = list(pee3=tpe.res$params)),
        sprintf("%s.3PE.RDS",job.name))
#
warnings()
# output results
cat("Fitted model parameters\n")
unscale.params = list(pee3=summarize.model.parameter(params = tpe.res$params,time = 1,scale = scale.coef)$params)
print(unscale.params)
cat("Fitted model AIC\n")
zero.terminal.pic = findTerminalPIC(phy = tree,x = sqrt(scale.coef)*trait[tree$tip.label],remove.zero = FALSE)
unscale.AIC = normalize.AIC(AIC = c(pee3=tpe.res$AIC), scale = scale.coef,
                            sample.size = Nnode(tree)-sum(zero.terminal.pic$pic==0)*as.numeric(ignore.zero))
print(unscale.AIC)
```
