## Supplementary material for "Microbial genomic trait evolution is dominated by frequent and rare pulsed evolution": fit_5models.R

```
# read in key arguments for the fitting
arg=commandArgs(TRUE)
fullarg = commandArgs(FALSE)
# setup the environment for model fitting
work.dir = getwd()
source.dir = fullarg[grep(pattern = "--file=",x = fullarg,fixed = TRUE)[1]]
source.dir = dirname(substr(source.dir,8,nchar(source.dir)))
setwd(source.dir)
source("fitPE_functions.R")
setwd(work.dir)
# interpret the arguments
if(arg[1]==arg[2]){
  # read in phylogeny and trait from 1 RDS file
  tree = readRDS(arg[1])$phy
  trait = readRDS(arg[1])$dat
}else{
  # read in the phylogeny
  tree = read.tree(arg[1])
  # read in the trait value
  trait = read.table(file = arg[2],stringsAsFactors = FALSE)[,2]
  names(trait) = read.table(file = arg[2],stringsAsFactors = FALSE)[,1]
}
# set up a name for saving the result of this analysis
job.name = arg[3]
if(is.na(job.name)) stop("Job name is required and cannot be omitted!")
# how many cores to use in the fitting
numCores=ceiling(as.numeric(arg[4]))
if(is.na(numCores)) numCores=1
# how far randomized starting points can go from the initial estimate
repRange = as.numeric(arg[5])
if(is.na(repRange)) repRange =10
# set manual fold-constraint between sizes
size.ratio = as.numeric(arg[6])
if(is.na(size.ratio)) size.ratio=10
#
ignore.zero = (as.numeric(arg[7])>0)
if(is.na(ignore.zero)) ignore.zero = FALSE
#
laplace = (as.numeric(arg[8])>0)
if(is.na(laplace)) laplace=TRUE
#
AIC.cutoff = as.numeric(arg[9])
if(is.na(AIC.cutoff)) AIC.cutoff=2
# what's the tolerance for coarse-grind model fitting?
reltol = as.numeric(arg[10])
if(is.na(reltol)) reltol = 1e-4
# set manual scaling coefficient
scale.coef = as.numeric(arg[11])
# organize data for easy transfer to other analysis
this.data = list(phy=tree,dat=trait[tree$tip.label])
if(any(is.na(this.data$dat))) stop("NAs are not supported in tip values!")
saveRDS(this.data,sprintf("%s.data.RDS",job.name))
# estimating scaling coefficient, epsilon and sigma term so the optimizer have a reasonable starting point
raw.contrast = pic(x = trait[tree$tip.label],phy=tree,scaled = FALSE,var.contrasts = TRUE)
if(is.na(scale.coef)) scale.coef = 2/unname(var(raw.contrast[raw.contrast[,2]<=quantile(raw.contrast[,2],probs=0.05),1]))
sprintf("The scaling coefficient is %.3f",scale.coef)
est.sigma = var(pic(x = trait[tree$tip.label],phy=tree))*scale.coef
median.l = median(tree$edge.length)
# getting a reasonable starting point by fitting the pulsed evolution model with only tip pairs
# get all tip pairs
terminal.pic = findTerminalPIC(phy = tree,x = sqrt(scale.coef)*trait[tree$tip.label],remove.zero = TRUE)
# fit PE1(+BM) models with tip-only data
term.pee = fitPE(x = sapply(terminal.pic$x,function(x){x[1]-x[2]}),l = sapply(terminal.pic$l,sum),numCores = numCores,laplace = laplace,
                 start.value = list(lambda=log(1/median.l),size=log(est.sigma*median.l),epsilon=log(1)),
                 fixed = list(sigma=log(0)))
term.pebme = fitPE(x = sapply(terminal.pic$x,function(x){x[1]-x[2]}),l = sapply(terminal.pic$l,sum),numCores = numCores,laplace = laplace,
                   start.value = list(lambda=log(1/median.l),size=log(0.9*est.sigma*median.l),sigma=log(0.1*est.sigma),epsilon=log(1)))
# fit PE2(+BM) models with tip-only data
term.pee2 = fitPE2(x = sapply(terminal.pic$x,function(x){x[1]-x[2]}),l = sapply(terminal.pic$l,sum),numCores = numCores,size.ratio = size.ratio,laplace = laplace,
                   start.value = list(lambda1=log(0.1/median.l),size1=log(5*est.sigma*median.l),lambda2=log(1.1/median.l),size2=log(1),epsilon=log(1)),
                   fixed=list(sigma=log(0)))
term.pebme2 = fitPE2(x = sapply(terminal.pic$x,function(x){x[1]-x[2]}),l = sapply(terminal.pic$l,sum),numCores = numCores,size.ratio = size.ratio,laplace = laplace,
                     start.value = list(lambda1=log(0.1/median.l),size1=log(5*est.sigma*median.l),lambda2=log(1.1/median.l),size2=log(1),sigma=log(0.1*est.sigma),epsilon=log(1)))
# fitting the models with the full phylogeny
# initialize key arguments
model.functions = c(bme="fitPE.phy",pee = "fitPE.phy",pebme ="fitPE.phy",
                    pee2= "fitPE2.phy",pebme2="fitPE2.phy")
fixed.params = list(bme=list(lambda=log(0),size=log(0)),
                    pee = list(sigma=log(0)),pebme =list(),
                    pee2= list(sigma=log(0)),pebme2=list())
initial.parameters = list(bme=c(lambda=0,size=0,sigma=est.sigma,epsilon=1),
                          pee = term.pee$params,
                          pebme = term.pebme$params,
                          pee2 = term.pee2$params,
                          pebme2 = term.pebme2$params)
ini.multiplier = matrix(data = c(1,1,repRange,repRange,repRange,1/repRange,1/repRange,repRange,1/repRange,1/repRange),nrow = 5,ncol = 2,byrow = TRUE)
# preallocate result data
res = list(bme=list(),pee = list(),pebme =list(),pee2= list(),pebme2=list())
# fit every model
for(i in rev(1:5)){
  print(names(res)[i])
  for(rr in 1:5){
    print(rr)
    if(length(initial.parameters[[i]])>4){
      ini.func = "initialize.parameters.MP"
      start.params = initialize.parameters.MP(initial.parameters[[i]]*
                                 c(ini.multiplier[rr,1],ini.multiplier[rr,1],ini.multiplier[rr,2],ini.multiplier[rr,2],1,1))
    }else{
      ini.func = "initialize.parameters.SP"
      start.params = initialize.parameters.SP(initial.parameters[[i]]*
                                 c(ini.multiplier[rr,1],ini.multiplier[rr,2],1,1))
    }
    if(any(sapply(start.params,is.nan))) next
    try({
      mdl= fit.model.byNM.2step(fit.func = model.functions[[i]],
                         start.params = list(start.value=start.params),
                         params1 =list(fixed=fixed.params[[i]],size.ratio=size.ratio,ignore.zero=ignore.zero,
                                       phy=tree,x=sqrt(scale.coef)*trait[tree$tip.label],numCores=numCores,laplace = laplace,reltol=reltol),
                         params2 =list(fixed=fixed.params[[i]],size.ratio=size.ratio,ignore.zero=ignore.zero,
                                       phy=tree,x=sqrt(scale.coef)*trait[tree$tip.label],numCores=numCores,laplace = laplace),
                         AIC.func = "get.AIC",
                         params.func = "get.params",
                         ini.func = ini.func,
                         AIC.cutoff = AIC.cutoff)
      res[[i]][[rr]] = mdl
      })
    #
    saveRDS(list(scale=scale.coef,res = res),
            sprintf("%s.models.RDS",job.name))
  }
}
# summarize results
model.summary = lapply(res,function(x){
  best.rep = which.min(sapply(x,function(y){if(is.null(y)){
    return(NA)
  }else{
    return(y$low$AIC)
  }}))
  best.AIC = x[[best.rep]]$low$AIC
  best.params = x[[best.rep]]$low$params
  return(list(AIC=best.AIC,params=best.params))
})
# save results
saveRDS(list(scale=scale.coef,res = res,stat=sapply(model.summary,get.AIC),params=lapply(model.summary,get.params)),
        sprintf("%s.models.RDS",job.name))
#
warnings()
# output results
cat("Fitted model parameters\n")
unscale.params = lapply(model.summary,function(x){summarize.model.parameter(params = get.params(x),time = 1,scale = scale.coef)$params})
print(unscale.params)
cat("Fitted model AIC\n")
zero.terminal.pic = findTerminalPIC(phy = tree,x = sqrt(scale.coef)*trait[tree$tip.label],remove.zero = FALSE)
unscale.AIC = normalize.AIC(AIC = sapply(model.summary,get.AIC), scale = scale.coef,
                            sample.size = Nnode(tree)-sum(zero.terminal.pic$pic==0)*as.numeric(ignore.zero))
print(unscale.AIC)
```
