## Supplementary material for "Microbial genomic trait evolution is dominated by frequent and rare pulsed evolution": fitPE_functions.R

```
# load required packages
library(ape)
library(bbmle)
library(pracma)
library(doParallel)
library(extraDistr)
# load extended functions to ape package
source("ape_addition.R")
# load probability functions
source("PEpois.R")

### function to select only independent tip-pair contrasts
## phy is a phylo-class from ape package
## x is a named vector that contains the trait value for each tip
## remove.zero determines if zero-contrasts are removed from the results
# output value of this function is a list whose elements are vectors/lists
# of PIC, trait value, branch length,node index and tip labels
findTerminalPIC = function (phy, x, remove.zero = FALSE) {
  # add node labels if not already exist
  if (is.null(phy$node.label)) 
    phy = makeNodeLabel(phy)
  # keep only tip-values
  x = x[phy$tip.label]
  # find nodes that have only tip-children
  tip.node = setdiff(sapply(1:Ntip(phy), getLatestAncestor, 
                            phy = phy), sapply(Ntip(phy) + 1:phy$Nnode, getLatestAncestor, 
                                               phy = phy))
  # tip labels for the tip-pairs
  tip.label = lapply(tip.node, function(n) {
    phy$tip.label[phy$edge[phy$edge[, 1] == n, 2]]
  })
  # branches leading to the tips
  term.l = lapply(tip.node, function(n) {
    phy$edge.length[phy$edge[, 1] == n]
  })
  # tip values of the tip-pairs
  term.x = lapply(tip.node, function(n) {
    x[phy$edge[phy$edge[, 1] == n, 2]]
  })
  # get all phylogenetically independent contrasts
  phy.pic = pic(x = x, phy = phy)
  # get PIC of tip-pairs
  term.pic = phy.pic[tip.node - Ntip(phy)]
  # remove zero-contrasts if required
  if (remove.zero) {
    p0 = sum(term.pic == 0)/length(term.pic)
    idx = term.pic != 0
    return(list(pic = term.pic[idx], x = term.x[idx], l = term.l[idx], 
                node = tip.node[idx],tip = tip.label[idx] ,p0 = p0))
  }
  else {
    return(list(pic = term.pic, x = term.x, l = term.l, node = tip.node, tip = tip.label))
  }
}

### function to quickly reconstruct ancestral states and calculate PIC with time-independent variation
## phy is a phylo-class from ape package
## x is a named vector that contains the trait value for each tip
## rate is the total rate of evolution (variance of trait change per unit branch length)
## epsilon is the variance of time-independent variation between nodes
# output value of this function is a list whose elements are vectors of tip and ancestral states, tip and
# ancestral uncertainty, and trait change, branch length, total uncertainty, total variance between nodes
reconstructAncestralStates = function(phy,x,rate,epsilon){
  # pre-allocate ancestral states and their corresponding variance
  trait = c(x[phy$tip.label],numeric(phy$Nnode))
  trait.var = c(rep(epsilon/2,Ntip(phy)),numeric(phy$Nnode))
  # pre-allocate trait difference, branch length and time-independent variation between nodes
  dx = numeric(phy$Nnode)
  dl = dx
  de = dx
  # find the order of branches to reconstruct from tip to root
  idx = reorder.phylo(x = phy,order = "postorder",index.only = TRUE)
  # reconstruct ancestral state using Felsenstein's method
  for(i in seq(1,length(idx),2)){
    # find the index of ancestor and descendants 
    anc.idx = phy$edge[idx[i],1]
    des.idx = phy$edge[idx[c(i,i+1)],2]
    # find trait values and branch lengths
    this.x = trait[des.idx]
    this.l = phy$edge.length[idx[c(i,i+1)]]
    # find time-independent variation
    this.epsilon = trait.var[des.idx]
    # reconstruct ancestral states by Felsenstein's PIC method
    trait[anc.idx] = (this.x[1]*(this.l[2]*rate+this.epsilon[2])+
                        this.x[2]*(this.l[1]*rate+this.epsilon[1]))/
      (sum(this.l)*rate+sum(this.epsilon))
    # the uncertainty (time-independent variation) of reconstructed ancestral states
    trait.var[anc.idx] = ((this.l[2]*rate+this.epsilon[2])*
                            (this.l[1]*rate+this.epsilon[1]))/
      (sum(this.l)*rate+sum(this.epsilon))
    # get trait change, total branch length and total time-independent variation betwee nodes
    dx[anc.idx-Ntip(phy)] = this.x[1]-this.x[2]
    dl[anc.idx-Ntip(phy)] = sum(this.l)
    de[anc.idx-Ntip(phy)] = sum(this.epsilon)
  }
  return(list(trait=trait,var=trait.var,contrast=dx,l = dl,epsilon=de,lprime=dl*rate+de))
}

### function to calculate pseudo-PIC that normalizes trait changes into standard normal distribution
## phy is a phylo-class from ape package
## x is a named vector that contains the trait value for each tip
## pfunc is one of the cumulative probability function in PEpois.R
## params is the parameter of the pfunc
# output value of this function is a vector of pseudo-PIC
pseudo.pic = function(phy,x,pfunc,params){
  # get the number of Poisson processes
  numPE = length(params)/2-1
  # calculate the total evolution rate (variance of trait change per unit branch length)
  rate = sum(sapply(1:numPE,function(i){
    prod(params[-1:0 + 2*i])
  }))+params[length(params)-1]
  # reconstruct ancestral states
  epsilon = params[length(params)]
  anc = reconstructAncestralStates(phy = phy,x=x,rate= rate,epsilon=epsilon)
  # calculate the cumulative probability of trait differences between nodes under the distribution defined by pfunc
  pp = sapply(1:length(anc$contrast),function(i){
    do.call(pfunc,c(list(q=anc$contrast[i],t=anc$l[i],epsilon=anc$epsilon[i]),as.list(params[-length(params)])))
    })
  # transform trait differences into normal quantiles (pseudo-PIC)
  p.pic = qnorm(p = pp)
  return(p.pic)
}

### function to get parameters from the fitting output
## x is a list with an element named params, which is the output of fitting functions in this script
get.params = function(x){x$params}

### function to get AIC from the fitting output
## x is a list with an element named AIC, which is the output of fitting functions in this script
get.AIC = function(x){x$AIC}

### function to calculate the process variance
## params is a named vector of parameters whose elements is oredered as:
## [lambda, size], sigma, epsilon
## the elements in the bracket can repeat several times
total.process.variance = function(params){
  # number of Poisson processes
  numPE = length(params)/2-1
  # process variance of each Poisson process
  process.variance = sapply(1:numPE,function(i){
    params[2*i-1]*params[2*i]
  })
  # the total process variance
  return(sum(process.variance)+params[2*numPE+1])
}

### function to log-transform the model parameters before fitting
## x is a named vector of parameters
## size.ratio is the constraint of jump size between each Poisson process
initialize.parameters = function(x,size.ratio){
  numPE = length(x)/2-1
  for(i in seq(numPE,2,-1)){
    x[paste0("size",i)]=x[paste0("size",i-1)]/x[paste0("size",i)]-size.ratio
  }
  return(as.list(log(x)))
}

### function to log-transform the model parameters before fitting (PE1)
## x is a named vector of parameters
initialize.parameters.SP = function(x,...){as.list(log(x))}

### function to log-transform the model parameters before fitting (PE2)
## x is a named vector of parameters
initialize.parameters.MP = function(x,...){
  x["size2"]=x["size1"]/x["size2"]-size.ratio
  return(as.list(log(x)))
}

### function to log-transform the model parameters before fitting (PE3)
## x is a named vector of parameters
initialize.parameters.TP = function(x,...){
  x["size3"]=x["size2"]/x["size3"]-size.ratio
  x["size2"]=x["size1"]/x["size2"]-size.ratio
  return(as.list(log(x)))
}

### function to fit a model through high-low tolerance phases
## fit.func is one of the model fitting functions defined in this script
## start.params is a list whose first element contains the list of starting model parameters
## params1 and params2 are the other arguments passed to the fitting function in the two phases
## AIC.func is the function to extract AIC from the results
## params.function is the function to extract parameters from the results
## ini.function is the function to transform the model parameters before fitting
## AIC.cutoff is the threshold to stop fitting
# output value of this function is a list that contains the final AIC, final parameter and results of each phase
fit.model.byNM.2step = function(fit.func="mle2",start.params,params1,params2,
                                AIC.func="AIC",params.func=function(x){x@coef},ini.func="as.list",
                                AIC.cutoff=2,...){
  # fit high-tolerance phase model
  res1 = fit.model.byNM(fit.func=fit.func,start.params = start.params,params=params1,
                        AIC.func=AIC.func,params.func=params.func,ini.func=ini.func,
                        AIC.cutoff=AIC.cutoff,...)
  # if infinite AIC is returned, stop fitting
  if(is.infinite(res1$AIC)) return(list(params = res1$params, AIC = res1$AIC,high=res1,low=NULL))
  # update starting parameters
  start.params[[1]] = do.call(ini.func,list(res1$params))
  # fit low-tolerance phase model
  res2 = fit.model.byNM(fit.func=fit.func,start.params=start.params,params=params2,
                        AIC.func=AIC.func,params.func=params.func,ini.func=ini.func,
                        AIC.cutoff=AIC.cutoff,...)
  return(list(params = res2$params, AIC = res2$AIC,high=res1,low=res2))
}

### function to fit a model by maximum likelihood with multiple steps
## fit.func is one of the model fitting functions defined in this script
## start.params is a list whose first element contains the list of starting model parameters
## params is the other arguments passed to the fitting function
## AIC.func is the function to extract AIC from the results
## params.function is the function to extract parameters from the results
## ini.function is the function to transform the model parameters before fitting
## AIC.cutoff is the threshold to stop fitting
# output value of this function is a list that contains the final AIC, final parameter and results of each step
fit.model.byNM = function(fit.func="mle2",start.params,params,
                          AIC.func="AIC",params.func=function(x){x@coef},ini.func="as.list",
                          AIC.cutoff=2,...){
  # initialize AICs
  lastAIC = Inf
  deltaAIC = Inf
  # initialize index
  i = 0
  # get initial parameters
  ini.params = start.params[[1]]
  # evaluate starting AIC
  res = list(do.call(fit.func,c(params,start.params,list(eval.only=TRUE))))
  # if starting AIC is infinite, stop fitting
  if(is.infinite(res[[1]]$AIC)) return(list(AIC = res[[1]]$AIC,params=res[[1]]$params,full=res))
  # continues to fit until AIC cutoff is reached
  while(deltaAIC>AIC.cutoff){
    t0= Sys.time()
    # update starting parameters
    start.params[[1]] = ini.params
    # fit the model again
    mdl = do.call(fit.func,c(params,start.params))
    # update the model parameters
    current.params = do.call(params.func,list(mdl))
    # transform the new model parameters
    ini.params = do.call(ini.func,list(current.params))
    # update AIC
    currentAIC = do.call(AIC.func,list(mdl))
    deltaAIC = lastAIC - currentAIC
    lastAIC = currentAIC
    # update index
    i = i+1
    t1=Sys.time()
    res[[i+1]] = list(AIC=currentAIC,params=current.params,time=list(t0=t0,t1=t1))
  }
  return(list(AIC = currentAIC,params=current.params,full=res))
}


### functions to fit BM and PE1 models without a phylogeny
## x is a vector of trait changes between nodes
## l is a vector of branch lengths between nodes
## start.value is a list of starting parameters
## numCores is the number of cores when run in parallel
## fixed is a list of fixed parameters
## laplace determines if Laplace distribution is used for time-independent variation
## eval.only determines if the function should only evaluate the AIC of starting parameters
## reltol is the relative tolerance of the optimizer
# output of these functions is a list of AIC, model parameter and detailed optimization results.
fitPE = function(x,l,start.value=NULL,numCores=1,fixed=list(),laplace=TRUE,eval.only=FALSE,reltol=sqrt(.Machine$double.eps),...){
  # set the probability function to use
  if(laplace){
    dfunc = dPEpoislaplace
  }else{
    dfunc = dPEpoisnorm
  }
  # set the likelihood function
  if(numCores>1){
    # Setup parallel backend to use many processors
    cores=detectCores()
    cat(sprintf("There's %d cores available\n",cores))
    #cl <- makeCluster(min(cores[1]-1,numCores),type="PSOCK") # Not to overload
    cl <- makeCluster(min(cores[1]-1,numCores),type = "FORK") # Not to overload
    cat(sprintf("Using %d cores\n",min(cores[1]-1,numCores)))
    registerDoParallel(cl)
    cat(sprintf("%d cluster registered\n",min(cores[1]-1,numCores)))
    LL = function(lambda,size,sigma,epsilon){
      lambda= exp(lambda)
      size = exp(size)
      sigma=exp(sigma)
      epsilon=exp(epsilon)
      pp = parSapply(cl,1:length(x),function(i){dfunc(x = x[i],t = l[i],lambda = lambda,size=size,sigma = sigma,epsilon=epsilon)})
      ll = -sum(log(pp))
      return(ll)
    }
  }else{
    LL = function(lambda,size,sigma,epsilon){
      lambda= exp(lambda)
      size = exp(size)
      sigma=exp(sigma)
      epsilon=exp(epsilon)
      pp = sapply(1:length(x),function(i){dfunc(x = x[i],t = l[i],lambda = lambda,size=size,sigma = sigma,epsilon=epsilon)})
      ll = -sum(log(pp))
      return(ll)
    }
  }
  # if there is no starting parameters, set one
  if(is.null(start.value)) start.value = list(lambda=log(1),size=log(1),sigma = log(var(x/sqrt(l))),epsilon=log(var(x)))
  # remove fixed parameters from the starting parameters
  start.value[names(fixed)]=NULL
  # set the method to do optimization
  if(length(start.value)<2){
    method  = "BFGS"
  }else{
    method = "Nelder-Mead"
  }
  # evaluate AIC only if instructed
  if(eval.only){
    res = exp(unlist(c(start.value,fixed)))
    names(res) = c(names(start.value),names(fixed))
    res = res[c("lambda","size","sigma","epsilon")]
    this.AIC = 2*do.call(LL,c(start.value,fixed))+2*length(start.value)
    if(numCores>1) stopCluster(cl)
    return(list(params=res,
                AIC=this.AIC,
                raw=NA))
  }
  # optimize for the best model parameters
  result = do.call("mle2", 
                   c(list(LL, 
                          start = start.value, 
                          data = list(x=x),
                          fixed = fixed,
                          method = method,
                          control = list(trace=TRUE,maxit=2000,reltol=reltol)
                   )))
  # transform the parameters back to the linear scale 
  res = exp(result@fullcoef)
  # stop parallel clusters
  if(numCores>1) stopCluster(cl)
  return(list(params=res,AIC=AIC(result),raw= result))
}

### functions to fit PE2 models without a phylogeny
## x is a vector of trait changes between nodes
## l is a vector of branch lengths between nodes
## start.value is a list of starting parameters
## numCores is the number of cores when run in parallel
## size.ratio restricts the ratio between sizes of different Poisson processes
## fixed is a list of fixed parameters
## laplace determines if Laplace distribution is used for time-independent variation
## eval.only determines if the function should only evaluate the AIC of starting parameters
## reltol is the relative tolerance of the optimizer
# output of these functions is a list of AIC, model parameter and detailed optimization results.
fitPE2 = function(x,l,start.value=NULL,numCores=1,size.ratio=10,fixed=list(),laplace=TRUE,eval.only=FALSE,reltol=sqrt(.Machine$double.eps),...){
  # set the probability function to use
  if(laplace){
    dfunc = dPEpois2laplace
  }else{
    dfunc = dPEpois2norm
  }
  # set the likelihood function
  if(numCores>1){
    # Setup parallel backend to use many processors
    cores=detectCores()
    cat(sprintf("There's %d cores available\n",cores))
    #cl <- makeCluster(min(cores[1]-1,numCores),type="PSOCK") # Not to overload
    cl <- makeCluster(min(cores[1]-1,numCores),type = "FORK") # Not to overload
    cat(sprintf("Using %d cores\n",min(cores[1]-1,numCores)))
    registerDoParallel(cl)
    cat(sprintf("%d cluster registered\n",min(cores[1]-1,numCores)))
    LL = function(lambda1,size1,lambda2,size2,sigma,epsilon){
      lambda1= exp(lambda1)
      size1 = exp(size1)
      lambda2= exp(lambda2)
      size2 = size1/(size.ratio+exp(size2))
      sigma=exp(sigma)
      epsilon=exp(epsilon)
      pp = parSapply(cl,1:length(x),function(i){
        dfunc(x = x[i],t = l[i],lambda1 = lambda1,size1=size1,lambda2 = lambda2,size2 = size2,sigma = sigma,epsilon=epsilon)
      })
      ll = -sum(log(pp))
      return(ll)
    }
  }else{
    LL = function(lambda1,size1,lambda2,size2,sigma,epsilon){
      lambda1= exp(lambda1)
      size1 = exp(size1)
      lambda2= exp(lambda2)
      size2 = size1/(size.ratio+exp(size2))
      sigma=exp(sigma)
      epsilon=exp(epsilon)
      pp = sapply(1:length(x),function(i){
        dfunc(x = x[i],t = l[i],lambda1 = lambda1,size1=size1,lambda2 = lambda2,size2 = size2,sigma = sigma,epsilon=epsilon)
      })
      ll = -sum(log(pp))
      return(ll)
    }
  }
  # if there is no starting parameters, set one
  if(is.null(start.value)) start.value = list(lambda1=log(1),size1=log(1),lambda2=log(1),size2=log(1),sigma = log(var(x/sqrt(l))),epsilon=log(sd(x)))
  # remove fixed parameters from the starting parameters
  start.value[names(fixed)]=NULL
  # set the method to do optimization
  if(length(start.value)<2){
    method  = "BFGS"
  }else{
    method = "Nelder-Mead"
  }
  # evaluate AIC only if instructed
  if(eval.only){
    res = exp(unlist(c(start.value,fixed)))
    names(res) = c(names(start.value),names(fixed))
    res = res[c("lambda1","size1","lambda2","size2","sigma","epsilon")]
    res[4] = res[2]/(size.ratio+res[4])
    this.AIC = 2*do.call(LL,c(start.value,fixed))+2*length(start.value)
    if(numCores>1) stopCluster(cl)
    return(list(params=res,
                AIC=this.AIC,
                raw=NA))
  }
  # optimize for the best model parameters
  result = do.call("mle2", 
                   c(list(LL, 
                          start = start.value, 
                          data = list(x=x),
                          fixed = fixed,
                          method = method,
                          control = list(trace=TRUE,maxit=2000,reltol=reltol)
                   )))
  # transform the parameters back to the linear scale 
  res = exp(result@fullcoef)
  res[4] = res[2]/(size.ratio+res[4])
  # stop parallel clusters
  if(numCores>1) stopCluster(cl)
  return(list(params=res,AIC=AIC(result),raw= result))
}

### functions to fit PE3 models without a phylogeny
## x is a vector of trait changes between nodes
## l is a vector of branch lengths between nodes
## start.value is a list of starting parameters
## numCores is the number of cores when run in parallel
## size.ratio restricts the ratio between sizes of different Poisson processes
## fixed is a list of fixed parameters
## laplace determines if Laplace distribution is used for time-independent variation
## eval.only determines if the function should only evaluate the AIC of starting parameters
## reltol is the relative tolerance of the optimizer
# output of these functions is a list of AIC, model parameter and detailed optimization results.
fitPE3 = function(x,l,start.value=NULL,numCores=1,size.ratio=10,fixed=list(),laplace=TRUE,eval.only=FALSE,reltol=sqrt(.Machine$double.eps),...){
  # set the probability function to use
  if(laplace){
    dfunc = dPEpois3laplace
  }else{
    dfunc = dPEpois3norm
  }
  # set the likelihood function
  if(numCores>1){
    # Setup parallel backend to use many processors
    cores=detectCores()
    cat(sprintf("There's %d cores available\n",cores))
    #cl <- makeCluster(min(cores[1]-1,numCores),type="PSOCK") # Not to overload
    cl <- makeCluster(min(cores[1]-1,numCores),type = "FORK") # Not to overload
    cat(sprintf("Using %d cores\n",min(cores[1]-1,numCores)))
    registerDoParallel(cl)
    cat(sprintf("%d cluster registered\n",min(cores[1]-1,numCores)))
    LL = function(lambda1,size1,lambda2,size2,lambda3,size3,sigma,epsilon){
      lambda1= exp(lambda1)
      size1 = exp(size1)
      lambda2= exp(lambda2)
      size2 = size1/(size.ratio+exp(size2))
      lambda3= exp(lambda3)
      size3 = size2/(size.ratio+exp(size3))
      sigma=exp(sigma)
      epsilon=exp(epsilon)
      pp = parSapply(cl,1:length(x),function(i){
        dfunc(x = x[i],t = l[i],lambda1 = lambda1,size1=size1,lambda2 = lambda2,size2 = size2,
              lambda3 = lambda3,size3=size3,sigma = sigma,epsilon=epsilon)
      })
      ll = -sum(log(pp))
      return(ll)
    }
  }else{
    LL = function(lambda1,size1,lambda2,size2,lambda3,size3,sigma,epsilon){
      lambda1= exp(lambda1)
      size1 = exp(size1)
      lambda2= exp(lambda2)
      size2 = size1/(size.ratio+exp(size2))
      lambda3= exp(lambda3)
      size3 = size2/(size.ratio+exp(size3))
      sigma=exp(sigma)
      epsilon=exp(epsilon)
      pp = sapply(1:length(x),function(i){
        dfunc(x = x[i],t = l[i],lambda1 = lambda1,size1=size1,lambda2 = lambda2,size2 = size2,
              lambda3 = lambda3,size3=size3,sigma = sigma,epsilon=epsilon)
      })
      ll = -sum(log(pp))
      return(ll)
    }
  }
  # check that starting parameters are supplied
  if(is.null(start.value)) stop("Initial values must be specified!")
  # remove fixed parameters from the starting parameters
  start.value[names(fixed)]=NULL
  # set the method to do optimization
  if(length(start.value)<2){
    method  = "BFGS"
  }else{
    method = "Nelder-Mead"
  }
  # evaluate AIC only if instructed
  if(eval.only){
    res = exp(unlist(c(start.value,fixed)))
    names(res) = c(names(start.value),names(fixed))
    res = res[c("lambda1","size1","lambda2","size2","lambda3","size3","sigma","epsilon")]
    res[4] = res[2]/(size.ratio+res[4])
    res[6] = res[4]/(size.ratio+res[6])
    this.AIC = 2*do.call(LL,c(start.value,fixed))+2*length(start.value)
    if(numCores>1) stopCluster(cl)
    return(list(params=res,
                AIC=this.AIC,
                raw=NA))
  }
  # optimize for the best model parameters
  result = do.call("mle2", 
                   c(list(LL, 
                          start = start.value, 
                          data = list(x=x),
                          fixed = fixed,
                          method = method,
                          control = list(trace=TRUE,maxit=2000,reltol=reltol)
                   )))
  # transform the parameters back to the linear scale 
  res = exp(result@fullcoef)
  res[4] = res[2]/(size.ratio+res[4])
  res[6] = res[4]/(size.ratio+res[6])
  # stop parallel clusters
  if(numCores>1) stopCluster(cl)
  return(list(params=res,AIC=AIC(result),raw= result))
}

### functions to fit PE3 models along a phylogeny
## phy is a phylo-class object from ape package
## x is a vector of trait values of the tips
## start.value is a list of starting parameters
## numCores is the number of cores when run in parallel
## size.ratio restricts the ratio between sizes of different Poisson processes
## fixed is a list of fixed parameters
## ignore.zero determines if zero-contrasts are removed from the optimization
## laplace determines if Laplace distribution is used for time-independent variation
## eval.only determines if the function should only evaluate the AIC of starting parameters
## bootstrap is a vector of node indexes that should be used in the optimization 
## reltol is the relative tolerance of the optimizer
# output of these functions is a list of AIC, model parameter and detailed optimization results.
fitPE3.phy = function(phy,x,start.value=NULL,numCores=1,size.ratio=10,fixed=list(),ignore.zero=TRUE,
                      laplace=TRUE,eval.only=FALSE,bootstrap=NULL,reltol=sqrt(.Machine$double.eps),...){
  # initialize node indexes if not supplied
  if(is.null(bootstrap)) bootstrap = 1:phy$Nnode
  # remove zero-contrasts if instructed
  if(ignore.zero) bootstrap = intersect(bootstrap,
                                        which(reconstructAncestralStates(phy = phy,x=x[phy$tip.label],rate = 1,epsilon = 0)$contrast!=0))
  # set probability function to use
  if(laplace){
    dfunc = dPEpois3laplace
  }else{
    dfunc = dPEpois3norm
  }
  # set likelihood function
  if(numCores>1){
    cores=detectCores()
    cat(sprintf("There's %d cores available\n",cores))
    cl <- makeCluster(min(cores[1]-1,numCores),type = "FORK") # Not to overload
    cat(sprintf("Using %d cores\n",min(cores[1]-1,numCores)))
    registerDoParallel(cl)
    cat(sprintf("%d cluster registered\n",min(cores[1]-1,numCores)))
    LL = function(lambda1,size1,lambda2,size2,lambda3,size3,sigma,epsilon){
      lambda1= exp(lambda1)
      size1 = exp(size1)
      lambda2= exp(lambda2)
      size2 = size1/(size.ratio+exp(size2))
      lambda3= exp(lambda3)
      size3 = size2/(size.ratio+exp(size3))
      sigma=exp(sigma)
      epsilon=exp(epsilon)
      rate = lambda1*size1+lambda2*size2+lambda3*size3+sigma
      anc = reconstructAncestralStates(phy = phy,x=x[phy$tip.label],rate = rate,epsilon = epsilon)
      pp = parSapply(cl,bootstrap,function(i){
        dfunc(x = anc$contrast[i],t = anc$l[i],lambda1 = lambda1,size1=size1,lambda2 = lambda2,size2 = size2,
              lambda3 = lambda3,size3 = size3,sigma = sigma,epsilon=anc$epsilon[i])
      })
      ll = -sum(log(pp))
      return(ll)
    }
  }else{
    LL = function(lambda1,size1,lambda2,size2,lambda3,size3,sigma,epsilon){
      lambda1= exp(lambda1)
      size1 = exp(size1)
      lambda2= exp(lambda2)
      size2 = size1/(size.ratio+exp(size2))
      lambda3= exp(lambda3)
      size3 = size2/(size.ratio+exp(size3))
      sigma=exp(sigma)
      epsilon=exp(epsilon)
      rate = lambda1*size1+lambda2*size2+lambda3*size3+sigma
      anc = reconstructAncestralStates(phy = phy,x=x[phy$tip.label],rate = rate,epsilon = epsilon)
      pp = sapply(bootstrap,function(i){
        dfunc(x = anc$contrast[i],t = anc$l[i],lambda1 = lambda1,size1=size1,lambda2 = lambda2,size2 = size2,
              lambda3 = lambda3,size3 = size3,sigma = sigma,epsilon=anc$epsilon[i])
      })
      ll = -sum(log(pp))
      return(ll)
    }
  }
  # if no starting parameters are supplied, set one 
  if(is.null(start.value)) initialize.parameters.TP(c(lambda1=0.1,size1=121,lambda2=1,size2=11,
                                                      lambda3=10,size3=1,sigma=1,epsilon=1))
  # remove fixed parameters from the starting parameters
  start.value[names(fixed)]=NULL
  # set optimization method
  if(length(start.value)<2){
    method  = "BFGS"
  }else{
    method = "Nelder-Mead"
  }
  # evaluate AIC only if instructed
  if(eval.only){
    res = exp(unlist(c(start.value,fixed)))
    names(res) = c(names(start.value),names(fixed))
    res = res[c("lambda1","size1","lambda2","size2","lambda3","size3","sigma","epsilon")]
    res[4] = res[2]/(size.ratio+res[4])
    res[6] = res[4]/(size.ratio+res[6])
    this.AIC = 2*do.call(LL,c(start.value,fixed))+2*length(start.value)
    if(numCores>1) stopCluster(cl)
    return(list(params=res,
                AIC=this.AIC,
                raw=NA))
  }
  # search for the best model parameters
  result = do.call("mle2", 
                   c(list(LL, 
                          start = start.value, 
                          data = list(x=x),
                          fixed = fixed,
                          method = method,
                          control = list(trace=TRUE,maxit=2000,reltol=reltol)
                   )))
  # transform model parameters back to the linear scale
  res = exp(result@fullcoef)
  res[4] = res[2]/(size.ratio+res[4])
  res[6] = res[4]/(size.ratio+res[6])
  # stop parallel clusters
  if(numCores>1) stopCluster(cl)
  return(list(params=res,AIC=AIC(result),raw= result))
}

### functions to fit BM and PE1 models along a phylogeny
## phy is a phylo-class object from ape package
## x is a vector of trait values of the tips
## start.value is a list of starting parameters
## numCores is the number of cores when run in parallel
## fixed is a list of fixed parameters
## ignore.zero determines if zero-contrasts are removed from the optimization
## laplace determines if Laplace distribution is used for time-independent variation
## eval.only determines if the function should only evaluate the AIC of starting parameters
## bootstrap is a vector of node indexes that should be used in the optimization 
## reltol is the relative tolerance of the optimizer
# output of these functions is a list of AIC, model parameter and detailed optimization results.
fitPE.phy = function(phy,x,start.value=NULL,numCores=1,fixed=list(),ignore.zero=TRUE,
                     laplace=TRUE,eval.only=FALSE,bootstrap=NULL,reltol=sqrt(.Machine$double.eps),...){
  # initialize node indexes if not supplied
  if(is.null(bootstrap)) bootstrap = 1:phy$Nnode
  # remove zero-contrasts if instructed
  if(ignore.zero) bootstrap = intersect(bootstrap,
                                        which(reconstructAncestralStates(phy = phy,x=x[phy$tip.label],rate = 1,epsilon = 0)$contrast!=0))
  # set probability function to use
  if(laplace){
    dfunc = dPEpoislaplace
  }else{
    dfunc = dPEpoisnorm
  }
  # set likelihood function
  if(numCores>1){
    # Setup parallel backend to use many processors
    cores=detectCores()
    cat(sprintf("There's %d cores available\n",cores))
    #cl <- makeCluster(min(cores[1]-1,numCores),type="PSOCK") # Not to overload
    cl <- makeCluster(min(cores[1]-1,numCores),type = "FORK") # Not to overload
    cat(sprintf("Using %d cores\n",min(cores[1]-1,numCores)))
    registerDoParallel(cl)
    cat(sprintf("%d cluster registered\n",min(cores[1]-1,numCores)))
    LL = function(lambda,size,sigma,epsilon){
      lambda= exp(lambda)
      size = exp(size)
      sigma=exp(sigma)
      epsilon=exp(epsilon)
      rate = lambda*size+sigma
      anc = reconstructAncestralStates(phy = phy,x=x[phy$tip.label],rate = rate,epsilon = epsilon)
      pp = parSapply(cl,bootstrap,function(i){
        dfunc(x = anc$contrast[i],t = anc$l[i],lambda = lambda,size=size,sigma = sigma,epsilon=anc$epsilon[i])
      })
      ll = -sum(log(pp))
      return(ll)
    }
  }else{
    LL = function(lambda,size,sigma,epsilon){
      lambda= exp(lambda)
      size = exp(size)
      sigma=exp(sigma)
      epsilon=exp(epsilon)
      rate = lambda*size+sigma
      anc = reconstructAncestralStates(phy = phy,x=x[phy$tip.label],rate = rate,epsilon = epsilon)
      pp = sapply(bootstrap,function(i){
        dfunc(x = anc$contrast[i],t = anc$l[i],lambda = lambda,size=size,sigma = sigma,epsilon=anc$epsilon[i])
      })
      ll = -sum(log(pp))
      return(ll)
    }
  }
  # if no starting parameters are supplied, set one
  if(is.null(start.value)) start.value = list(lambda=log(1),size=log(1),sigma = log(var(pic(x = x[phy$tip.label],phy = phy))),epsilon=log(1))
  # remove fixed parameters from the starting parameters
  start.value[names(fixed)]=NULL
  # set optimization method
  if(length(start.value)<2){
    method  = "BFGS"
  }else{
    method = "Nelder-Mead"
  }
  # evaluate AIC only if instructed
  if(eval.only){
    res = exp(unlist(c(start.value,fixed)))
    names(res) = c(names(start.value),names(fixed))
    res = res[c("lambda","size","sigma","epsilon")]
    this.AIC = 2*do.call(LL,c(start.value,fixed))+2*length(start.value)
    if(numCores>1) stopCluster(cl)
    return(list(params=res,
                AIC=this.AIC,
                raw=NA))
  }
  # search for the best model parameters
  result = do.call("mle2", 
                   c(list(LL, 
                          start = start.value, 
                          data = list(x=x),
                          fixed = fixed,
                          method = method,
                          control = list(trace=TRUE,maxit=2000,reltol=reltol)
                   )))
  # transform model parameters back to the linear scale
  res = exp(result@fullcoef)
  # stop parallel clusters
  if(numCores>1) stopCluster(cl)
  return(list(params=res,AIC=AIC(result),raw= result))
}

### functions to fit PE2 models without a phylogeny
## phy is a phylo-class object from ape package
## x is a vector of trait values of the tips
## start.value is a list of starting parameters
## numCores is the number of cores when run in parallel
## size.ratio restricts the ratio between sizes of different Poisson processes
## fixed is a list of fixed parameters
## ignore.zero determines if zero-contrasts are removed from the optimization
## laplace determines if Laplace distribution is used for time-independent variation
## eval.only determines if the function should only evaluate the AIC of starting parameters
## bootstrap is a vector of node indexes that should be used in the optimization 
## reltol is the relative tolerance of the optimizer
# output of these functions is a list of AIC, model parameter and detailed optimization results.
fitPE2.phy = function(phy,x,start.value=NULL,numCores=1,size.ratio=10,fixed=list(),ignore.zero=TRUE,
                      laplace=TRUE,eval.only=FALSE,bootstrap=NULL,reltol=sqrt(.Machine$double.eps),...){
  # initialize node indexes if not supplied
  if(is.null(bootstrap)) bootstrap = 1:phy$Nnode
  # remove zero-contrasts if instructed
  if(ignore.zero) bootstrap = intersect(bootstrap,
                                        which(reconstructAncestralStates(phy = phy,x=x[phy$tip.label],rate = 1,epsilon = 0)$contrast!=0))
  # set probability function to use
  if(laplace){
    dfunc = dPEpois2laplace
  }else{
    dfunc = dPEpois2norm
  }
  # set likelihood function
  if(numCores>1){
    # Setup parallel backend to use many processors
    cores=detectCores()
    cat(sprintf("There's %d cores available\n",cores))
    #cl <- makeCluster(min(cores[1]-1,numCores),type="PSOCK") # Not to overload
    cl <- makeCluster(min(cores[1]-1,numCores),type = "FORK") # Not to overload
    cat(sprintf("Using %d cores\n",min(cores[1]-1,numCores)))
    registerDoParallel(cl)
    cat(sprintf("%d cluster registered\n",min(cores[1]-1,numCores)))
    LL = function(lambda1,size1,lambda2,size2,sigma,epsilon){
      lambda1= exp(lambda1)
      size1 = exp(size1)
      lambda2= exp(lambda2)
      size2 = size1/(size.ratio+exp(size2))
      sigma=exp(sigma)
      epsilon=exp(epsilon)
      rate = lambda1*size1+lambda2*size2+sigma
      anc = reconstructAncestralStates(phy = phy,x=x[phy$tip.label],rate = rate,epsilon = epsilon)
      pp = parSapply(cl,bootstrap,function(i){
        dfunc(x = anc$contrast[i],t = anc$l[i],lambda1 = lambda1,size1=size1,lambda2 = lambda2,size2 = size2,
                        sigma = sigma,epsilon=anc$epsilon[i])
      })
      ll = -sum(log(pp))
      return(ll)
    }
  }else{
    LL = function(lambda1,size1,lambda2,size2,sigma,epsilon){
      lambda1= exp(lambda1)
      size1 = exp(size1)
      lambda2= exp(lambda2)
      size2 = size1/(size.ratio+exp(size2))
      sigma=exp(sigma)
      epsilon=exp(epsilon)
      rate = lambda1*size1+lambda2*size2+sigma
      anc = reconstructAncestralStates(phy = phy,x=x[phy$tip.label],rate = rate,epsilon = epsilon)
      pp = sapply(bootstrap,function(i){
        dfunc(x = anc$contrast[i],t = anc$l[i],lambda1 = lambda1,size1=size1,lambda2 = lambda2,size2 = size2,
                        sigma = sigma,epsilon=anc$epsilon[i])
      })
      ll = -sum(log(pp))
      return(ll)
    }
  }
  # if no starting parameters are supplied, set one
  if(is.null(start.value)) start.value = list(lambda1=log(1),size1=log(1),lambda2=log(1),size2=log(1),sigma = log(var(pic(x = x[phy$tip.label],phy = phy))),epsilon=log(1))
  # remove fixed parameters from the starting parameters
  start.value[names(fixed)]=NULL
  # set optimization method
  if(length(start.value)<2){
    method  = "BFGS"
  }else{
    method = "Nelder-Mead"
  }
  # evaluate AIC only if instructed
  if(eval.only){
    res = exp(unlist(c(start.value,fixed)))
    names(res) = c(names(start.value),names(fixed))
    res = res[c("lambda1","size1","lambda2","size2","sigma","epsilon")]
    res[4] = res[2]/(size.ratio+res[4])
    this.AIC = 2*do.call(LL,c(start.value,fixed))+2*length(start.value)
    if(numCores>1) stopCluster(cl)
    return(list(params=res,
                AIC=this.AIC,
                raw=NA))
  }
  # search for the best model parameters
  result = do.call("mle2", 
                   c(list(LL, 
                          start = start.value, 
                          data = list(x=x),
                          fixed = fixed,
                          method = method,
                          control = list(trace=TRUE,maxit=2000,reltol=reltol)
                   )))
  # transform model parameters back to the linear scale
  res = exp(result@fullcoef)
  res[4] = res[2]/(size.ratio+res[4])
  # stop parallel clusters
  if(numCores>1) stopCluster(cl)
  return(list(params=res,AIC=AIC(result),raw= result))
}

### function to simulate trait change between nodes
## l is a vector of branch lengths
## lambda, size, sigma and epsilon are parameters as defined in PEpois.R
## nj specifies the number of jumps between a specific pair of nodes, if supplied
# the output of this function is a list whose elements are vectors of trait change
# introduced by different processes
simulatePET = function(l,align.length=NA,lambda,size,sigma,epsilon,nj=NULL){
  # get the number of trait changes to simulate
  n = length(l)
  # adjust branch length if necessary (not used)
  if(is.na(align.length)){
    t=l
  }else{
    t = rgamma(n=n,shape=l*align.length+1,scale = 1/align.length)
  }
  # generate random number of jumps
  if(is.null(nj)) nj = rpois(n = n,lambda = lambda*t)
  # generate random trait change by each process
  dxbm = rnorm(n=n)*sqrt(t*sigma)
  dxpe = rnorm(n=n)*sqrt(nj*size)
  ep = runif(n = n)
  ebm = qnorm(p =ep)*sqrt(epsilon)
  elaplace = qlaplace(p = ep)*sqrt(epsilon/4)
  return(list(l = l, t = t,nj=nj,
              dxbm = dxbm, dxpe = dxpe,
              ep=ep,ebm = ebm, elaplace = elaplace))
}

### functions to simulate BM and PE1 trait value over phylogeny
## phy is a phylo-class object from ape package
## lambda, size, sigma and epsilon are parameters as defined in PEpois.R
## start is the trait value of the common ancestor (root)
# the output of this function is a vector of trait values
# BM and PE1 model
simulatePET.phy = function(phy,lambda,size,sigma,epsilon,start=0){
  # initialize trait values
  trait.sim = numeric(length = Ntip(phy)+Nnode(phy))+start
  # set node labels if not already exist
  if(is.null(phy$node.label)){
    names(trait.sim) = c(phy$tip.label,as.character(Ntip(phy)+(1:phy$Nnode)))
  }else{
    names(trait.sim) = c(phy$tip.label,phy$node.label)
  }
  # reorder edges for root-to-tip simulation 
  phy = reorder.phylo(phy,order = "postorder")
  # simulate trait changes between nodes
  trait.delta = simulatePET(l = phy$edge.length,lambda = lambda,size=size,sigma = sigma,epsilon = epsilon)
  # add up trait changes to get trait values from root to tip
  for(i in 1:length(phy$edge.length)){
    l = length(phy$edge.length)+1-i
    trait.sim[phy$edge[l,2]] = trait.sim[phy$edge[l,1]] + trait.delta$dxbm[l] + trait.delta$dxpe[l]
    # add time-independent variation for tips
    if(phy$edge[l,2]<=Ntip(phy)) trait.sim[phy$edge[l,2]] = trait.sim[phy$edge[l,2]] + trait.delta$elaplace[l]
  }
  return(trait.sim)
}

### functions to simulate PE2 trait value over phylogeny
## phy is a phylo-class object from ape package
## lambda1, size1, lambda2, size2, sigma and epsilon are parameters as defined in PEpois.R
## start is the trait value of the common ancestor (root)
# the output of this function is a vector of trait values
simulatePET2.phy = function(phy,lambda1,size1,lambda2,size2,sigma,epsilon,start=0){
  # initialize trait values
  trait.sim = numeric(length = Ntip(phy)+Nnode(phy))+start
  # set node labels if not already exist
  if(is.null(phy$node.label)){
    names(trait.sim) = c(phy$tip.label,as.character(Ntip(phy)+(1:phy$Nnode)))
  }else{
    names(trait.sim) = c(phy$tip.label,phy$node.label)
  }
  # reorder edges for root-to-tip simulation
  phy = reorder.phylo(phy,order = "postorder")
  # simulate trait changes between nodes
  trait.delta1 = simulatePET(l = phy$edge.length,lambda = lambda1,size=size1,sigma = sigma,epsilon = epsilon)
  trait.delta2 = simulatePET(l = phy$edge.length,lambda = lambda2,size=size2,sigma = 0,epsilon = 0)
  # add up trait changes to get trait values from root to tip
  for(i in 1:length(phy$edge.length)){
    l = length(phy$edge.length)+1-i
    trait.sim[phy$edge[l,2]] = trait.sim[phy$edge[l,1]] + trait.delta1$dxbm[l] + 
      trait.delta1$dxpe[l] + trait.delta2$dxpe[l]
    # add time-independent variation for tips
    if(phy$edge[l,2]<=Ntip(phy)) trait.sim[phy$edge[l,2]] = trait.sim[phy$edge[l,2]] + trait.delta1$elaplace[l]
  }
  return(trait.sim)
}

### functions to simulate PE3 trait value over phylogeny
## phy is a phylo-class object from ape package
## lambda1, size1, lambda2, size2, lambda3, size3, sigma and epsilon are parameters as defined in PEpois.R
## start is the trait value of the common ancestor (root)
# the output of this function is a vector of trait values
simulatePET3.phy = function(phy,lambda1,size1,lambda2,size2,lambda3,size3,sigma,epsilon,start=0,laplace=TRUE){
  # initialize trait values
  trait.sim = numeric(length = Ntip(phy)+Nnode(phy))+start
  # set node labels if not already exist
  if(is.null(phy$node.label)){
    names(trait.sim) = c(phy$tip.label,as.character(Ntip(phy)+(1:phy$Nnode)))
  }else{
    names(trait.sim) = c(phy$tip.label,phy$node.label)
  }
  # reorder edges for root-to-tip simulation
  phy = reorder.phylo(phy,order = "postorder")
  # simulate trait changes between nodes
  trait.delta1 = simulatePET(l = phy$edge.length,lambda = lambda1,size=size1,sigma = sigma,epsilon = epsilon)
  trait.delta2 = simulatePET(l = phy$edge.length,lambda = lambda2,size=size2,sigma = 0,epsilon = 0)
  trait.delta3 = simulatePET(l = phy$edge.length,lambda = lambda3,size=size3,sigma = 0,epsilon = 0)
  # add up trait changes to get trait values from root to tip
  for(i in 1:length(phy$edge.length)){
    l = length(phy$edge.length)+1-i
    trait.sim[phy$edge[l,2]] = trait.sim[phy$edge[l,1]] + trait.delta1$dxbm[l] + 
      trait.delta1$dxpe[l] + trait.delta2$dxpe[l] + trait.delta3$dxpe[l]
    # add time-independent variation for tips
    if(phy$edge[l,2]<=Ntip(phy)){
      if(laplace){
        trait.sim[phy$edge[l,2]] = trait.sim[phy$edge[l,2]] + trait.delta1$elaplace[l]
      }else{
        trait.sim[phy$edge[l,2]] = trait.sim[phy$edge[l,2]] + trait.delta1$ebm[l]
      }
    }
  }
  return(trait.sim)
}

### function to calculate the posterior probability that jump happens over a certain branch given the trait
## x is the trait change between nodes
## t is the total branch length between nodes
## lambda, size, sigma, epsilon is the parameters of PE model as defined in PEpois.R
## laplace determines if Laplace or normal distribution is used for time-independent variation
# output value of this function is a vector whose elements denotes the probability that no jumps happen between
# nodes, the most likely number of jumps happened between nodes and its probability, expected number of jumps happened
# given the trait, total probability density covered by the function, and the probability density of having the trait
posteriorJumpProbability = function(x,t,lambda,size,sigma,epsilon,laplace=TRUE,margin=1e-6){
  # forcing to use normal distribution if there is no time-independent variation
  if(epsilon<=0) laplace=FALSE
  # initialize expected number of jumps
  Ej=0
  # initialize probability density covered and the probability density of trait
  if(laplace){
    sumP = dconv.norm.laplace(x = x,sigma = sigma*t,epsilon = epsilon)*dpois(x = 0,lambda = lambda*t)
    pall = dPEpoislaplace(x = x,t = t,lambda = lambda,size = size,sigma = sigma,epsilon = epsilon,margin = margin)
  }else{
    sumP = dnorm(x = x,mean = 0,sd = sqrt(sigma*t+epsilon))*dpois(x = 0,lambda = lambda*t)
    pall = dPEpoisnorm(x = x,t = t,lambda = lambda,size = size,sigma = sigma,epsilon = epsilon,margin = margin)
  }
  # probability density given no jump
  p0 = sumP
  # initialize most likely number of jumps and its probability density
  maxP = sumP
  maxK = 0
  # set number of jumps to 1
  k=1
  # calculate the probability density of having the trait given there is 1 jumps
  if(laplace){
    deltaP = dconv.norm.laplace(x = x,sigma=sigma*t+k*size,epsilon = epsilon)*dpois(x = k,lambda = lambda*t)
  }else{
    deltaP = dnorm(x = x,mean = 0,sd = sqrt(sigma*t+k*size+epsilon))*dpois(x = k,lambda = lambda*t)
  }
  # check if we cover a sufficiently large proportion of probability density
  while((pall-sumP)>=margin){
    # update most likely number of jumps and its probability density
    if(deltaP>maxP){
      maxP = deltaP
      maxK = k
    }
    # update probability covered
    sumP = sumP+deltaP
    # update expected number of jumps
    Ej = Ej+deltaP*k
    # update number of jumps
    k = k+1
    # the probability density of having the trait given there is k jumps
    if(laplace){
      deltaP = dconv.norm.laplace(x = x,sigma=sigma*t+k*size,epsilon = epsilon)*dpois(x = k,lambda = lambda*t)
    }else{
      deltaP = dnorm(x = x,mean = 0,sd = sqrt(sigma*t+k*size+epsilon))*dpois(x = k,lambda = lambda*t)
    }
  }
  return(c(p0=p0/pall,k=maxK,pk=maxP/pall,Ej=Ej/pall,sumP=sumP,pall=pall))
}

### function to calculate CI from bootstrapping
## stat is the parameter from the empirical data
## bootstrap.stat are the parameters from the bootstrapped data
## alpha is the probability that the true value is outside the confidence interval
# output value of this function is a vector of length two denoting the lower and
# upper limits of the confidence interval
calculateBootstrapCI = function(stat,bootstrap.stat,alpha=0.05){
  # the difference between empirical and bootstrapped parameters
  delta = bootstrap.stat-stat
  # critical quantile of the difference
  bootstrap.q = quantile(delta,probs=c(1-alpha/2,alpha/2))
  # the confidence interval
  return(unname(stat-bootstrap.q))
}

### calculates the expected waiting time before a jump larger than a certain size occur
## lambda is the number of jumps per unit branch length
## size is the variance of jump sizes
## threshold is a vector of different jump sizes
## unitTime is the scale of the branch length
expectedWaitingTime = function(lambda,size,threshold,unitTime=1){
  waitTime = sapply(threshold,function(tt){
    # calculate the probability that a jump is larger than a certain size
    pp=2*pnorm(q = tt,mean = 0,sd = sqrt(size),lower.tail = FALSE)
    # calculate the expected waiting time
    return(unname(unitTime/sum(lambda*pp)))
  })
  return(waitTime)
}

### function to calculate the posterior probability that jump happens over a certain branch given the trait (PE2)
## x is the trait change between nodes
## t is the total branch length between nodes
## lambda1, lambda2, size1, size2, sigma, epsilon is the parameters of PE model as defined in PEpois.R
# output value of this function is a vector whose elements denotes the probability that no jumps happen between
# nodes, no jumps in the first Poisson process happen between the nodes, no jumps in the second Poisson process
# happen between the nodes, and the probability of having the traits
posteriorJumpProbability.laplace2 = function(x,t,lambda1,size1,lambda2,size2,sigma,epsilon,margin=1e-6){
  # the probability density of having no jumps
  p0 = dconv.norm.laplace(x = x,sigma = sigma*t,epsilon = epsilon)*dpois(x = 0,lambda = (lambda1+lambda2)*t)
  # get the range of jumps that will cover sufficiently large probability mass
  k1 = qpois(p = c(margin/2,1-margin/2),lambda = t*lambda1)
  k2 = qpois(p = c(margin/2,1-margin/2),lambda = t*lambda2)
  # the probability density of having no jumps in the first Poisson process
  p01 = sum(sapply(floor(k2[1]):ceiling(k2[2]),function(k){
    dconv.norm.laplace(x = x,sigma = size2*k+sigma*t,epsilon = epsilon)*dpois(x = 0,lambda = lambda1*t)*dpois(x=k,lambda=lambda2*t)
    }))
  # the probability density of having no jumps in the second Poisson process
  p02 = sum(sapply(floor(k1[1]):ceiling(k1[2]),function(k){
    dconv.norm.laplace(x = x,sigma = size1*k+sigma*t,epsilon = epsilon)*dpois(x = 0,lambda = lambda2*t)*dpois(x=k,lambda=lambda1*t)
  }))
  # the probability density of having the traits
  sumP =  dPEpois2laplace(x = x,t = t,lambda1 = lambda1,size1=size1,lambda2=lambda2,size2 = size2,sigma = sigma,epsilon = epsilon,margin=margin)
  return(c(p0=p0/sumP,p01=p01/sumP,p02=p02/sumP,sumP=sumP))
}

### function to calculate the posterior probability that jump happens over a certain branch given the trait (PE3)
## x is the trait change between nodes
## t is the total branch length between nodes
## lambda1, lambda2, lambda3, size1, size2, size3, sigma, epsilon is the parameters of PE model as defined in PEpois.R
# output value of this function is a vector whose elements denotes the probability that no jumps happen between
# nodes, no jumps in the first, the second, and the third Poisson process happen between the nodes, jumps only
# occur in the first, the second, and the third Poisson process between the nodes, and the probability of having the traits
posteriorJumpProbability.laplace3 = function(x,t,lambda1,size1,lambda2,size2,lambda3,size3,
                                             sigma,epsilon,margin=1e-6){
  # probability density of having no jumps
  p0 = dconv.norm.laplace(x = x,sigma = sigma*t,epsilon = epsilon)*dpois(x = 0,lambda = (lambda1+lambda2)*t)
  # probability density of having no jumps in the first Poisson process
  p01 = dPEpois2laplace(x = x,t = t,lambda1=lambda2,size1=size2,lambda2=lambda3,size2=size3,
                        sigma = sigma,epsilon = epsilon,margin = margin)*dpois(x = 0,lambda = lambda1*t)
  # probability density of having no jumps in the second Poisson process
  p02 = dPEpois2laplace(x = x,t = t,lambda1=lambda1,size1=size1,lambda2=lambda3,size2=size3,
                        sigma = sigma,epsilon = epsilon,margin = margin)*dpois(x = 0,lambda = lambda2*t)
  # probability density of having no jumps in the third Poisson process
  p03 = dPEpois2laplace(x = x,t = t,lambda1=lambda1,size1=size1,lambda2=lambda2,size2=size2,
                        sigma = sigma,epsilon = epsilon,margin = margin)*dpois(x = 0,lambda = lambda3*t)
  # probability density of having jumps only in the first Poisson process
  p11 = dPEpoislaplace(x = x,t = t,lambda = lambda1,size = size1,
                       sigma = sigma,epsilon = epsilon,margin = margin)*dpois(x = 0,lambda = (lambda2+lambda3)*t)
  # probability density of having jumps only in the second Poisson process
  p12 = dPEpoislaplace(x = x,t = t,lambda = lambda2,size = size2,
                       sigma = sigma,epsilon = epsilon,margin = margin)*dpois(x = 0,lambda = (lambda1+lambda3)*t)
  # probability density of having jumps only in the third Poisson process
  p13 = dPEpoislaplace(x = x,t = t,lambda = lambda3,size = size3,
                       sigma = sigma,epsilon = epsilon,margin = margin)*dpois(x = 0,lambda = (lambda1+lambda2)*t)
  # probability density of having the traits
  sumP =  dPEpois3laplace(x = x,t = t,lambda1 = lambda1,size1=size1,lambda2=lambda2,size2 = size2,lambda3=lambda3,size3=size3,
                          sigma = sigma,epsilon = epsilon,margin=margin)
  return(c(p0=p0/sumP,p01=p01/sumP,p02=p02/sumP,p03 = p03/sumP,
           p11=p11/sumP,p12=p12/sumP,p13=p13/sumP,sumP=sumP))
}

### function to calculate the relative distance to the root
## phy is a phylo-class object from the ape package
# output value of this function is a vector of relative distance to the root
relativeDepth.phy = function(phy){
  # calculate all pairwise distances
  dist.phy = dist.nodes(x = phy)
  # initialize relative distances
  relative.depth = numeric(Ntip(phy)+phy$Nnode)+1
  # for each node, calculate relative distance
  for(i in Ntip(phy)+1:phy$Nnode){
    # distance to the root
    d2root = dist.phy[i,Ntip(phy)+1]
    # find all tip descendants of the node
    tip.clade = extract.clade(phy = phy,node = i)
    # calculate the average distance from the node to the tips
    d2tip = dist.phy[i,match(tip.clade$tip.label,phy$tip.label)]
    # calculate the relative distance
    relative.depth[i] = d2root/(d2root+mean(d2tip))
  }
  return(relative.depth)
}

### function to summarize model parameters
## AIC is a vector of AIC values from the optimizer
## scale is the scale used to linearly transform the data
## sample.size is the number of nodes that is evaluated in the likelihood function
# output value of this function is a vector of corrected AIC values
# note that this function should not change the best model for any trait
summarize.model.parameter = function(params,time=2000,scale=1){
  # get the number of Poisson processes
  numPE = length(params)/2-1
  # correct for scaling
  unscale.coef = rep(scale,length(params))
  unscale.coef[2*(1:numPE)-1] = 1
  unscale.params = params/unscale.coef
  # calculating average waiting time
  wait.time = sapply(1:numPE,function(i){
    unname(time/params[2*i-1])
  })
  # calculate relative jump size
  relative.size = sapply(1:numPE,function(i){
    sqrt(unname(params[2*i]/params[2*numPE+2]))
  })
  # calculate jump rate per unit time
  jump.rate = 1/wait.time
  # calculate process variance (trait change variance per unit branch length)
  process.variance = sapply(1:numPE,function(i){
    params[2*i-1]*params[2*i]
  })
  # calculate relative contribution
  contribution = sapply(1:numPE,function(i){
    process.variance[i]/(sum(process.variance)+params[2*numPE+1])
  })
  return(list(summary=data.frame(time=wait.time,rate=jump.rate,size=relative.size,contribution = contribution),params=unscale.params))
}

### function to estimate stasis time
## pfunc is one of the cumulative probability function in PEpois.R
## params is the parameter of the pfunc
## alpha is the probability that trait is outside the confidence interval
## threshold is the maximum relative increase in confidence interval width before the stasis ends
## l.max and y.max provides a upper limit for searching the numerical solution
# output value of this function is the stasis time as branch length
estimate.stasis.period = function(params,pfunc,alpha=0.05,threshold=0.1,l.max=10,y.max=10){
  # define a function for solving cumulative probability equations
  this.p = function(q,t){
    (1-alpha/2)-do.call(what = pfunc,args = c(list(q=q,t=t),as.list(params)))
  }
  # calculate the confidence interval at zero-branch lengths
  base.root = uniroot(f = this.p,
                      interval = c(0,qlaplace(p = 1-alpha/4,mu = 0,sigma = sqrt(params[length(params)]/2))),
                      t=0)
  base.CI = base.root$root
  # define a function for finding the stasis breaking point
  this.CI = function(t){
    this.root = uniroot(f = this.p,
                        interval = c(0,y.max),
                        t=t)
    (1+threshold)*base.CI - this.root$root
  }
  # find the stasis breaking point
  CI.root = uniroot(f=this.CI,interval = c(0,l.max))
  return(CI.root$root)
}

### function to correct for the effect of scaling
## AIC is a vector of AIC values from the optimizer
## scale is the scale used to linearly transform the data
## sample.size is the number of nodes that is evaluated in the likelihood function
# output value of this function is a vector of corrected AIC values
# note that this function should not change the best model for any trait
normalize.AIC = function(AIC,scale=1,sample.size=6667){
  return(AIC-log(scale)*sample.size)
}

### function to calculate statistics like contribution
## params is the vector of parameters of PE models
# the output value of this function is a vector of parameters and other statistics
expand.model.parameter = function(params){
  # get the number of Poisson processes
  numPE = length(params)/2-1
  # calculate variance of trait change per unit branch length for each process
  process.variance = c(sapply(1:numPE,function(i){
    params[2*i-1]*params[2*i]
  }),params[2*numPE+1])
  # calculate relative contribution
  contribution = process.variance/sum(process.variance)
  # combine the results
  expand.params = c(params,process.variance,contribution)
  return(expand.params)
}
#
simulatePET2.phy.withTrack = function(phy,lambda1,size1,lambda2,size2,sigma,epsilon,start=0){
  # initialize trait values
  trait.sim = numeric(length = Ntip(phy)+Nnode(phy))+start
  numJump2 = numJump1 = numeric(length = Ntip(phy)+Nnode(phy))
  # set node labels if not already exist
  if(is.null(phy$node.label)){
    names(trait.sim) = c(phy$tip.label,as.character(Ntip(phy)+(1:phy$Nnode)))
  }else{
    names(trait.sim) = c(phy$tip.label,phy$node.label)
  }
  # reorder edges for root-to-tip simulation
  phy = reorder.phylo(phy,order = "postorder")
  # simulate trait changes between nodes
  trait.delta1 = simulatePET(l = phy$edge.length,lambda = lambda1,size=size1,sigma = sigma,epsilon = epsilon)
  trait.delta2 = simulatePET(l = phy$edge.length,lambda = lambda2,size=size2,sigma = 0,epsilon = 0)
  # add up trait changes to get trait values from root to tip
  for(l in rev(1:length(phy$edge.length))){
    trait.sim[phy$edge[l,2]] = trait.sim[phy$edge[l,1]] + trait.delta1$dxbm[l] + 
      trait.delta1$dxpe[l] + trait.delta2$dxpe[l]
    numJump1[phy$edge[l,2]] = trait.delta1$nj[l]
    numJump2[phy$edge[l,2]] = trait.delta2$nj[l]
  }
  return(list(heritable.trait=trait.sim,nj1=numJump1,nj2=numJump2,
              observed.trait = trait.sim+rlaplace(n = length(trait.sim),)*sqrt(epsilon/4)))
}
#
dVRgamma = function(x,t,prob=1,k,theta,epsilon,num.cat = 20,log=FALSE,laplace=TRUE){
  VR.bin = seq(0,1,length.out=num.cat+1)
  VR.rate = c(qgamma(p = (VR.bin[-1]+VR.bin[-(num.cat+1)])/2,shape = k,scale = theta),k*theta)
  VR.priors = c(rep(prob/num.cat,num.cat),1-prob)
  if(laplace){
    VR.pp = sapply(VR.rate,function(rr){
      dconv.norm.laplace(x = x,sigma = rr*t,epsilon = epsilon)
      })
  }else{
    VR.pp=dnorm(x = x,mean = 0,sd = sqrt(VR.rate*t+epsilon))
  }
  VR.p = sum(VR.pp*VR.priors)
  if(log) return(log(VR.p))
  return(VR.p)
}
pVRgamma = function(q,t,prob=1,k,theta,epsilon,num.cat = 20,log=FALSE,laplace=TRUE){
  VR.bin = seq(0,1,length.out=num.cat+1)
  VR.rate = c(qgamma(p = (VR.bin[-1]+VR.bin[-(num.cat+1)])/2,shape = k,scale = theta),k*theta)
  VR.priors = c(rep(prob/num.cat,num.cat),1-prob)
  if(laplace){
    VR.pp = sapply(VR.rate,function(rr){
      pconv.norm.laplace(q = q,sigma = rr*t,epsilon = epsilon)
    })
  }else{
    VR.pp=pnorm(q = q,mean = 0,sd = sqrt(VR.rate*t+epsilon))
  }
  VR.p = sum(VR.pp*VR.priors)
  if(log) return(log(VR.p))
  return(VR.p)
}
#
fitPE.phy.gamma = function(phy,x,start.value=NULL,numCores=1,fixed=list(),ignore.zero=TRUE,
                           laplace=TRUE,eval.only=FALSE,bootstrap=NULL,reltol=sqrt(.Machine$double.eps),...){
  # initialize node indexes if not supplied
  if(is.null(bootstrap)) bootstrap = 1:phy$Nnode
  # remove zero-contrasts if instructed
  if(ignore.zero) bootstrap = intersect(bootstrap,
                                        which(reconstructAncestralStates(phy = phy,x=x[phy$tip.label],rate = 1,epsilon = 0)$contrast!=0))
  # set probability function to use
  dfunc = dVRgamma
  LL = function(prob,k,theta,epsilon){
    prob=sigmoid(prob)
    k= exp(k)
    theta=exp(theta)
    epsilon=exp(epsilon)
    rate = k*theta
    anc = reconstructAncestralStates(phy = phy,x=x[phy$tip.label],rate = rate,epsilon = epsilon)
    pp = do.call(c,mclapply(bootstrap,function(i){
      dfunc(x = anc$contrast[i],t = anc$l[i],k=k,theta=theta,epsilon=anc$epsilon[i],prob=prob,laplace=laplace)
    },mc.cores = numCores))
    ll = -sum(log(pp))
    return(ll)
  }
  # if no starting parameters are supplied, set one
  if(is.null(start.value)) start.value = list(prob=logit(0.99),k=log(1),theta=log(var(pic(x = x[phy$tip.label],phy = phy))),epsilon=log(1))
  # remove fixed parameters from the starting parameters
  start.value[names(fixed)]=NULL
  # set optimization method
  if(length(start.value)<2){
    method  = "BFGS"
  }else{
    method = "Nelder-Mead"
  }
  # evaluate AIC only if instructed
  if(eval.only){
    res = exp(unlist(c(start.value,fixed)))
    names(res) = c(names(start.value),names(fixed))
    res["prob"] = sigmoid(log(res["prob"]))
    this.AIC = 2*do.call(LL,c(start.value,fixed))+2*length(start.value)
    return(list(params=res,
                AIC=this.AIC,
                raw=NA))
  }
  # search for the best model parameters
  result = do.call("mle2", 
                   c(list(LL, 
                          start = start.value, 
                          data = list(x=x),
                          fixed = fixed,
                          method = method,
                          control = list(trace=TRUE,maxit=2000,reltol=reltol)
                   )))
  # transform model parameters back to the linear scale
  res = exp(result@fullcoef)
  res["prob"] = sigmoid(log(res["prob"]))
  # stop parallel clusters
  return(list(params=res,AIC=AIC(result),raw= result))
}
#
simulateGamma.phy = function(tree,prob,k,theta,epsilon,start= 0){
  node.idx = Ntip(tree)+(1:Nnode(tree))
  node.rate = rgamma(n = length(node.idx),shape = k,scale = theta)
  node.var = rbinom(n = length(node.idx),size = 1,prob = prob)
  new.tree = tree
  for(i in 1:length(node.idx)){
    nn = node.idx[[i]]
    edge.idx = which(new.tree$edge[,1]==nn)
    if(node.var[i]>0){
      new.tree$edge.length[edge.idx] = tree$edge.length[edge.idx]*node.rate[i]
    }else{
      new.tree$edge.length[edge.idx] = tree$edge.length[edge.idx]*k*theta
    }
  }
  sim.trait = simulateBMTrait(phy = new.tree,start=start)
  obs.trait = sim.trait + rlaplace(n = length(sim.trait))*sqrt(epsilon/4)
  return(list(heritable.trait=sim.trait,observed.trait=obs.trait,phy=new.tree))
}
#
simulatePET3.phy.withTrack = function(phy,lambda1,size1,lambda2,size2,lambda3,size3,sigma,epsilon,start=0){
  # initialize trait values
  trait.sim = numeric(length = Ntip(phy)+Nnode(phy))+start
  numJump3 = numJump2 = numJump1 = numeric(length = Ntip(phy)+Nnode(phy))
  # set node labels if not already exist
  if(is.null(phy$node.label)){
    names(trait.sim) = c(phy$tip.label,as.character(Ntip(phy)+(1:phy$Nnode)))
  }else{
    names(trait.sim) = c(phy$tip.label,phy$node.label)
  }
  # reorder edges for root-to-tip simulation
  phy = reorder.phylo(phy,order = "postorder")
  # simulate trait changes between nodes
  trait.delta1 = simulatePET(l = phy$edge.length,lambda = lambda1,size=size1,sigma = sigma,epsilon = epsilon)
  trait.delta2 = simulatePET(l = phy$edge.length,lambda = lambda2,size=size2,sigma = 0,epsilon = 0)
  trait.delta3 = simulatePET(l = phy$edge.length,lambda = lambda3,size=size3,sigma = 0,epsilon = 0)
  
  # add up trait changes to get trait values from root to tip
  for(l in rev(1:length(phy$edge.length))){
    trait.sim[phy$edge[l,2]] = trait.sim[phy$edge[l,1]] + trait.delta1$dxbm[l] + 
      trait.delta1$dxpe[l] + trait.delta2$dxpe[l] + trait.delta3$dxpe[l]
    numJump1[phy$edge[l,2]] = trait.delta1$nj[l]
    numJump2[phy$edge[l,2]] = trait.delta2$nj[l]
    numJump3[phy$edge[l,2]] = trait.delta3$nj[l]
  }
  return(list(heritable.trait=trait.sim,nj1=numJump1,nj2=numJump2,nj3 = numJump3,
              observed.trait = trait.sim+rlaplace(n = length(trait.sim),)*sqrt(epsilon/4)))
}
#
```
