## Supplementary material for "Microbial genomic trait evolution is dominated by frequent and rare pulsed evolution": PEpois.R

```
#
library(pracma)
library(extraDistr)
# DPR functions for 1PE model with normal residuals
dPEpoisnorm = function(x,t,lambda,size,sigma,epsilon=0,margin=1e-6){
  En = lambda*t
  if(En>1e4){
    p = dnorm(x = x,mean = 0,sd = sqrt(size*En+sigma*t+epsilon))
  }else{
    k = qpois(p = c(margin/2,1-margin/2),lambda = En)
    k= floor(k[1]):ceiling(k[2])
    probs = dpois(x = k,lambda = En)
    p = sum(dnorm(x = x,mean = 0,sd = sqrt(size*k+sigma*t+epsilon))*probs) 
  }
  return(p)
}
pPEpoisnorm = function(q,t,lambda,size,sigma,epsilon=0,margin=1e-6,lower.tail=TRUE){
  En = lambda*t
  if(En>1e4){
    p = pnorm(q = q,mean = 0,sd = sqrt(size*En+sigma*t+epsilon),lower.tail = lower.tail)
  }else{
    k = qpois(p = c(margin/2,1-margin/2),lambda = En)
    k= floor(k[1]):ceiling(k[2])
    probs = dpois(x = k,lambda = En)
    p = sum(pnorm(q = q,mean = 0,sd = sqrt(size*k+sigma*t+epsilon),lower.tail = lower.tail)*probs) 
  }
  return(p)
}
rPEpoisnorm = function(n,t,lambda,size,sigma,epsilon=0){
  En = lambda*t
  k = rpois(n = n,lambda = En)
  sds = sqrt(k*size+sigma*t+epsilon)
  x = rnorm(n = n,mean = 0,sd = sds)
  return(x)
}
# DPR functions for 2PE model with normal residuals
dPEpois2norm = function(x,t,lambda1,lambda2,size1,size2,sigma,epsilon,margin=1e-6){
  En1 = lambda1*t
  En2 = lambda2*t
  if(En1>1e2){
    p = dPEpoisnorm(x = x,t = t, lambda = lambda2, size = size2, sigma = size1*lambda1+sigma, epsilon = epsilon,margin=margin)
  }else{
    if(En2>1e2){
      p = dPEpoisnorm(x = x,t = t, lambda = lambda1, size = size1, sigma = size2*lambda2+sigma, epsilon = epsilon,margin=margin)
    }else{
      k1 = qpois(p = c(margin/2,1-margin/2),lambda = En1)
      k1= floor(k1[1]):ceiling(k1[2])
      k2 = qpois(p = c(margin/2,1-margin/2),lambda = En2)
      k2= floor(k2[1]):ceiling(k2[2])
      probs1 = dpois(x = k1,lambda = En1)
      probs2 = dpois(x = k2,lambda = En2)
      p = sapply(1:length(probs1),function(i){
        sum(sapply(k2,function(kk2){
          dnorm(x = x,sd = sqrt(size1*k1[i]+size2*kk2+sigma*t+epsilon))
        })*probs2)*probs1[i]
      })
    }
  }
  return(sum(p))
}
pPEpois2norm = function(q,t,lambda1,lambda2,size1,size2,sigma,epsilon,margin=1e-6,lower.tail=TRUE){
  En1 = lambda1*t
  En2 = lambda2*t
  if(En1>1e2){
    p = pPEpoisnorm(q = q,t = t, lambda = lambda2, size = size2, sigma = size1*lambda1+sigma, epsilon = epsilon,margin=margin,lower.tail = lower.tail)
  }else{
    if(En2>1e2){
      p = pPEpoisnorm(q = q,t = t, lambda = lambda1, size = size1, sigma = size2*lambda2+sigma, epsilon = epsilon,margin=margin,lower.tail = lower.tail)
    }else{
      k1 = qpois(p = c(margin/2,1-margin/2),lambda = En1)
      k1= floor(k1[1]):ceiling(k1[2])
      k2 = qpois(p = c(margin/2,1-margin/2),lambda = En2)
      k2= floor(k2[1]):ceiling(k2[2])
      probs1 = dpois(x = k1,lambda = En1)
      probs2 = dpois(x = k2,lambda = En2)
      p = sapply(1:length(probs1),function(i){
        sum(sapply(k2,function(kk2){
          pnorm(q = q,sd = sqrt(size1*k1[i]+size2*kk2+sigma*t+epsilon),lower.tail = lower.tail)
        })*probs2)*probs1[i]
      })
    }
  }
  return(sum(p))
}
rPEpois2norm = function(n,t,lambda1,lambda2,size1,size2,sigma,epsilon){
  En1 = lambda1*t
  En2 = lambda2*t
  k1 = rpois(n = n,lambda = En1)
  k2 = rpois(n = n,lambda = En2)
  sds = sqrt(k1*size1 + k2*size2 +sigma*t + epsilon)
  x = rnorm(n = n, mean = 0, sd = sds)
  return(x)
}
# DPR functions for 1PE model with Laplace residuals
dPEpoislaplace = function(x,t,lambda,size,sigma,epsilon,margin=1e-6){
  En = lambda*t
  if(En>1e4){
    p = dconv.norm.laplace(x = x,sigma = size*En+sigma*t,epsilon = epsilon)
  }else{
    k = qpois(p = c(margin/2,1-margin/2),lambda = En)
    k= floor(k[1]):ceiling(k[2])
    probs = dpois(x = k,lambda = En)
    p = sapply(k,function(kk){dconv.norm.laplace(x = x,sigma = size*kk+sigma*t,epsilon = epsilon)})*probs
  }
  return(sum(p))
}
pPEpoislaplace = function(q,t,lambda,size,sigma,epsilon,margin=1e-6,lower.tail=TRUE){
  En = lambda*t
  if(En>1e4){
    p = pconv.norm.laplace(q = q,sigma = size*En+sigma*t,epsilon = epsilon,lower.tail=lower.tail)
  }else{
    k = qpois(p = c(margin/2,1-margin/2),lambda = En)
    k= floor(k[1]):ceiling(k[2])
    probs = dpois(x = k,lambda = En)
    p = sapply(k,function(kk){
      pconv.norm.laplace(q = q,sigma = size*kk+sigma*t,epsilon = epsilon,lower.tail=lower.tail)
    })*probs
  }
  return(sum(p))
}
rPEpoislaplace = function(n,t,lambda,size,sigma,epsilon=0){
  En = lambda*t
  k = rpois(n = n,lambda = En)
  sds = sqrt(k*size+sigma*t)
  x = rnorm(n = n,mean = 0,sd = sds)+rlaplace(n=n,mu=0,sigma=sqrt(epsilon/2))
  return(x)
}
# DPR functions for 2PE model with Laplace residuals
dPEpois2laplace = function(x,t,lambda1,lambda2,size1,size2,sigma,epsilon,margin=1e-6){
  En1 = lambda1*t
  En2 = lambda2*t
  if(En1>1e2){
    p = dPEpoislaplace(x = x,t = t, lambda = lambda2, size = size2, sigma = size1*lambda1+sigma, epsilon = epsilon,margin=margin)
  }else{
    if(En2>1e2){
      p = dPEpoislaplace(x = x,t = t, lambda = lambda1, size = size1, sigma = size2*lambda2+sigma, epsilon = epsilon,margin=margin)
    }else{
      k1 = qpois(p = c(margin/2,1-margin/2),lambda = En1)
      k1= floor(k1[1]):ceiling(k1[2])
      k2 = qpois(p = c(margin/2,1-margin/2),lambda = En2)
      k2= floor(k2[1]):ceiling(k2[2])
      probs1 = dpois(x = k1,lambda = En1)
      probs2 = dpois(x = k2,lambda = En2)
      p = sapply(1:length(probs1),function(i){
        sum(sapply(k2,function(kk2){
          dconv.norm.laplace(x = x,sigma = size1*k1[i]+size2*kk2+sigma*t,epsilon = epsilon)
        })*probs2)*probs1[i]
      })
    }
  }
  return(sum(p))
}
pPEpois2laplace = function(q,t,lambda1,lambda2,size1,size2,sigma,epsilon,margin=1e-6,lower.tail=TRUE){
  En1 = lambda1*t
  En2 = lambda2*t
  if(En1>1e2){
    p = pPEpoislaplace(q = q,t = t, lambda = lambda2, size = size2, sigma = size1*lambda1+sigma, epsilon = epsilon,
                       margin=margin,lower.tail = lower.tail)
  }else{
    if(En2>1e2){
      p = pPEpoislaplace(q = q,t = t, lambda = lambda1, size = size1, sigma = size2*lambda2+sigma, epsilon = epsilon,
                         margin=margin,lower.tail=lower.tail)
    }else{
      k1 = qpois(p = c(margin/2,1-margin/2),lambda = En1)
      k1= floor(k1[1]):ceiling(k1[2])
      k2 = qpois(p = c(margin/2,1-margin/2),lambda = En2)
      k2= floor(k2[1]):ceiling(k2[2])
      probs1 = dpois(x = k1,lambda = En1)
      probs2 = dpois(x = k2,lambda = En2)
      p = sapply(1:length(probs1),function(i){
        sum(sapply(k2,function(kk2){
          pconv.norm.laplace(q = q,sigma = size1*k1[i]+size2*kk2+sigma*t,epsilon = epsilon,lower.tail=lower.tail)
        })*probs2)*probs1[i]
      })
    }
  }
  return(sum(p))
}
rPEpois2laplace = function(n,t,lambda1,lambda2,size1,size2,sigma,epsilon){
  En1 = lambda1*t
  En2 = lambda2*t
  k1 = rpois(n = n,lambda = En1)
  k2 = rpois(n = n,lambda = En2)
  sds = sqrt(k1*size1 + k2*size2 +sigma*t)
  x = rnorm(n = n, mean = 0, sd = sds)+rlaplace(n = n,mu = 0,sigma = sqrt(epsilon/2))
  return(x)
}
#
#
dconv.norm.laplace = function(x,sigma,epsilon){
  if(sigma>(1e6*epsilon)) return(dnorm(x = x,mean = 0,sd = sqrt(sigma+epsilon)))
  scale = 2/epsilon
  x=x*sqrt(scale)
  sigma=sigma*scale
  p=exp(x+sigma/2+pnorm(q = x,mean = -sigma,sd = sqrt(sigma),lower.tail = FALSE,log.p = TRUE))+
    exp(sigma/2-x+pnorm(q = x,mean = sigma,sd = sqrt(sigma),log.p = TRUE))
  return(sqrt(scale)*p/2)
}
pconv.norm.laplace = function(q,sigma,epsilon,lower.tail=TRUE){
  if(sigma>(1e6*epsilon)) return(pnorm(q = q,mean = 0,sd = sqrt(sigma+epsilon),lower.tail = lower.tail))
  scale = 2/epsilon
  q=q*sqrt(scale)
  sigma=sigma*scale
  p = pnorm(q = q,mean = 0,sd = sqrt(sigma),lower.tail = TRUE)+
    exp(q+sigma/2+pnorm(q = q,mean = -sigma,sd = sqrt(sigma),lower.tail = FALSE,log.p = TRUE))/2-
    exp(sigma/2-q+pnorm(q = q,mean = sigma,sd = sqrt(sigma),log.p = TRUE))/2
  if(lower.tail){
    return(p)
  }else{
    return(1-p)
  }
}
#
dPEpois3norm = function(x,t,lambda1,lambda2,lambda3,size1,size2,size3,sigma,epsilon,margin=1e-6){
  En1 = lambda1*t
  En2 = lambda2*t
  En3 = lambda3*t
  if(En1>1e2){
    p = dPEpois2norm(x = x,t = t, lambda1=lambda2, size1=size2, lambda2=lambda3, size2=size3, sigma=size1*lambda1+sigma, epsilon = epsilon,margin=margin)
  }else{
    if(En2>1e2){
      p = dPEpois2norm(x = x,t = t, lambda1=lambda1, size1=size1, lambda2=lambda3, size2=size3, sigma=size2*lambda2+sigma, epsilon = epsilon,margin=margin)
    }else{
      if(En3>1e2){
        p = dPEpois2norm(x = x,t = t, lambda1=lambda1, size1=size1, lambda2=lambda2, size2=size2, sigma=size3*lambda3+sigma, epsilon = epsilon,margin=margin)
      }else{
        k1 = qpois(p = c(margin/2,1-margin/2),lambda = En1)
        k1= floor(k1[1]):ceiling(k1[2])
        k2 = qpois(p = c(margin/2,1-margin/2),lambda = En2)
        k2= floor(k2[1]):ceiling(k2[2])
        k3 = qpois(p = c(margin/2,1-margin/2),lambda = En3)
        k3= floor(k3[1]):ceiling(k3[2])
        probs1 = dpois(x = k1,lambda = En1)
        probs2 = dpois(x = k2,lambda = En2)
        probs3 = dpois(x = k3,lambda = En3)
        p = sum(sapply(1:length(probs1),function(i){
          sum(sapply(1:length(probs2),function(j){
            sum(sapply(k3,function(kk3){
              dnorm(x = x,sd = sqrt(size1*k1[i]+size2*k2[j]+size3*kk3+sigma*t+epsilon))
            })*probs3)*probs1[i]*probs2[j]
          }))
        }))
      }
    }
  }
  return(p)
}
pPEpois3norm = function(q,t,lambda1,lambda2,lambda3,size1,size2,size3,sigma,epsilon,margin=1e-6,lower.tail=TRUE){
  En1 = lambda1*t
  En2 = lambda2*t
  En3 = lambda3*t
  if(En1>1e2){
    p = pPEpois2norm(q = q,t = t,
                     lambda1=lambda2, size1=size2, lambda2=lambda3, size2=size3, sigma=size1*lambda1+sigma, epsilon=epsilon,
                     margin=margin,lower.tail = lower.tail)
  }else{
    if(En2>1e2){
      p = pPEpois2norm(q = q,t = t,
                       lambda1=lambda1, size1=size1, lambda2=lambda3, size2=size3, sigma=size2*lambda2+sigma, epsilon=epsilon,
                       margin=margin,lower.tail = lower.tail)
    }else{
      if(En3>1e2){
        p = pPEpois2norm(q = q,t = t,
                         lambda1=lambda1, size1=size1, lambda2=lambda2, size2=size2, sigma=size3*lambda3+sigma, epsilon=epsilon,
                         margin=margin,lower.tail = lower.tail)
      }else{
        k1 = qpois(p = c(margin/2,1-margin/2),lambda = En1)
        k1= floor(k1[1]):ceiling(k1[2])
        k2 = qpois(p = c(margin/2,1-margin/2),lambda = En2)
        k2= floor(k2[1]):ceiling(k2[2])
        k3 = qpois(p = c(margin/2,1-margin/2),lambda = En3)
        k3= floor(k3[1]):ceiling(k3[2])
        probs1 = dpois(x = k1,lambda = En1)
        probs2 = dpois(x = k2,lambda = En2)
        probs3 = dpois(x = k3,lambda = En3)
        p = sum(sapply(1:length(probs1),function(i){
          sum(sapply(1:length(probs2),function(j){
            sum(sapply(k3,function(kk3){
              pnorm(q = q,sd = sqrt(size1*k1[i]+size2*k2[j]+size3*kk3+sigma*t+epsilon))
            })*probs3)*probs1[i]*probs2[j]
          }))
        }))
      }
    }
  }
  return(p)
}
rPEpois3norm = function(n,t,lambda1,lambda2,lambda3,size1,size2,size3,sigma,epsilon){
  En1 = lambda1*t
  En2 = lambda2*t
  En3 = lambda3*t
  k1 = rpois(n = n,lambda = En1)
  k2 = rpois(n = n,lambda = En2)
  k3 = rpois(n = n,lambda = En3)
  sds = sqrt(k1*size1 + k2*size2 + k3*size3 +sigma*t + epsilon)
  x = rnorm(n = n, mean = 0, sd = sds)
  return(x)
}
#
dPEpois3laplace = function(x,t,lambda1,lambda2,lambda3,size1,size2,size3,sigma,epsilon,margin=1e-6){
  En1 = lambda1*t
  En2 = lambda2*t
  En3 = lambda3*t
  if(En1>1e2){
    p = dPEpois2laplace(x = x,t = t, lambda1=lambda2, size1=size2, lambda2=lambda3, size2=size3, sigma=size1*lambda1+sigma, epsilon = epsilon,margin=margin)
  }else{
    if(En2>1e2){
      p = dPEpois2laplace(x = x,t = t, lambda1=lambda1, size1=size1, lambda2=lambda3, size2=size3, sigma=size2*lambda2+sigma, epsilon = epsilon,margin=margin)
    }else{
      if(En3>1e2){
        p = dPEpois2laplace(x = x,t = t, lambda1=lambda1, size1=size1, lambda2=lambda2, size2=size2, sigma=size3*lambda3+sigma, epsilon = epsilon,margin=margin)
      }else{
        k1 = qpois(p = c(margin/2,1-margin/2),lambda = En1)
        k1= floor(k1[1]):ceiling(k1[2])
        k2 = qpois(p = c(margin/2,1-margin/2),lambda = En2)
        k2= floor(k2[1]):ceiling(k2[2])
        k3 = qpois(p = c(margin/2,1-margin/2),lambda = En3)
        k3= floor(k3[1]):ceiling(k3[2])
        probs1 = dpois(x = k1,lambda = En1)
        probs2 = dpois(x = k2,lambda = En2)
        probs3 = dpois(x = k3,lambda = En3)
        p = sum(sapply(1:length(probs1),function(i){
          sum(sapply(1:length(probs2),function(j){
            sum(sapply(k3,function(kk3){
              dconv.norm.laplace(x = x,sigma = size1*k1[i]+size2*k2[j]+size3*kk3+sigma*t,epsilon = epsilon)
            })*probs3)*probs1[i]*probs2[j]
          }))
        }))
      }
    }
  }
  return(p)
}
pPEpois3laplace = function(q,t,lambda1,lambda2,lambda3,size1,size2,size3,sigma,epsilon,margin=1e-6,lower.tail=TRUE){
  En1 = lambda1*t
  En2 = lambda2*t
  En3 = lambda3*t
  if(En1>1e2){
    p = pPEpois2laplace(q = q,t = t,
                     lambda1=lambda2, size1=size2, lambda2=lambda3, size2=size3, sigma=size1*lambda1+sigma, epsilon=epsilon,
                     margin=margin,lower.tail = lower.tail)
  }else{
    if(En2>1e2){
      p = pPEpois2laplace(q = q,t = t,
                       lambda1=lambda1, size1=size1, lambda2=lambda3, size2=size3, sigma=size2*lambda2+sigma, epsilon=epsilon,
                       margin=margin,lower.tail = lower.tail)
    }else{
      if(En3>1e2){
        p = pPEpois2laplace(q = q,t = t,
                         lambda1=lambda1, size1=size1, lambda2=lambda2, size2=size2, sigma=size3*lambda3+sigma, epsilon=epsilon,
                         margin=margin,lower.tail = lower.tail)
      }else{
        k1 = qpois(p = c(margin/2,1-margin/2),lambda = En1)
        k1= floor(k1[1]):ceiling(k1[2])
        k2 = qpois(p = c(margin/2,1-margin/2),lambda = En2)
        k2= floor(k2[1]):ceiling(k2[2])
        k3 = qpois(p = c(margin/2,1-margin/2),lambda = En3)
        k3= floor(k3[1]):ceiling(k3[2])
        probs1 = dpois(x = k1,lambda = En1)
        probs2 = dpois(x = k2,lambda = En2)
        probs3 = dpois(x = k3,lambda = En3)
        p = sum(sapply(1:length(probs1),function(i){
          sum(sapply(1:length(probs2),function(j){
            sum(sapply(k3,function(kk3){
              pconv.norm.laplace(q = q,sigma = size1*k1[i]+size2*k2[j]+size3*kk3+sigma*t,epsilon = epsilon,lower.tail=lower.tail)
            })*probs3)*probs1[i]*probs2[j]
          }))
        }))
      }
    }
  }
  return(p)
}
rPEpois3laplace = function(n,t,lambda1,lambda2,lambda3,size1,size2,size3,sigma,epsilon){
  En1 = lambda1*t
  En2 = lambda2*t
  En3 = lambda3*t
  k1 = rpois(n = n,lambda = En1)
  k2 = rpois(n = n,lambda = En2)
  k3 = rpois(n = n,lambda = En3)
  sds = sqrt(k1*size1 + k2*size2 + k3*size3 +sigma*t)
  x = rnorm(n = n, mean = 0, sd = sds)+rlaplace(n = n,mu = 0,sigma = sqrt(epsilon/2))
  return(x)
}
```
