## Supplementary material for "Microbial genomic trait evolution is dominated by frequent and rare pulsed evolution": seg_lm.R

```
# segmented linear regression
#
seg.1.lm = function(l1,l,dx,baseline=1e-3){
  tdx = log(abs(dx)+baseline)
  ll1 = log(l[l<=l1])
  tdx1 = tdx[l<=l1]
  lm1 = lm(tdx1~ll1)
  y1 = log(l1)*lm1$coefficients[2]+lm1$coefficients[1]
  ll2 = log(l[l>l1])-log(l1)
  tdx2 = tdx[l>l1]-y1
  lm2 = lm(tdx2~ll2+0)
  return(list(model=list(y0=lm1$coefficients[1],k1=lm1$coefficients[2],l1=l1,y1=y1,k2=lm2$coefficients[1]),
              aic = AIC(lm1)+AIC(lm2)+2))
}
seg.1.lm.fix = function(l1,l,dx,baseline=1e-3){
  tdx = log(abs(dx)+baseline)
  ll1 = log(l[l<=l1])
  tdx1 = tdx[l<=l1]
  lm1 = lm(tdx1~1)
  y1 = lm1$coefficients[1]
  ll2 = log(l[l>l1])-log(l1)
  tdx2 = tdx[l>l1]-y1
  lm2 = lm(tdx2~ll2+0)
  return(list(model=list(y0=lm1$coefficients[1],k1=0,l1=l1,y1=y1,k2=lm2$coefficients[1]),
              aic = AIC(lm1)+AIC(lm2)+2))
}
seg.2.lm = function(l1,l2,l,dx,baseline=1e-3){
  tdx = log(abs(dx)+baseline)
  ll1 = log(l[l<=l1])
  tdx1 = tdx[l<=l1]
  lm1 = lm(tdx1~ll1)
  y1 = log(l1)*lm1$coefficients[2]+lm1$coefficients[1]
  ll2 = log(l[l>l1&l<=l2])-log(l1)
  tdx2 = tdx[l>l1&l<=l2]-y1
  lm2 = lm(tdx2~ll2+0)
  y2 = (log(l2)-log(l1))*lm2$coefficients[1]+y1
  ll3 = log(l[l>l2])-log(l2)
  tdx3 = tdx[l>l2]-y2
  lm3 = lm(tdx3~ll3+0)
  return(list(model=list(y0=lm1$coefficients[1],
                         k1=lm1$coefficients[2],l1=l1,y1=y1,
                         k2=lm2$coefficients[1],l2=l2,y2=y2,
                         k3 = lm3$coefficients[1]),
              aic = AIC(lm1)+AIC(lm2)+AIC(lm3)+4))
}
seg.2.lm.fix = function(l1,l2,l,dx,baseline=1e-3){
  tdx = log(abs(dx)+baseline)
  ll1 = log(l[l<=l1])
  tdx1 = tdx[l<=l1]
  lm1 = lm(tdx1~1)
  y1 = lm1$coefficients[1]
  ll2 = log(l[l>l1&l<=l2])-log(l1)
  tdx2 = tdx[l>l1&l<=l2]-y1
  lm2 = lm(tdx2~ll2+0)
  y2 = (log(l2)-log(l1))*lm2$coefficients[1]+y1
  ll3 = log(l[l>l2])-log(l2)
  tdx3 = tdx[l>l2]-y2
  lm3 = lm(tdx3~ll3+0)
  return(list(model=list(y0=lm1$coefficients[1],
                         k1=0,l1=l1,y1=y1,
                         k2=lm2$coefficients[1],l2=l2,y2=y2,
                         k3 = lm3$coefficients[1]),
              aic = AIC(lm1)+AIC(lm2)+AIC(lm3)+4))
}
find.best.1.break = function(l,dx){
  ll = function(l1,l,dx){
    seg.1.lm(l1 = l1,l = l,dx = dx)$aic
  }
  best.break = optim(par = median(l),fn = ll,method = "Brent",lower = quantile(unique(l),probs = 0.025),upper = quantile(unique(l),probs = 0.975),l=l,dx=dx)
  best.lm = seg.1.lm(l = l,dx = dx,l1 = best.break$par)
  return(list(l1 = best.break$par,aic = best.break$value,model = best.lm$model))
}
find.best.1.break.fix = function(l,dx){
  ll = function(l1,l,dx){
    seg.1.lm.fix(l1 = l1,l = l,dx = dx)$aic
  }
  best.break = optim(par = median(l),fn = ll,method = "Brent",lower = quantile(unique(l),probs = 0.025),upper = quantile(unique(l),probs = 0.975),l=l,dx=dx)
  best.lm = seg.1.lm.fix(l = l,dx = dx,l1 = best.break$par)
  return(list(l1 = best.break$par,aic = best.break$value,model = best.lm$model))
}
find.best.2.break = function(l,dx){
  ll = function(rl,l,dx){
    l1 = quantile(unique(l),probs = rl[1])
    l2 = quantile(unique(l),probs = rl[1]+rl[2]*(1-rl[1]))
    seg.2.lm(l1 = l1,l2=l2,l = l,dx = dx)$aic
  }
  best.break = optim(par = c(0.5,0.5),fn = ll,method = "L-BFGS-B",lower = c(0.025,0.025),upper = c(0.975,0.975),l=l,dx=dx)
  l1=quantile(unique(l),probs = best.break$par[1])
  l2=quantile(unique(l),probs = best.break$par[1]+best.break$par[2]*(1-best.break$par[1]))
  best.lm = seg.2.lm(l = l,dx = dx,l1 = l1,l2 = l2)
  return(list(l1 = l1,l2 = l2,aic = best.break$value,model=best.lm$model))
}
find.best.2.break.fix = function(l,dx){
  ll = function(rl,l,dx){
    l1 = quantile(unique(l),probs = rl[1])
    l2 = quantile(unique(l),probs = rl[1]+rl[2]*(1-rl[1]))
    seg.2.lm.fix(l1 = l1,l2=l2,l = l,dx = dx)$aic
  }
  best.break = optim(par = c(0.5,0.5),fn = ll,method = "L-BFGS-B",lower = c(0.025,0.025),upper = c(0.975,0.975),l=l,dx=dx)
  l1=quantile(unique(l),probs = best.break$par[1])
  l2=quantile(unique(l),probs = best.break$par[1]+best.break$par[2]*(1-best.break$par[1]))
  best.lm = seg.2.lm.fix(l = l,dx = dx,l1 = l1,l2 = l2)
  return(list(l1 = l1,l2 = l2,aic = best.break$value,model=best.lm$model))
}
```
