## Supplementary material for "Microbial genomic trait evolution is dominated by frequent and rare pulsed evolution": simulate_with_track.R

```
#
source("fitPE_functions.R")
#
arg = commandArgs(TRUE)
#
params = list(BM = c(lambda1=0,size1=0,lambda2=0,size2=0,sigma=5000,epsilon=1),
              PE1 = c(lambda1=5,size1=1000,lambda2=0,size2=0,sigma=0,epsilon=1),
              PEBM1 = c(lambda1=5,size1=900,lambda2=0,size2=0,sigma=500,epsilon=1),
              PE2 = c(lambda1=5,size1=900,lambda2=100,size2=5,sigma=0,epsilon=1),
              PEBM2 = c(lambda1=5,size1=800,lambda2=100,size2=5,sigma=500,epsilon=1),
              PE3 = c(lambda1=0.2,size1=10000,lambda2=5,size2=400,lambda3=100,size3=10,sigma=0,epsilon=1),
              VRG = c(prob=0.5,k=2,theta=2500,epsilon=1))
models = c("BM","PE1","PEBM1","PE2","PEBM2","PE3","VRG")
tree = read.tree(arg[1])
#
for(i in 1:20){
  for(mm in models){
    if(mm=="PE3"){
      sim.truth = do.call("simulatePET3.phy.withTrack",c(list(phy=tree),as.list(params[[mm]])))
    }else{
      if(mm=="VRG"){
        sim.truth = do.call("simulateGamma.phy",c(list(tree=tree),as.list(params[[mm]])))
      }else{
        sim.truth = do.call("simulatePET2.phy.withTrack",c(list(phy=tree),as.list(params[[mm]])))
      }
    }
    saveRDS(sim.truth,sprintf("Sim_%s_%d.RDS",mm,i))
    saveRDS(list(phy=tree,dat=sim.truth$observed.trait[tree$tip.label]),sprintf("Sim_%s_%d.data.RDS",mm,i))
    if(mm=="PE2") write.table(sim.truth$observed.trait[tree$tip.label],
                              file = sprintf("Sim_%s_%d.txt",mm,i),quote = FALSE,
                              sep = "\t",row.names = TRUE,col.names = FALSE)
  }
}
```
